# Neuronal temperature perception induces specific defenses that enable C. elegans to cope with the enhanced reactivity of hydrogen peroxide at high temperature

**DOI:** 10.1101/2022.03.21.485202

**Authors:** Francesco A. Servello, Rute Fernandes, Matthias Eder, Nathan Harris, Olivier M. F. Martin, Natasha Oswal, Anders Lindberg, Nohelly Derosiers, Piali Sengupta, Nicholas Stroustrup, Javier Apfeld

## Abstract

Hydrogen peroxide is the most common reactive chemical that organisms face on the microbial battlefield. The rate with which hydrogen peroxide damages biomolecules required for life increases with temperature, yet little is known about how organisms cope with this temperature-dependent threat. Here, we show that *Caenorhabditis elegans* nematodes use temperature information perceived by sensory neurons to cope with the temperature-dependent threat of hydrogen peroxide produced by the pathogenic bacterium *Enterococcus faecium*. These nematodes preemptively induce the expression of specific hydrogen peroxide defenses in response to perception of high temperature by a pair of sensory neurons. These neurons communicate temperature information to target tissues expressing those defenses via an insulin/IGF1 hormone. This strategy, which we call “enhancer sensing,” is the first example of a multicellular organism inducing their defenses to a chemical when they sense an inherent enhancer of the reactivity of that chemical.

## Introduction

Reactive chemicals in the environment pose a lethal threat to organisms by changing the chemical composition of their macromolecules. But organisms are not passive chemical substrates, they have sophisticated defense mechanisms that deal with the threat posed by those chemicals. That threat is inherently temperature-dependent because chemical reactions occur at faster rates at higher temperatures (Arrhenius, 1889; Evans and Polanyi, 1935; Eyring, 1935). However, the extent to which the defense mechanisms protecting the organism from reactive chemicals are adjusted to balance the temperature-dependent threat posed by those chemicals remains poorly understood. In the present study, we used the nematode *C. elegans* as a model system to explore the extent to which temperature regulates how multicellular organisms deal with the threat of hydrogen peroxide.

Hydrogen peroxide (H_2_O_2_) is the most common reactive chemical that organisms face on the microbial battlefield (Mishra and Imlay, 2012). Bacteria, fungi, plants, and animal cells have long been known to use H_2_O_2_ as an offensive weapon that damages the nucleic acids, proteins, and lipids of their targets (Avery and Morgan, 1924; Imlay, 2018). *C. elegans* encounter a wide variety of bacteria in their ecological setting (Samuel et al., 2016; Schiffer et al., 2021), including many genera known to produce H_2_O_2_ (Passardi et al., 2007). H_2_O_2_ produced by a bacterium from the *C. elegans* microbiome, *Neorhizobium sp.,* causes DNA damage to the nematodes (Kniazeva and Ruvkun, 2019) and many bacteria—including *Enterococcus faecium, Streptococcus pyogenes*, *Streptococcus pneumoniae*, and *Streptococcus oralis*—kill *C. elegans* by producing millimolar concentrations of hydrogen peroxide (Bolm et al., 2004; Jansen et al., 2002; Moy et al., 2004).

Prevention and repair of the damage that hydrogen peroxide inflicts on macromolecules are critical for cellular health and survival (Chance et al., 1979). To avoid damage from H_2_O_2_, *C. elegans* rely on conserved cellular defenses, including H_2_O_2_-degrading catalases (Chavez et al., 2007; Schiffer et al., 2020). However, inducing those defenses at inappropriate times can cause undesirable side effects, including developmental defects (Doonan et al., 2008; Kramer-Drauberg et al., 2020), because H_2_O_2_ modulates the function of proteins involved in a wide variety of cellular processes, including signal transduction and differentiation (Hourihan et al., 2016; Kramer-Drauberg *et al*., 2020; Meng et al., 2021; Veal et al., 2007). We recently found that ten classes of sensory neurons in the nematode’s brain manage the challenge of deciding when the nematode’s tissues induce H_2_O_2_ defenses (Schiffer *et al*., 2020). Sensory neurons might be able to integrate a wider variety of inputs than the individual tissues expressing those defenses could integrate, enabling a better assessment of the threat of hydrogen peroxide.

In their habitat, *C. elegans* face daily and seasonal variations in temperature, which can affect a wide variety of processes, including development, reproduction, and lifespan (Golden and Riddle, 1984; Klass, 1977). Temperature also affects the growth of bacteria (Barber, 1908; Rosso et al., 1993) and, therefore, likely affects the interactions of *C. elegans* with beneficial and pathogenic bacteria (Samuel *et al*., 2016; Zhang et al., 2017). While nematodes do not regulate their own body temperature, they adjust their behavior and physiology in response to the perception of temperature by sensory neurons, enabling them to seek temperatures conducive to survival, avoid noxious temperature ranges, and induce heat defenses (Goodman and Sengupta, 2019; Hedgecock and Russell, 1975; Prahlad et al., 2008). Nematodes perceive temperature, in part, via seven classes of sensory neurons (Beverly et al., 2011; Biron et al., 2008; Chatzigeorgiou et al., 2010; Kuhara et al., 2008; Liu et al., 2012; Mori and Ohshima, 1995; Schild et al., 2014). Previously, we found that four of those classes of neurons regulate *C. elegans* peroxide resistance (Schiffer *et al*., 2020), suggesting that nematodes might adjust their peroxide defenses in response to temperature perception.

Here, we show that the lethality of hydrogen peroxide to *C. elegans* increases with temperature. Nematodes partially compensate for this by preemptively inducing their hydrogen peroxide defenses at high temperature. The temperature-dependent regulation of peroxide defenses is directed by the AFD sensory neurons. At high temperature, the AFD neurons repress the expression of the INS-39 insulin/IGF1 hormone and thereby alleviate inhibition by insulin/IGF1 signaling of the nematodes’ peroxide defenses. The insulin/IGF1 effector DAF-16/FOXO functions in intestinal cells to determine the size of the gene-expression changes induced by the absence of signals from the AFD neurons. This adaptive response to temperature enables the nematodes to better cope with H_2_O_2_ produced by the pathogenic bacterium *Enterococcus faecium.* By coupling the induction of H_2_O_2_ defenses to the perception of high temperature—an inherent enhancer of the reactivity of H_2_O_2_—the nematodes are assessing faithfully the threat that H_2_O_2_ poses. We refer to this sensing strategy as “enhancer sensing.”

## Results

### *C. elegans* induces long-lasting peroxide defenses in response to high temperature

In their natural habitat, *C. elegans* nematodes encounter many threats that can shorten their lifespan. A major chemical threat that *C. elegans* face is hydrogen peroxide (H_2_O_2_). Bacteria can produce millimolar concentrations of H_2_O_2_ (Bolm *et al*., 2004; Jansen *et al*., 2002; Moy *et al*., 2004), shortening *C. elegans* lifespan over ten-fold (Chavez *et al*., 2007). A major physical threat that *C. elegans* face is high temperature. Even a small increase in temperature from 20°C to 25°C—within the range of ambient temperatures that *C. elegans* prefers in nature (Crombie et al., 2019)—shortens *C. elegans* lifespan from 19 days to 15 days (Klass, 1977; Lee and Kenyon, 2009; Stroustrup et al., 2016).

To investigate the extent to which cultivation temperature might influence *C. elegans* survival in the presence of environmental peroxides, we measured the peroxide resistance of nematodes cultured within the nematodes’ preferred temperature range (Crombie *et al*., 2019). We cultured the nematodes at either 20°C or 25°C until the second day of adulthood, and then determined their subsequent survival at those temperatures in the presence of a peroxide in their environment. We used tert-butyl hydroperoxide (tBuOOH) because this peroxide, unlike H_2_O_2_, is not degraded efficiently by *Escherichia coli*—the nematodes’ conventional food in the laboratory (Brenner, 1974). Nematodes grown at 20°C survived an average of 1.6 days at 20°C, while those grown and assayed at 25°C survived 30% shorter (Figure 1A). Therefore, *C. elegans* peroxide resistance is temperature-dependent.

**Figure 1.**
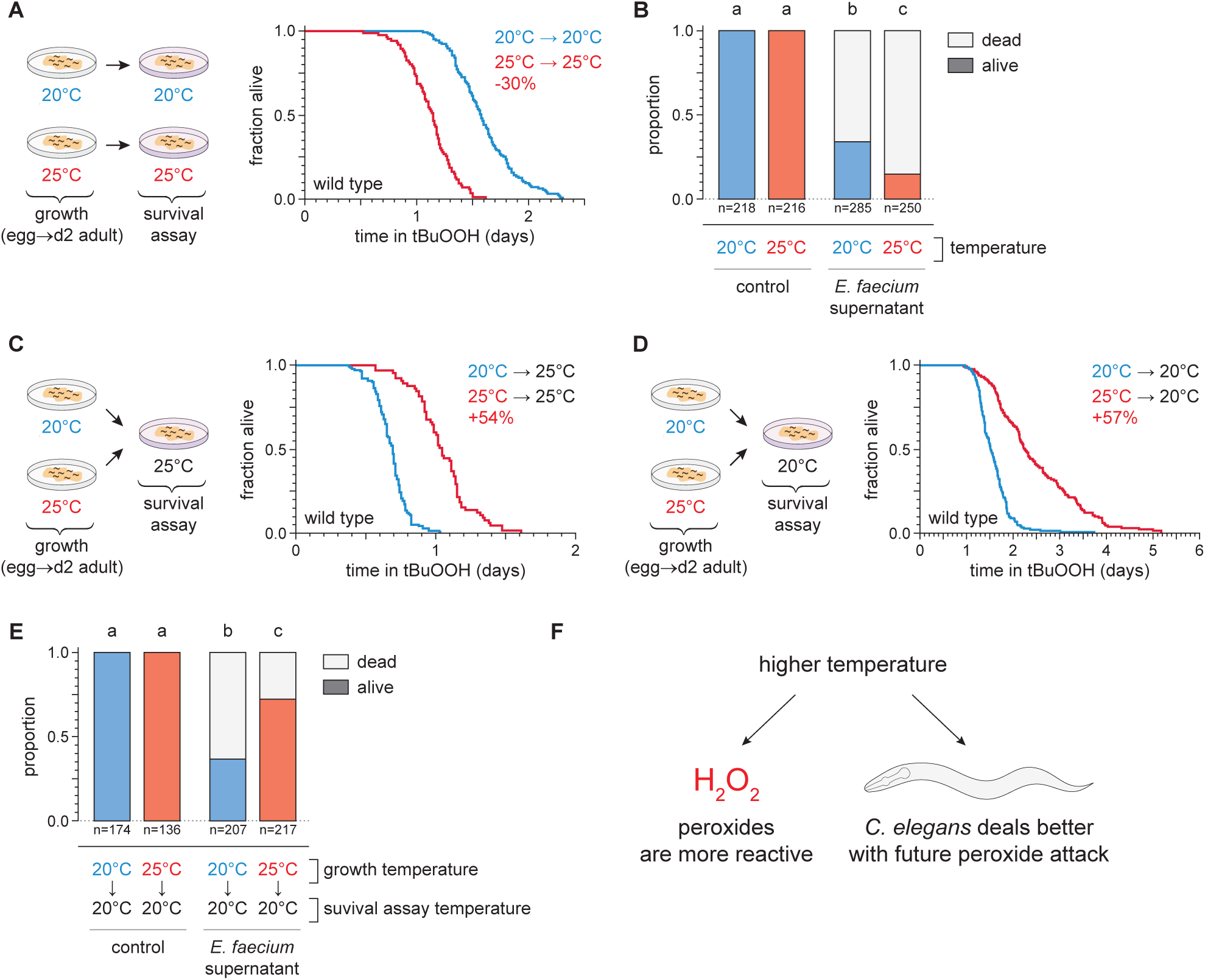
Temperature regulates the peroxide resistance of *C. elegans*. (A) Peroxide resistance of wild-type *C. elegans* grown and assayed at 20°C or 25°C. The fraction of nematodes remaining alive in the presence of 6 mM tert-butyl hydroperoxide (tBuOOH) is plotted against time. (B) Survival of wild-type *C. elegans* 16 hours after exposure to *E. faecium* E007 liquid-culture supernatant. Nematodes were grown and assayed at 20°C or 25°C. Groups labeled with different letters exhibited significant differences (*P* < 0.001, ordinal logistic regression) otherwise (*P* > 0.05). (C) Peroxide resistance at 25°C of wild-type *C. elegans* grown at 20°C or 25°C. (D) Peroxide resistance at 20°C of wild-type *C. elegans* grown at 20°C or 25°C. (E) Survival of wild-type *C. elegans* 16 hours after exposure to *E. faecium* E007 liquid-culture supernatant. Nematodes were grown at 20°C or 25°C and assayed at 20°C. Groups labeled with different letters exhibited significant differences (*P* < 0.001, ordinal logistic regression) otherwise (*P* > 0.05). (F) Peroxides killed *C. elegans* more quickly at 25°C than at 20°C, but nematodes grown at 25°C could better survive a subsequent peroxide exposure than those grown at 20°C. Statistical analyses for panels (A,C, and D) are in Supplementary Table 1.

We speculated that the nematodes’ survival to bacterially-produced H_2_O_2_ would, likewise, be shorter at 25°C than at 20°C. H_2_O_2_ produced by the pathogenic bacterium *Enterococcus faecium* is lethal to *C. elegans* (Chavez *et al*., 2007; Moy *et al*., 2004). We exposed day 2 adult nematodes that fed on *E. coli* JI377—a *katG katE ahpCF* triple null mutant strain which cannot degrade environmental H_2_O_2_ (Seaver and Imlay, 2001)—to the supernatant of an *E. faecium* liquid culture and, after 16 hours, determined the proportion of nematodes that survived. Compared to nematodes grown and assayed at 20°C, nematodes grown and assayed at 25°C were less likely to survive the *E. faecium* supernatant (Figure 1B), indicating that H_2_O_2_ was more lethal to *C. elegans* at the higher temperature. Together, these observations indicated that at the upper end of *C. elegans’* natural temperature range, multiple types of peroxides were more lethal to the nematodes.

We expected that increasing temperature would make peroxides more lethal to *C. elegans* because temperature increases the rate of chemical reactions, including those that mediate peroxide-dependent killing. If this were the only mechanism by which temperature affected *C. elegans’* peroxide resistance, then peroxide resistance should have been determined by the temperature the nematodes experienced during the peroxide-resistance assay and not by the temperature they experienced before they were exposed to peroxide. Alternatively, the temperature the nematodes experienced before encountering peroxides in the environment may have influenced the nematodes’ subsequent sensitivity to peroxide. For example, a high cultivation temperature may have irreversibly damaged the nematodes, thus rendering them more sensitive to peroxide-dependent killing.

To distinguish between these possibilities, we measured the effects of the nematodes’ growth-temperature history (before peroxide exposure) on their subsequent peroxide resistance by performing temperature-shift experiments where nematode populations grown at 20°C or 25°C were transferred to assay plates containing 6 mM tBuOOH at either 20°C or 25°C. To our surprise, we found that in survival assays performed at 25°C the nematodes grown at 25°C lived 54% longer than those grown at 20°C (Figure 1C). Similarly, in assays performed at 20°C, the nematodes grown at 25°C lived 57% longer than those grown at 20°C (Figure 1D). We also found that, compared with nematodes grown at 20°C, a higher proportion of nematodes grown at 25°C survived exposure to *E. faecium* liquid-culture supernatant at 20°C (Figure 1E). Therefore, nematodes grown at 25°C were more peroxide resistant than those grown at 20°C.

Our findings contradicted a model where temperature affected how quickly the nematodes were killed by peroxides only by influencing the reactivity of peroxides. In addition, those findings contradicted a prediction that high temperature would irreversibly render the nematodes more sensitive to peroxide-dependent killing. Instead, we conclude that even though peroxides killed *C. elegans* more quickly at 25°C than at 20°C, nematodes grown at 25°C could better survive a subsequent peroxide exposure than those grown at 20°C. These findings suggested that *C. elegans* nematodes induced their peroxide defenses when grown at the higher temperature to prepare for the increased lethal threat posed by peroxides at high temperature (Figure 1F).

To determine the extent to which these small differences in the nematodes’ growth temperature had lasting effects on their subsequent peroxide resistance, we repeated the temperature-shift experiments, but this time we transferred the nematodes to the higher or lower temperature one day before the peroxide survival assay (on day 1 of adulthood), and two days before (at the onset of adulthood). Shifting from 20°C to 25°C for two days was sufficient to improve peroxide survival at 25°C, but shifting only one day before the assay was not sufficient (Figure S1A). Therefore, nematodes grown at 20°C could increase their peroxide resistance in response to a temperature increase during adulthood. Nematodes down-shifted from 25°C to 20°C two days, one day, or immediately before the assay were all more peroxide resistant at 20°C than those grown continuously at 20°C (Figure S1B). Therefore, growth at 25°C could increase the nematodes’ peroxide resistance even days after they had been transferred to 20°C. Together, these observations suggested that *C. elegans* can induce long-lasting peroxide defenses in response to the higher cultivation temperature.

### AFD sensory neurons are required for the temperature-dependence of *C. elegans* peroxide resistance

We recently found that sensory neurons regulate *C. elegans* sensitivity to peroxides in the environment (Schiffer *et al*., 2020). To investigate whether temperature might regulate the nematode’s peroxide defenses via sensory neurons, we determined whether mutations that cause defects in the transduction of sensory information within neurons affected the extent to which temperature influenced the nematodes’ peroxide resistance. We examined *tax-4* cyclic GMP-gated channel mutants, which are defective in the transduction of several sensory stimuli, including temperature (Coburn and Bargmann, 1996; Komatsu et al., 1996). When grown at 20°C, *tax-4* mutants exhibited an over two-fold increase in peroxide resistance at 20°C relative to wild-type controls (Figures 2A,B). In contrast, when grown and assayed at 25°C, *tax-4* mutants exhibited a smaller increase in peroxide resistance, 49% (Figure 2C). Therefore, neuronal sensory transduction by TAX-4 channels normally lowers the nematodes’ peroxide resistance to a lesser extent at high cultivation temperature.

**Figure 2.**
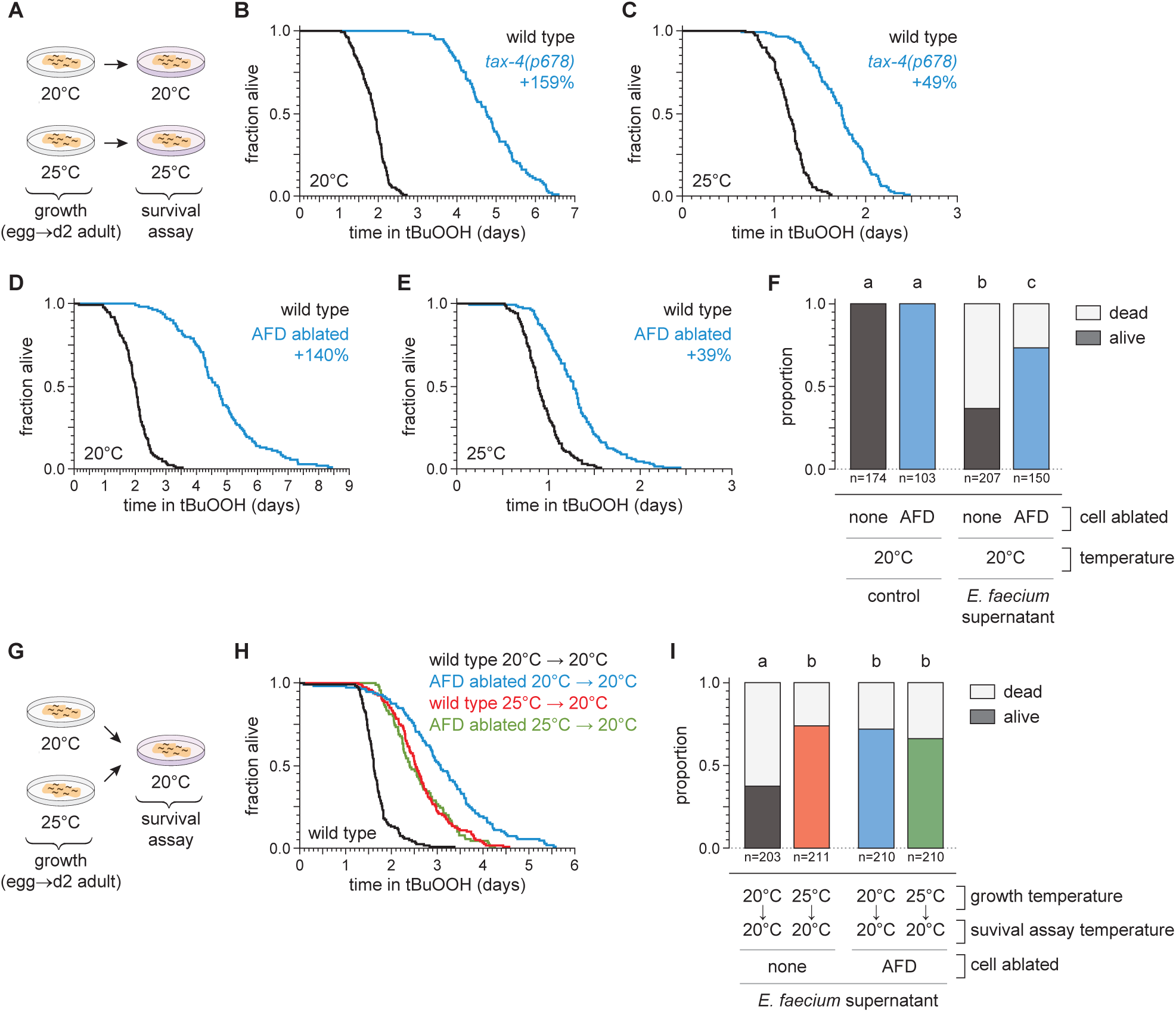
The AFD sensory neurons are required for the temperature-dependence of *C. elegans* peroxide resistance. (A) Diagram summarizing experimental strategy for panels (B-E). (B-C) The *tax-4(p678)* mutation increased peroxide resistance by a greater factor at 20°C (B) than at 25°C (C). (D-E) Genetic ablation of AFD increased peroxide resistance by a greater factor at 20°C (D) than at 25°C (E). (F) Genetic ablation of AFD increased the proportion of nematodes that survived 16 hours after exposure to *E. faecium* E007 liquid-culture supernatant. Nematodes were grown and assayed at 20°C. Groups labeled with different letters exhibited significant differences (*P* < 0.001, ordinal logistic regression) otherwise (*P* > 0.05). (G) Diagram summarizing experimental strategy for panels (H-I). (H) AFD-ablated nematodes grown at 25°C did not exhibit a further increase in peroxide resistance at 20°C, unlike wild-type (unablated) nematodes. (I) Growth at 25°C did not further increase the proportion of AFD-ablated nematodes that survived after 16 hours exposure to *E. faecium* E007 liquid-culture supernatant at 20°C, unlike in wild-type (unablated) nematodes. Groups labeled with different letters exhibited significant differences (*P* < 0.001, ordinal logistic regression) otherwise (*P* > 0.05). Statistical analyses for panels (B,C,D,E, and H) are in Supplementary Table 2.

To identify specific neurons that regulate the nematodes’ peroxide defenses in response to temperature, we focused on a single pair of neurons, the AFD neurons, chosen from the small subset of sensory neurons in which TAX-4 channels are expressed (Coburn and Bargmann, 1996; Komatsu *et al*., 1996). Previously, we found that genetic ablation of the AFD neurons via neuron-specific expression of caspase (Chelur and Chalfie, 2007; Glauser et al., 2011) increased *C. elegans* peroxide resistance (Schiffer *et al*., 2020). Because the AFD neurons respond to temperature via TAX-4 channels to regulate diverse temperature-dependent behaviors (Hedgecock and Russell, 1975; Mori and Ohshima, 1995), we speculated that they might also regulate peroxide resistance in response to temperature. To determine whether AFD neurons lowered the nematodes’ peroxide resistance in a temperature-dependent manner, we measured the effects of AFD ablation at 20°C and 25°C. Compared with wild-type nematodes, when grown and assayed at 20°C, the AFD-ablated nematodes exhibited an over two-fold increase in resistance to tBuOOH (Figure 2D) and H_2_O_2_ (Figure S2A), and were more likely to survive exposure to *E. faecium* liquid-culture supernatant (Figure 2F). At 25°C, AFD ablation increased resistance to tBuOOH by 31% (Figure 2E), a much smaller amount than at 20°C. Therefore, the AFD temperature-sensing neurons normally lower the nematodes’ peroxide defenses in a temperature-dependent manner.

If the AFD neurons were blocking the induction of peroxide defenses, we hypothesized that ablation of both AFD neurons might result in induction of peroxide defenses at lower temperatures similar to those seen in unablated nematodes at the higher temperature. Therefore, we predicted that AFD-ablated nematodes grown at 25°C would not exhibit a further increase in peroxide resistance at 20°C, unlike wild-type nematodes. Consistent with that prediction, we found that AFD-ablated nematodes grown at 25°C exhibited the same levels of resistance as wild-type nematodes grown at 25°C (Figures 2G,H). AFD-ablated nematodes grown continuously at 20°C exhibited the highest levels of peroxide resistance (Figure 2H). Similarly, in assays at 20°C measuring nematode survival after exposure to a supernatant derived from a liquid culture of *E. faecium*, AFD-ablated nematodes grown at either 20°C or 25°C survived as well as wild-type nematodes grown at 25°C (Figure 2I). We propose that, in wild-type nematodes, the extent to which the AFD neurons lower peroxide defenses is reduced in response to higher temperature.

### Hydrogen peroxide defenses are induced by high cultivation temperature and by AFD ablation

To investigate whether the higher cultivation temperature and the ablation of the AFD sensory neurons increased the nematodes’ peroxide resistance through a common defense mechanism, we used mRNA sequencing (mRNA-seq) to compare the extent to which those interventions affected gene expression. Collecting mRNA from day 2 adults grown at 20°C and 25°C and from AFD-ablated and unablated (wild-type) nematodes grown at 20°C, we identified differentially expressed transcripts. Relative to nematodes grown at 20°C, those grown at 25°C had lower expression of 2,446 genes and higher expression of 809 genes, out of 18,039 genes detected (q value < 0.01) (Figures 3A and S3A). These changes in gene expression were consistent with previous studies comparing gene expression in nematodes grown at 20°C and 25°C (Gomez-Orte et al., 2018) (Figures S3C,D) and in nematodes shifted from 23°C to 17°C (Sugi et al., 2011) (Figures S3F,G). AFD ablation lowered the expression of 2,077 genes and increased the expression of 2,225 genes, out of 7,912 genes detected (q value < 0.01) (Figures 3B and S3B). Therefore, both the higher cultivation temperature and the ablation of the AFD sensory neurons induced broad changes in gene expression.

**Figure 3.**
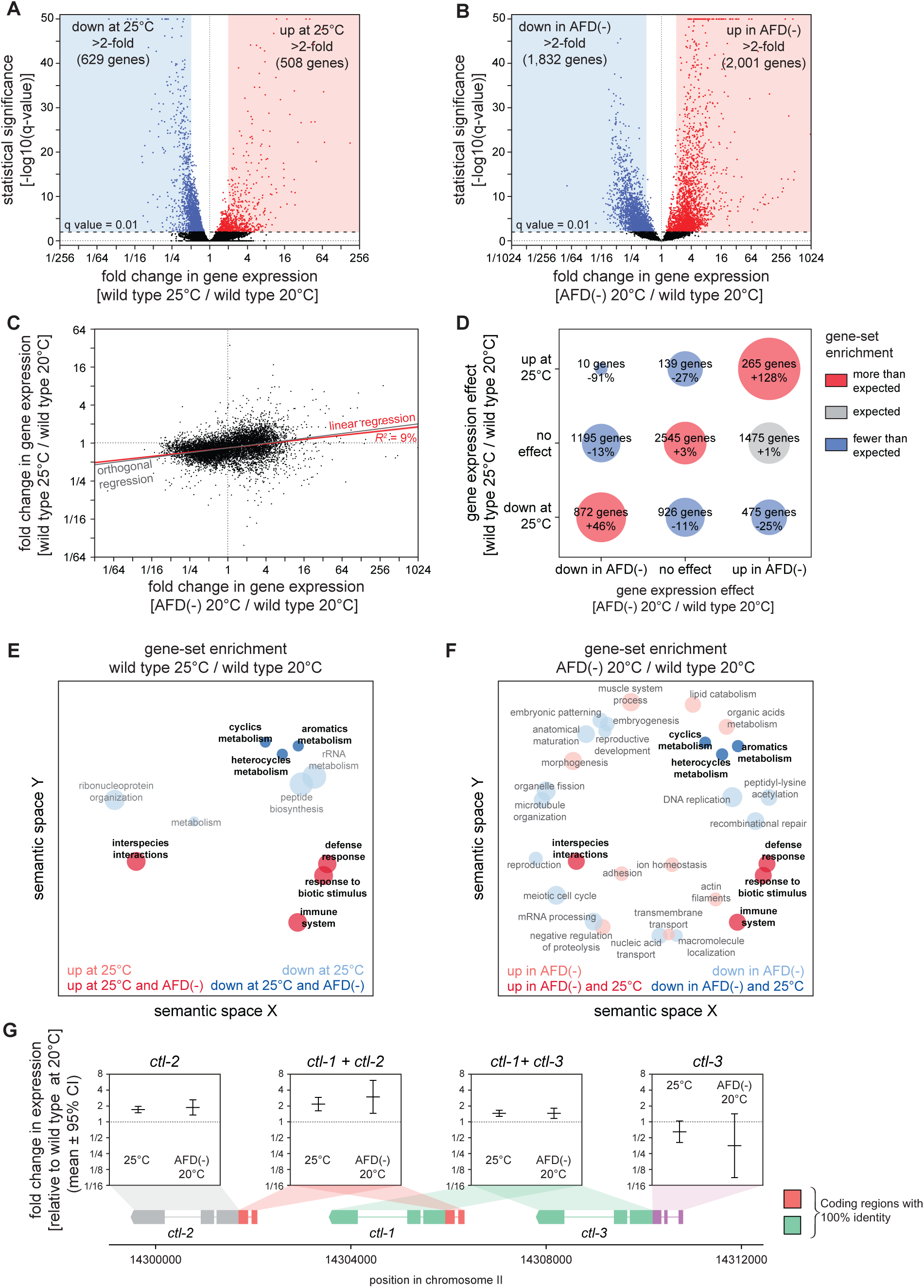
Hydrogen peroxide defenses are induced by high cultivation temperature and AFD ablation. (A-B) Volcano plots showing the level and statistical significance of changes in gene expression induced (A) in wild-type nematodes by growth at 25°C relative to growth at 20°C and (B) by AFD ablation in nematodes grown at 20°C relative to wild-type (unablated) nematodes grown at 20°C. Genes up- and down-regulated significantly (q value < 0.01) are shown in red and blue, respectively. (C) Growth at 25°C and AFD ablation at 20°C induced correlated changes in gene expression. Linear regression fit is shown as a red line flanked by a red area marking the 95% confidence interval of the fit. The orthogonal regression fit (grey line) makes no assumptions about the dependence or independence of the variables. (D) Co-regulation of genes up- and down-regulated significantly (q value < 0.01) by growth at 25°C and by AFD ablation at 20°C. Bubble size is proportional to gene-set enrichment (observed/expected). Gene-sets with significantly more or fewer genes than expected (*P* < 0.001, cell chi-square test) are colored red and blue, respectively; gene-sets of the expected size (*P* > 0.05) are colored grey. (E-F) Gene Ontology (GO) term enrichment analysis. (E) Biological processes associated with the set of 508 upregulated genes (red bubbles) and the set of 629 downregulated genes (blue bubbles) with a statistically significant and greater than two-fold change in expression in wild-type nematodes grown at 25°C relative to those grown at 20°C. (F) Biological processes associated with the set of 2,001 upregulated genes (red bubbles) and the set of 1,832 downregulated genes (blue bubbles) with a statistically significant and greater than two-fold change in expression in AFD-ablated nematodes grown at 20°C relative to wild-type (unablated) nematodes grown at 20°C. Bubble size is proportional to the statistical significance [-log_10_(*P* value)] of enrichment. Biological processes that were induced or repressed by both interventions are bolded and shaded with darker red and blue colors, respectively. (G) Average changes in expression and 95% confidence intervals induced by growth at 25°C and AFD ablation at 20°C within intervals in the genomic region encoding the three *C. elegans* catalase genes. Gene models show the positions and splicing pattern of each catalase gene, intervals with 100% nucleotide identity are shown in orange (*ctl-1* and *ctl-2*) and green (*ctl-1* and *ctl-3*), and unique intervals are show in grey (*ctl-2*) and purple (*ctl-3*).

Next, we asked whether higher cultivation temperature and ablation of AFD altered gene expression for each transcript by the same amount and in the same direction. We found that both interventions induced changes in gene expression that were linearly correlated in a positive manner (*R*^2^ = 9%, *P* < 0.0001, Figure 3C). We then asked whether this weak correlation was due to co-induction of upregulated genes, co-repression of down-regulated genes, or both, using categorical analysis. We found that genes with either higher or lower expression in both wild-type nematodes at 25°C and AFD-ablated nematodes at 20°C were disproportionally enriched, and that almost no genes were upregulated at 25°C but downregulated in AFD-ablated nematodes at 20°C (Figure 3D). Therefore, we conclude that cultivation temperature and AFD ablation induced overlapping changes in gene expression.

We then determined whether genes previously shown to be regulated between various temperature ranges were co-regulated by growth at 25°C and by ablation of AFD at 20°C. Genes expressed at a higher level at 25°C than at 20°C (Gomez-Orte *et al*., 2018) were upregulated by ablation of AFD at 20°C and were also, as expected, upregulated by growth at 25°C (Figure S3C); however, genes expressed at a higher level at 15°C than at 20°C (Gomez-Orte *et al*., 2018), were downregulated by growth at 25°C but were upregulated by ablation of AFD at 20°C (Figures S3D). In addition, genes induced more than two-fold when nematodes at 25°C were heat shocked by shifting them to 30°C (McCarroll et al., 2004) were upregulated by ablation of AFD at 20°C, but were unaffected by growth at 25°C (Figures S3E). We conclude that, in nematodes cultivated at 20°C, the AFD sensory neurons not only repress genes induced at a higher cultivation temperature (25°C), but also repress genes induced at a lower cultivation temperature (15°C) and in response to heat shock (30°C).

To identify processes that may be influenced by the transcriptomic changes induced by the higher cultivation temperature and by the ablation of the AFD neurons, we used Gene Ontology (GO) term enrichment analysis (Angeles-Albores et al., 2016; Ashburner et al., 2000) and clustered enriched GO terms based on semantic similarity (Supek et al., 2011), focusing on genes with more than a two-fold increase or decrease in expression between wild-type nematodes at 25°C and 20°C and between AFD-ablated and unablated nematodes at 20°C. We found that both higher temperature and AFD ablation downregulated genes associated with reproduction and expression in the germline and upregulated genes associated with defense and immune responses and expression in the intestine (Figures 3E,F and Table S4). To expand this analysis, we determined the extent to which higher cultivation temperature and ablation of AFD co-regulated the expression of gene sets affecting similar biological processes. We assigned each gene to a set of nested categories based on their physiological function and then their molecular function or cellular location using WormCat annotations (Higgins et al., 2021; Holdorf et al., 2020). Higher temperature and AFD ablation induced positively correlated changes in the average gene expression level of those gene-sets (*R*^2^ = 24%, *P* < 0.0001, Figure S4A). Therefore, the higher cultivation temperature and ablation of the AFD sensory neurons appeared to induce consistent changes in the expression of genes affecting similar biological processes.

We next determined whether genes induced when nematodes were exposed to tert-butyl hydroperoxide (Oliveira et al., 2009) were also induced by the higher cultivation temperature and by the ablation of the AFD neurons. Both growth at 25°C and ablation of AFD at 20°C increased the expression of those genes (Figure S4B). Therefore, in the absence of peroxide exposure, genes induced by peroxides were pre-induced in both nematodes cultivated at the higher temperature and in AFD-ablated nematodes, suggesting those nematodes were better prepared to deal with peroxides in the environment.

To identify specific peroxide defenses induced by the higher cultivation temperature and by the ablation of the AFD neurons, we focused on the catalase genes, which encode enzymes that degrade hydrogen peroxide (Loew, 1901; Nicholls, 2012; Togo et al., 2000). The *C. elegans* genome contains three catalase genes in tandem—two-newly duplicated cytosolic catalases, *ctl-1* and *ctl-3*, and a peroxisomal catalase, *ctl-2* (Petriv and Rachubinski, 2004)—that when overexpressed can more than double the nematode’s hydrogen peroxide resistance (Schiffer *et al*., 2020). *ctl-1* and *ctl-2* can function to increase *C. elegans* resistance to H_2_O_2_-dependent killing (Chavez *et al*., 2007; Schiffer *et al*., 2020). In our mRNA-seq analysis, we inferred that wild-type nematodes grown at 25°C had 46% higher levels of *ctl-1* expression and 73% higher levels of *ctl-2* expression compared to nematodes grown at 20°C (Figure 3G), and that ablation of the AFD neurons increased *ctl-1* expression by 46% and increased *ctl-2* expression by 89% (Figure 3G). Therefore, the cultivation temperature and the AFD neurons regulated the expression of hydrogen peroxide defenses.

Last, we determined the extent to which the higher cultivation temperature and the ablation of the AFD neurons affected the expression of genes induced by toxic organic compounds, toxic metals, and radiation (Eom et al., 2014; Greiss et al., 2008; Huffman et al., 2004; Lewis et al., 2009; Mueller et al., 2014; Sahu et al., 2013; Starnes et al., 2016). Growth at 25°C did not increase the expression of genes induced by acrylamide, formaldehyde, benzene, silver, cadmium, arsenic, UVB rays, X rays, and gamma rays (Figures S5A-I), but ablation of AFD at 20°C induced all of those gene sets (Figures S5A-I). Therefore, ablation of the AFD sensory neurons pre-induced genes induced by a wide variety of stressors, but the higher cultivation temperature only pre-induced a specific subset of genes that included hydrogen peroxide defenses and genes induced by peroxides.

### The high temperature-repressed INS-39 insulin/IGF1 hormone from the AFD sensory neurons lowers the nematode’s peroxide resistance

To investigate how the AFD sensory neurons regulated the nematode’s peroxide resistance, we explored whether they signaled to target tissues via insulin/IGF1 peptide hormones. Previous studies, including our own, showed that the insulin/IGF1 signaling pathway is a major determinant of peroxide resistance in *C. elegans* (Schiffer *et al*., 2020; Tullet et al., 2008). A recent single-neuron mRNA-seq study by the *C. elegans* Neuronal Gene Expression Map and Network (CeNGEN) consortium showed that AFD expresses many classes of peptide-hormone coding genes, including a subset of the 40 insulin/IGF1 genes in the genome (Taylor et al., 2019). We focused on the *ins-39* gene, which was highly expressed in AFD (Q. Ch’ng and J. Alcedo, personal communication) (Taylor *et al*., 2019).

To examine the expression of the *ins-39* gene, we used CRISPR/Cas9 genome editing to engineer a ‘‘transcriptional’’ reporter that preserved the 5’ and 3’ cis-acting regulatory elements of the *ins-39* gene (Tursun et al., 2009) by inserting into the *ins-39* gene locus a SL2-spliced intercistronic region fused to the coding sequence of the green fluorescent protein (GFP) (Figure 4A). The *ins-39* reporter was expressed exclusively in the AFD neurons (Figure 4B). The level of *ins-39* gene expression in AFD was higher in nematodes grown at 20°C than in those grown at 25°C (Figures 4C,D). Therefore, temperature regulated *ins-39* gene expression in the AFD sensory neurons. Temperature perception by the AFD neurons requires TAX-4 cyclic GMP-gated channels, as *tax-4* mutants do not exhibit changes in calcium dynamics or thermoreceptor currents in response to warming or cooling (Kimura et al., 2004; Ramot et al., 2008). The temperature-dependent expression of the *ins-39* gene in the AFD neurons required the function of *tax-4*, as a *tax-4* null mutation nearly abolished *ins-39* gene expression in nematodes grown at either 20°C or 25°C (Figures 4E and S6A). Taken together, these findings suggested that the AFD neurons lowered the expression of the INS-39 insulin/IGF1 hormone in response to TAX-4-dependent perception of the cultivation temperature.

**Figure 4.**
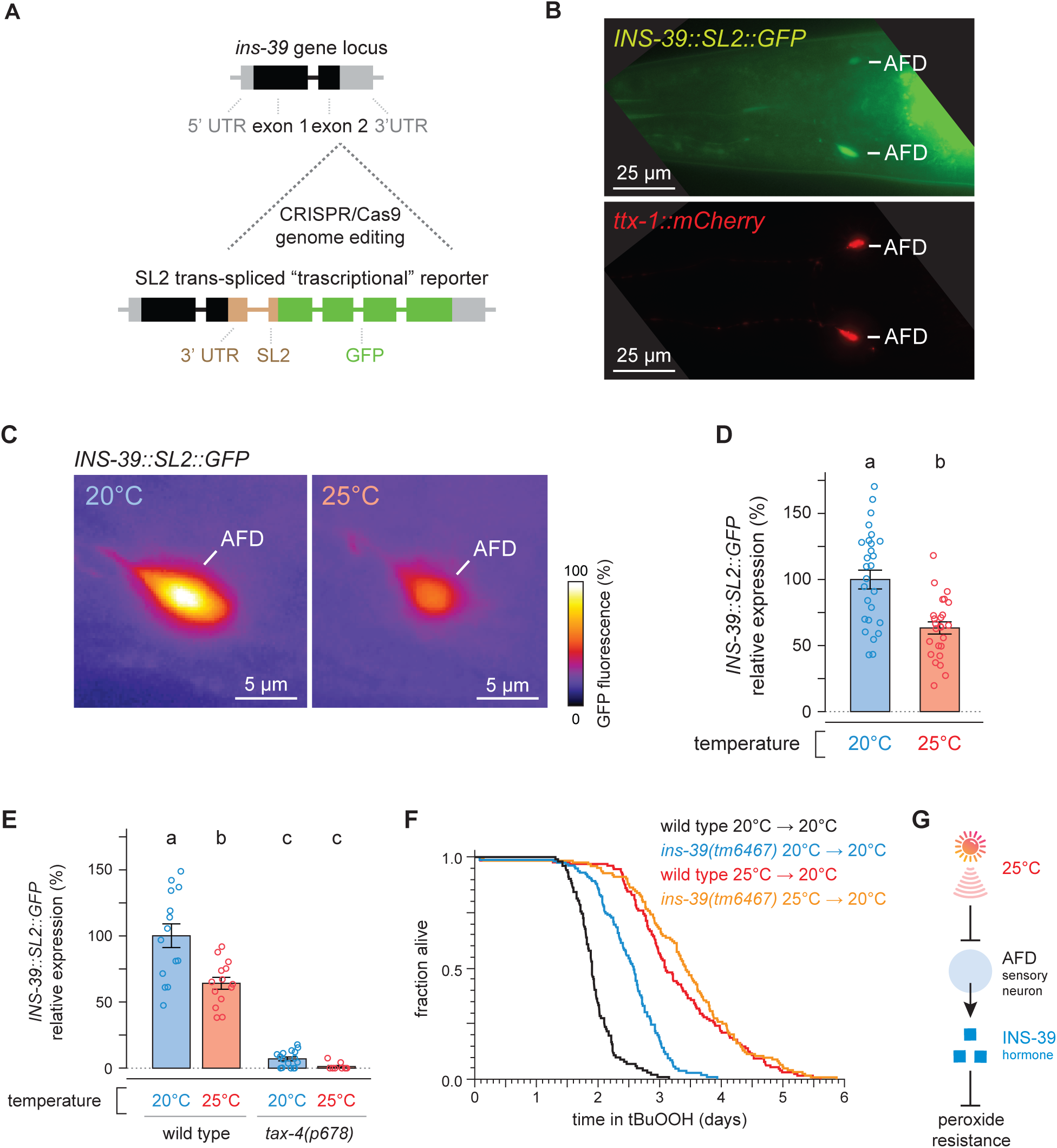
The high temperature-repressed INS-39 insulin/IGF1 hormone from the AFD sensory neurons lowers the nematode’s peroxide resistance. (A) Schematic of the CRISPR/Cas9 genome editing strategy used to engineer the *ins-39(oy167[ins-39::SL2::GFP])* ‘‘transcriptional’’ reporter. (B) Example animal co-expressing the *ins-39(oy167[ins-39::SL2::GFP])* reporter (top panel) and the AFD-specific reporter *Ex[ttx-1p::TagRFP]* (bottom panel). The head region is shown and only the AFD neurons are detected. Lines indicate the AFD soma. Scale bar = 25 µm. (C) Representative images of the expression of the *ins-39(oy167[ins-39::SL2::GFP])* reporter in nematodes grown at 20°C (left panel) and 25°C (right panel) in one of the bilateral AFD neurons. Scale bar = 5 µm. (D-E) Quantification of the expression of the *ins-39(oy167[ins-39::SL2::GFP])* reporter. (D) Reporter expression was lower in nematodes grown at 20°C than at 25°C. Groups labeled with different letters exhibited significant differences (*P* < 0.0001, ANOVA). (E) Reporter expression was nearly abolished in *tax-4(p678)* mutants. Groups labeled with different letters exhibited significant differences (*P* < 0.0001, Tukey HSD test) otherwise (*P* > 0.05). (F) The *ins-39(tm6467)* mutation increased peroxide resistance in nematodes grown and assayed at 20°C, but did not further increase peroxide resistance in nematodes grown at 25°C and assayed at 20°C. (G) Sensory perception of the cultivation temperature regulates the nematodes’ subsequent peroxide resistance. A high cultivation temperature lowers the expression of the AFD-specific INS-39 hormone, leading to the de-repression of the nematodes’ peroxide defenses. Statistical analysis for panel (F) is in Supplementary Table 5.

Next, we determined whether the INS-39 signal from AFD regulated the nematode’s peroxide resistance in response to cultivation temperature by examining the effects of a null mutation in *ins-39*. In nematodes grown and assayed at 20°C, the *ins-39(tm6467)* null mutation increased peroxide resistance by 29% relative to wild-type controls (Figure 4F). In contrast, *ins-39(tm6467)* did not further increase peroxide resistance in nematodes grown at 25°C and assayed at 20°C (Figure 4F). Therefore, INS-39 lowered the nematodes’ peroxide resistance in a manner dependent on the growth temperature history of the nematodes. We propose that at 20°C the AFD-specific hormone INS-39 represses the nematodes’ peroxide defenses (Figure 4G). At 25°C, however, the AFD neurons express lower levels of INS-39, leading to the de-repression of the nematodes’ peroxide defenses (Figure 4G).

Of note, we also determined whether the AFD neurons regulated the nematodes’ peroxide resistance through a process that required the neurotransmitter serotonin. Previous studies showed that serotonin is required for the induction of heat shock proteins in somatic tissues by AFD neurons in response to perception of a noxious 34°C heat shock (Prahlad *et al*., 2008; Tatum et al., 2015), a much higher temperature than the 20°C and 25°C cultivation temperatures we used in our studies. Serotonin biosynthesis requires the TPH-1 tryptophan hydroxylase (Shivers et al., 2009; Sze et al., 2000). We found that the peroxide resistance of AFD-ablated nematodes was unaffected by the *tph-1(n4622)* null mutation (Figure S6B). Therefore, the AFD neurons regulated peroxide resistance in a serotonin-independent manner.

### DAF-16/FOXO functions in the intestine to increase the nematode’s peroxide resistance in response to temperature-dependent signals from the AFD sensory neurons

To identify molecular determinants that might enable AFD to regulate the nematode’s peroxide defenses via INS-39 in response to temperature, we investigated whether the changes in gene expression induced by temperature and AFD ablation mimicked those induced by specific transcription factors in response to reduced insulin/IGF1 signaling. The FOXO transcription factor DAF-16 is essential for the increase in peroxide resistance and most other phenotypes of mutants with reduced signaling by the insulin/IGF1 receptor, DAF-2 (Kenyon et al., 1993; Lin et al., 1997; Ogg et al., 1997; Schiffer *et al*., 2020). Both higher temperature and AFD ablation increased the expression of genes directly upregulated by DAF-16 (Kumar et al., 2015) (Figures 5A,B) and increased the expression of genes upregulated in a *daf-16-*dependent manner in *daf-2(-)* mutants (Murphy et al., 2003) (Figure S7A,B). Genes directly upregulated by DAF-16 were disproportionately enriched among those upregulated significantly (q value < 0.01) by temperature and by AFD ablation, but not among those downregulated significantly by either intervention (Table S6). These findings were consistent with previous studies showing that the degree of nuclear localization of DAF-16 increases from 20°C to 25°C (Wolf et al., 2008). Together, these findings suggested that in response to cultivation temperature, reduced signaling by the AFD neurons might induce the nematodes’ peroxide defenses by increasing the activity of the DAF-16/FOXO transcription factor in target tissues.

**Figure 5.**
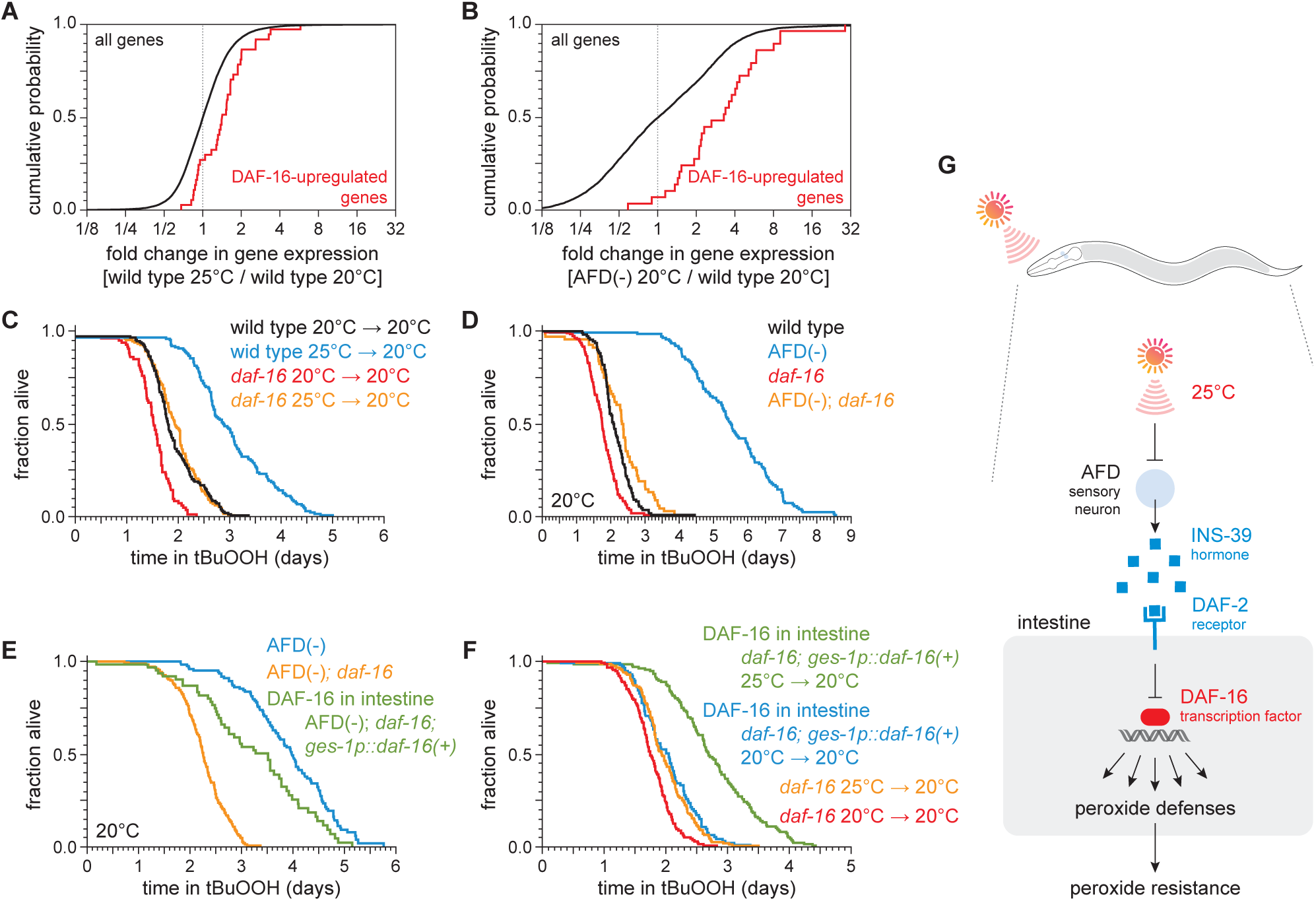
DAF-16/FOXO functions in the intestine to increase the nematode’s peroxide resistance in response to temperature-dependent signals from the AFD sensory neurons. (A-B) Genes directly upregulated by DAF-16 (Kumar *et al*., 2015) had higher expression (A) in nematodes grown at 25°C than in nematodes grown at 20°C and (B) in AFD-ablated nematodes grown at 20°C than in wild-type (unablated) nematodes grown at 20°C. (C-D) *daf-16(mu86)* suppressed most of the increased peroxide resistance of (C) nematodes grown at 25°C and assayed at 20°C and (D) AFD-ablated nematodes grown at 20°C. (E) Peroxide resistance of AFD-ablated nematodes expressing *daf-16(+)* only in the intestine, AFD-ablated *daf-16(mu86)* controls, and AFD-ablated nematodes for reference. Nematodes were grown and assayed at 20°C. (F) Peroxide resistance of transgenic nematodes expressing *daf-16(+)* only in the intestine, and *daf-16(mu86)* controls. Nematodes were grown at the indicated temperatures and assayed at 20°C. (G) The AFD sensory neurons repress the expression of H_2_O_2_-protection services in the nematodes’ intestine via insulin/IGF1 signaling. AFD expresses high levels of INS-39 at the lower cultivation temperature (20°C), leading to repression of the CTL-1 and CTL-2 H_2_O_2_-degrading catalases and of other peroxide defenses. At the higher cultivation temperature (25°C), AFD lowers INS-39 expression, de-repressing the DAF-16/FOXO factor that increases the expression of peroxide defenses in the intestine. Statistical analyses for panels (A-B) are in Supplementary Table 3 and statistical analyses for panels (C-F) are in Supplementary Table 7.

To determine whether DAF-16 was required for the regulation of peroxide resistance by the AFD sensory neurons and by cultivation temperature, we examined the effects of a null mutation in *daf-16*. The *daf-16(mu86)* null mutation decreased peroxide resistance in nematodes grown at 25°C and assayed at 20°C by 35%, a greater extent than in nematodes grown and assayed at 20°C (Figure 5C). Similarly, in nematodes grown and assayed at 20°C, the *daf-16(mu86)* null mutation decreased the peroxide resistance of AFD-ablated nematodes by 58%, but caused only a 18% reduction in peroxide resistance in unablated (wild-type) nematodes (Figure 5D). Therefore, the regulation of peroxide resistance by the AFD sensory neurons and by cultivation temperature was, in part, dependent on the DAF-16/FOXO transcription factor.

Next, we set out to identify which target tissues were important for increasing the nematode’s peroxide resistance via DAF-16 in response temperature-dependent signals from the AFD sensory neurons. First, we determined the extent to which restoring *daf-16(+)* expression in a specific tissue, using a tissue-specific promoter, increased peroxide resistance in AFD-ablated *daf-16* mutants. We speculated that *daf-16* might function in the intestine, because our transcriptomic analysis showed that both higher temperature and AFD ablation upregulated gene expression in the intestine (Table S4). Consistent with that prediction, in AFD-ablated *daf-16(mu86)* mutants grown and assayed at 20°C, restoring *daf-16(+)* expression only in the intestine was sufficient to partially rescue peroxide resistance to a level almost comparable to that of AFD-ablated wild-type nematodes (Figure 5E). Therefore, *daf-16(+)* functioned in the intestine to increase peroxide resistance in AFD-ablated nematodes.

We followed a similar scheme to determine whether intestinal DAF-16 increased the nematode’s peroxide resistance in response to cultivation temperature. In *daf-16(mu86)* mutants grown and assayed at 20°C, restoring *daf-16(+)* expression only in the intestine increased peroxide resistance by a small amount, 15% (Figure 5F), indicating that *daf-16(+)* function in the intestine was sufficient to increase peroxide resistance. Notably, in *daf-16(mu86)* mutants grown at 25°C and assayed at 20°C, restoring *daf-16(+)* expression only in the intestine increased peroxide resistance to a greater extent, 39%, than in nematodes grown and assayed at 20°C (Figure 5F). Therefore, temperature regulated the size of the increase in peroxide resistance induced by *daf-16(+)* function in the intestine.

Based on these observations, we propose that communication between AFD sensory neurons and the intestine via insulin/IGF1 signaling enables the nematode to regulate their peroxide defenses in response to perception of the cultivation temperature (Figure 5G). At a higher cultivation temperature, lower INS-39 expression by AFD leads to a decrease in signaling by the DAF-2 receptor, which enables DAF-16/FOXO transcription factors to induce peroxide defenses.

### SKN-1/NRF and DAF-16/FOXO collaborate to increase the nematodes’ peroxide resistance in response to signals from the AFD sensory neurons

We next examined whether other transcription factors might act with DAF-16 to increase peroxide resistance in AFD-ablated nematodes at 20°C. The DAF-3/coSMAD transcription factor (Patterson et al., 1997) is required for almost all of the increase in peroxide resistance induced by lack of DAF-7/TGFβ signaling from the ASI sensory neurons (Schiffer *et al*., 2020). In contrast, the *daf-3(mgDf90)* null mutation did not affect the peroxide resistance of AFD-ablated nematodes (Figure S7C). Therefore, unlike the ASI neurons, the AFD neurons did not regulate the nematodes’ peroxide resistance via DAF-3/coSMAD.

Like DAF-16, the NRF orthologue SKN-1 increases the nematodes’ peroxide resistance in response to reduced DAF-2 signaling (Tullet *et al*., 2008). The expression of genes upregulated by *skn-1(+)* in wild type nematodes (Oliveira *et al*., 2009) and in *daf-2* loss-of-function mutants (Ewald et al., 2015) was increased by AFD ablation but was not increased by higher cultivation temperature (Figure 6A and S7D-F), suggesting that SKN-1 might increase peroxide resistance in AFD-ablated nematodes. Knockdown of *skn-1* via RNA interference (RNAi) decreased the peroxide resistance of AFD-ablated nematodes by 58%, but caused a 27% reduction in peroxide resistance in wild-type nematodes (Figure 6B). RNAi of *skn-1* also decreased the peroxide resistance of AFD-ablated *daf-16* mutants (Figure 6C). In addition, RNAi of *skn-1* caused a larger reduction in peroxide resistance in *daf-16* mutants when the AFD neurons were ablated than when those neurons were present (Figure 6C), suggesting that DAF-16 and SKN-1 had non-overlapping roles in promoting peroxide resistance in AFD-ablated nematodes. We propose that when nematodes are cultured at 20°C, the AFD neurons promote signaling by the DAF-2/insulin/IGF1 receptor in target tissues, which subsequently lowers the nematode’s peroxide resistance by repressing transcriptional activation by SKN-1/NRF and DAF-16/FOXO.

**Figure 6.**
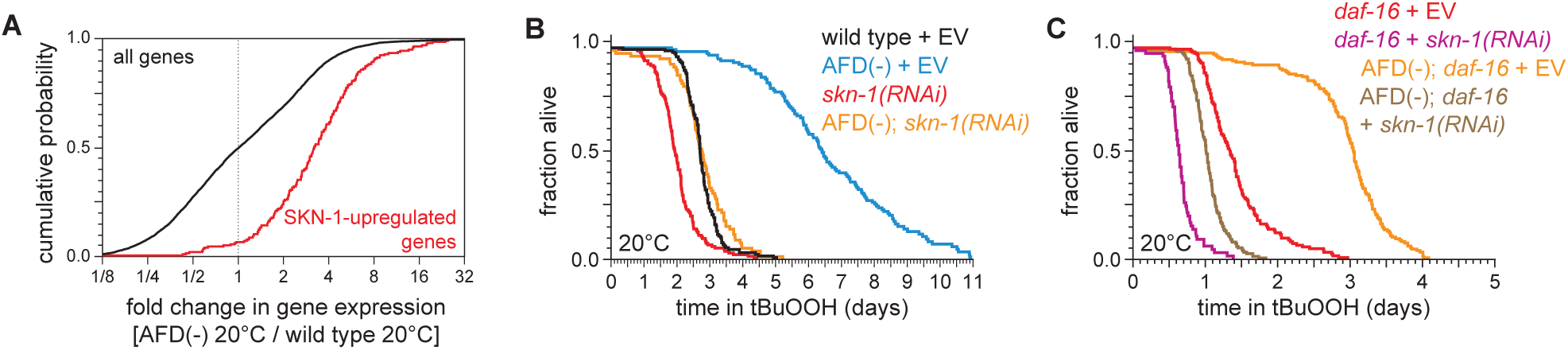
SKN-1/NRF and DAF-16/FOXO collaborate to increase the nematodes’ peroxide resistance in response to signals from the AFD sensory neurons. (A) Genes upregulated by *skn-1(+)* in wild type nematodes (Oliveira *et al*., 2009) had higher expression in AFD-ablated nematodes grown at 20°C than in wild-type (unablated) nematodes grown at 20°C. (B) *skn-1(RNAi)* suppressed most of the increased peroxide resistance of AFD-ablated nematodes grown at 20°C. Control RNAi consisted of feeding the nematodes the same bacteria but with the empty vector (EV) plasmid pL4440 instead of a plasmid targeting *skn-1*. (C) *skn-1(RNAi)* lowered peroxide resistance to a greater extent in AFD-ablated *daf-16(mu86)* mutants at 20°C than in (unablated) *daf-16(mu86)* mutants at 20°C. Statistical analysis for panel (A) is in Supplementary Table 3 and statistical analyses for panels (B-C) are in Supplementary Table 8.

### DAF-16/FOXO potentiates the changes in gene expression induced by the AFD sensory neurons

What role does the DAF-16/FOXO transcription factor play in regulating gene expression in response to signals from the AFD sensory neurons? In principle, DAF-16 could mediate all, some, or none of the changes in gene expression induced by AFD ablation. To distinguish between these possibilities, we used genome-wide epistasis analysis (Angeles-Albores et al., 2018) to compare the transcriptomes of unablated *daf-16(+)* [wild type] nematodes, unablated *daf-16(mu86)* mutants, AFD-ablated *daf-16(+)* nematodes, and AFD-ablated *daf-16(mu86)* mutants, on day 2 of adulthood and grown at 20°C. This analysis quantified the extent to which DAF-16 affected gene expression differently in AFD-ablated and unablated nematodes (Figures 7A and S8A).

**Figure 7.**
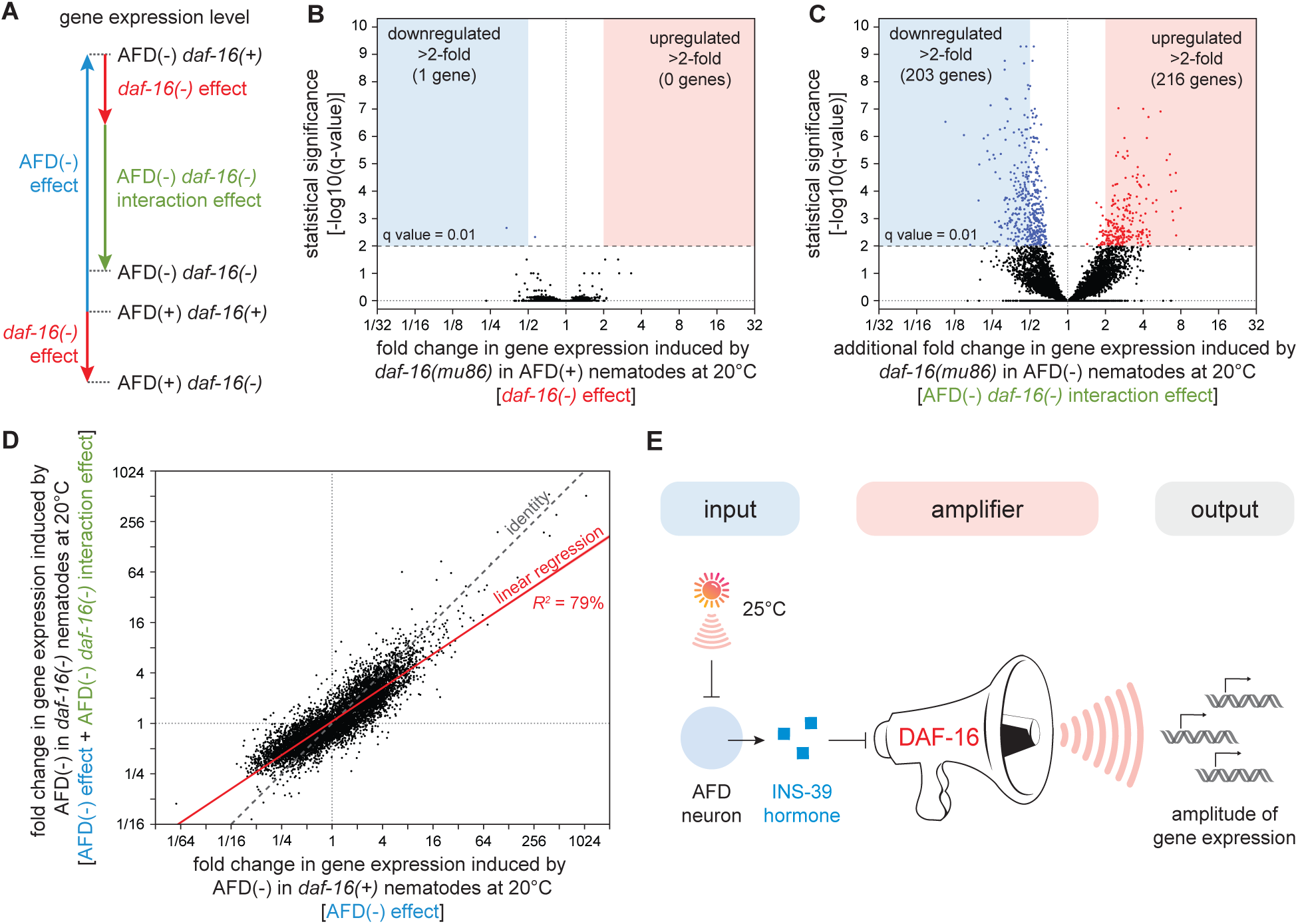
DAF-16/FOXO potentiates the changes in gene expression induced by the AFD sensory neurons. (A) We performed mRNA-seq on wild type [AFD(+) *daf-16(+)*], *daf-16(mu86)* null mutants [AFD(+) *daf-16(-)*], AFD-ablated nematodes [AFD(-) *daf-16(+)*], and AFD-ablated *daf-16(mu86)* null mutants [AFD(-) *daf-16(-)*] grown at 20°C, and used an epistasis model to quantify the extent to which AFD-ablation and *daf-16* mutation affected the expression of each gene, relative to wild type, in terms of the independent effects induced by AFD ablation (blue arrow) and by lack *daf-16* gene function (red arrow), and the additional effect induced by the interaction between AFD ablation and lack *daf-16* gene function (green arrow). (B-C) Volcano plots showing the level and statistical significance of (B) the changes in gene expression induced by lack *daf-16* gene function in unablated nematodes at 20°C and (C) the additional changes in gene expression induced by lack *daf-16* gene function in AFD-ablated nematodes at 20°C. Genes up- and down-regulated significantly (q value < 0.01) are shown in red and blue, respectively. (D) The effect of AFD ablation on gene expression at 20°C was systematically smaller in *daf-16(mu86)* mutants (y-axis) than in *daf-16(+)* nematodes (x-axis). Linear regression fit is shown as a red line flanked by a red area marking the 95% confidence interval of the fit. (E) The DAF-16/FOXO transcription factor amplifies the changes in gene expression induced by AFD ablation. This means that DAF-16 determines the gene-expression responsiveness, but not the response to temperature-dependent signals from the AFD sensory neurons.

Lack of *daf-16* gene function in unablated nematodes significantly lowered the expression of just 2 genes and significantly increased the expression of no genes, out of 7,387 genes detected (q value < 0.01) (Figure 7B). While, as expected, lack of *daf-16* gene function in unablated nematodes lowered the expression of genes directly upregulated by DAF-16 (Kumar *et al*., 2015) (Figure S8B,C) and lowered the expression of genes upregulated in a *daf-16-*dependent manner in *daf-2(-)* mutants (Murphy *et al*., 2003) (Figure S8D), these effects were small, averaging to just a 10% decrease in expression. Therefore, in nematodes grown at 20°C, DAF-16 was mostly unnecessary for regulating gene expression when the AFD neurons were present.

In contrast, when the AFD neurons were ablated, lack of *daf-16* gene function induced much broader changes in gene expression, lowering the expression of 431 genes and increasing the expression of 238 genes (q value < 0.01) (Figure 7C). In addition, in AFD ablated nematodes, lack of *daf-16* lowered the expression of genes upregulated in a *daf-16-*dependent manner in *daf-2(-)* mutants (Murphy *et al*., 2003) to a greater degree than in unablated nematodes (Figure S8E). Taken together, these findings showed that DAF-16 played a more consequential role in regulating gene expression when the AFD neurons were ablated.

Finally, we determined whether DAF-16 changed how the AFD neurons affected gene expression. The AFD neurons and DAF-16 did not independently regulate gene expression, because the extent to which DAF-16 affected gene expression deviated systematically from the level of gene expression predicted if AFD and DAF-16 acted independently (Figure S8F). To examine how the AFD neurons and DAF-16 jointly regulated gene expression, we compared the extent to which ablation of the AFD neurons affected gene expression in *daf-16(m86)* mutants and in *daf-16(+)* nematodes. The effect of AFD ablation on gene expression was systematically smaller in *daf-16(mu86)* mutants than in *daf-16(+)* nematodes (*R*^2^ = 79%, slope = 0.67, *P* < 0.0001, Figures 7D). Using simulations, we showed that effect was robust despite the uncertainty in our estimates of how much AFD and DAF-16 affected the expression of each gene (see Materials and Methods). In addition, we found that the extent to which AFD ablation affected the expression of sets of genes with related functions (Higgins *et al*., 2021; Holdorf *et al*., 2020) was systematically lower in *daf-16(mu86)* mutants than in *daf-16(+)* nematodes (*R*^2^ = 86%, slope = 0.67, *P* < 0.0001, Figure S8G). Therefore, the size of the effect of AFD ablation on gene expression was systematically smaller when the contribution of DAF-16 to gene expression was removed. We conclude that the DAF-16/FOXO transcription factor potentiates the changes in gene expression induced by ablation of the AFD sensory neurons (Figure 7E).

## Discussion

Across the tree of life, organisms face the lethal threat from hydrogen peroxide attack (Avery and Morgan, 1924; Imlay, 2018). This threat is inherently temperature dependent, because the reactivity of hydrogen peroxide increases with temperature (Evans and Polanyi, 1935; Eyring, 1935). In this study, we found that *C. elegans* nematodes use temperature information to deal with the lethal threat of hydrogen peroxide produced by the pathogenic bacterium *Enterococcus faecium*: when a pair of the nematodes’ neurons sensed a high cultivation temperature, they preemptively induced the nematodes’ hydrogen peroxide defenses. To our knowledge, the findings described here provide the first evidence of a multicellular organism inducing their defenses to a chemical when they sense an inherent enhancer of the reactivity of that chemical.

### Temperature perception by sensory neurons regulates *C. elegans* hydrogen peroxide defenses

We show here that a small increase in temperature—within the range that *C. elegans* nematodes prefer in nature (Crombie *et al*., 2019)—increases the nematodes’ sensitivity to killing by environmental peroxides and by hydrogen peroxide (H_2_O_2_) produced by the pathogenic bacterium *E. faecium*. These effects were not due to damage to the nematodes by the higher temperature but, instead, occurred despite the nematodes inducing protective defenses in response to experiencing the higher temperature before peroxide exposure.

We found that *C. elegans* deals with the enhanced threat posed by environmental peroxides at high cultivation temperature by coupling the induction of their H_2_O_2_ defenses to the perception of temperature by their AFD sensory neurons. These neurons have specialized sensory endings that are the primary thermoreceptors of the nematode, enabling them to adjust their behavior and heat defenses in response to temperature (Goodman and Sengupta, 2019; Hedgecock and Russell, 1975; Prahlad *et al*., 2008). The AFD sensory neurons used an INS-39 insulin/IGF1 hormone—which they expressed exclusively—to relay temperature information to the intestine, the tissue that provided H_2_O_2_-protection services to the nematode. At a low cultivation temperature, AFD expressed high levels of INS-39, leading to repression of the CTL-1 and CTL-2 catalases and of other peroxide-induced genes. However, at a high cultivation temperature AFD lowered INS-39 expression, leading to the induction of peroxide defenses by the DAF-16/FOXO transcriptional activator.

This repression of peroxide-protection services by AFD neurons at the lower cultivation temperature did not rely on the neurotransmitter serotonin, unlike the induction of heat defenses by AFD neurons in response to 34°C heat shock (Tatum *et al*., 2015). AFD ablation at 20°C also induced gene sets expressed at higher levels in response to low (15°C) cultivation temperature (Gomez-Orte *et al*., 2018) and gene sets induced by high heat (30°C) (McCarroll *et al*., 2004), but those gene sets were not induced by high cultivation temperature (25°C). Therefore, the AFD sensory neurons repressed gene-sets regulated by noxious heat, high cultivation temperature, and low temperature, and some of these gene-sets were regulated by distinct signals from AFD. In addition to expressing INS-39, these neurons express other peptide hormones—including hormones in the insulin/IGF1, FMRFamide, pigment dispersal factor, and oxytocin-vasopressin families (Barrios et al., 2012; Beets et al., 2012; Chen et al., 2016; Kim and Li, 2004; Taylor *et al*., 2019)—whose regulation could enable AFD to elicit specific responses to different temperature ranges. We conclude that the AFD thermosensory neurons play a central role in the regulation of distinct systemic responses to temperature.

### Target tissues control their responsiveness to sensory signals via DAF-16/FOXO

Using genome-wide epistasis analysis we showed that the DAF-16/FOXO transcription factor potentiated the changes in gene expression induced by AFD ablation. This means that DAF-16 determined the responsiveness, but not the response, of the intestine to temperature-dependent signals from AFD. We reason that while *C. elegans* intestinal cells manage the challenge of deciding when to induce their H_2_O_2_ defenses by relinquishing control of that decision to cultivation temperature-dependent signals from the AFD sensory neurons, they retain control of their responsiveness to those signals via the DAF-16/FOXO factor. It will be interesting to determine whether other processes regulating the function of intestinal DAF-16 determine the responsiveness of the induction of intestinal H_2_O_2_ defenses to cultivation temperature-dependent signals from the AFD neurons.

Multiple types of information appear to converge in a common mechanism to regulate the induction of intestinal peroxide defenses in *C. elegans*. Previously, we found that the SKN-1 and DAF-16 transcription factors collaborated to mediate the induction of peroxide defenses in response to information about food levels, sensed by the ASI neurons and communicated to the intestine via a TGFβ-Insulin/IGF1 hormone relay (Schiffer *et al*., 2020). DAF-16 and SKN-1 functions in the intestine are also regulated by signals from the germline (Berman and Kenyon, 2006; Ghazi et al., 2009; Steinbaugh et al., 2015), and TRPA-1 channels in the intestine regulate DAF-16 function in that tissue at 15°C (Xiao et al., 2013). In insects and mammals, insulin/IGF1 signaling components also regulate cellular antioxidant defenses (Brunet et al., 2004; Clancy et al., 2001; Holzenberger et al., 2003; Tatar et al., 2003). It will be interesting to explore the extent to which sensory-neuronal signals regulate hydrogen peroxide defenses via insulin/IGF1 signaling in all animals.

### Faithful threat assessment by sensing the enhancer of a threat

Why did *C. elegans* evolve a mechanism that couples the induction of hydrogen peroxide defenses to sensory perception of high cultivation temperature? One possibility is that temperature defenses might induce cross-protection from the stress caused by H_2_O_2_. However, high cultivation temperature increased the expression of the intestinal catalases CTL-1 and CTL-2, which are enzymes specialized for degrading H_2_O_2_ (Amara et al., 2001; Loew, 1901; Mishra and Imlay, 2012).

A second possibility is that the high cultivation temperature induced a broad set of defense responses that included those triggered by peroxides. However, that was not the case, because gene-sets induced by toxic metals, organic compounds, and high-energy radiation were not induced at 25°C. Therefore, the defenses induced by high cultivation temperature were specialized and included those for coping with the stress induced by hydrogen peroxide.

A third possibility is that *C. elegans* relies on temperature information to preemptively induce hydrogen peroxide defenses because when nematodes encounter a higher cultivation temperature in their natural habitat, they are more likely to subsequently encounter H_2_O_2_ and, therefore, need H_2_O_2_ protection. Such a sensing strategy, called adaptive prediction (Mitchell et al., 2009), is used by the bacteria *E. coli* and *Vibrio cholerae*, and by the yeasts *Saccharomyces cerevisiae* and *Candida albicans*, to sequentially induce specific defenses based on the typical order of stresses they encounter in their respective ecological settings (Mitchell *et al*., 2009; Rodaki et al., 2009; Schild et al., 2007; Tagkopoulos et al., 2008). Adaptive prediction would provide *C. elegans* with a good guess of when to induce their H_2_O_2_ defenses based on how often and how quickly high temperature is followed by H_2_O_2_ exposure in *C. elegans’* ecological setting (Levins, 1968; Mitchell and Pilpel, 2011).

However, adaptive prediction fails to capture how the reactivity of hydrogen peroxide is inherently enhanced by increasing temperature. Contrary to the expectation from adaptive prediction, *C. elegans* nematodes are not guessing that in their ecological setting increasing temperature leads to a higher H_2_O_2_ threat: in all ecological settings the nematodes’ proteins, nucleic acids, and lipids are inherently more likely to be damaged by H_2_O_2_ with increasing temperature because those chemical reactions necessarily run faster with increasing temperature (Arrhenius, 1889; Evans and Polanyi, 1935; Eyring, 1935). This means that by coupling the induction of H_2_O_2_ defenses to the perception of high temperature, the nematodes are assessing faithfully the threat that hydrogen peroxide poses. We refer to this sensing strategy as “enhancer sensing” (Figure 8).

**Figure 8.**
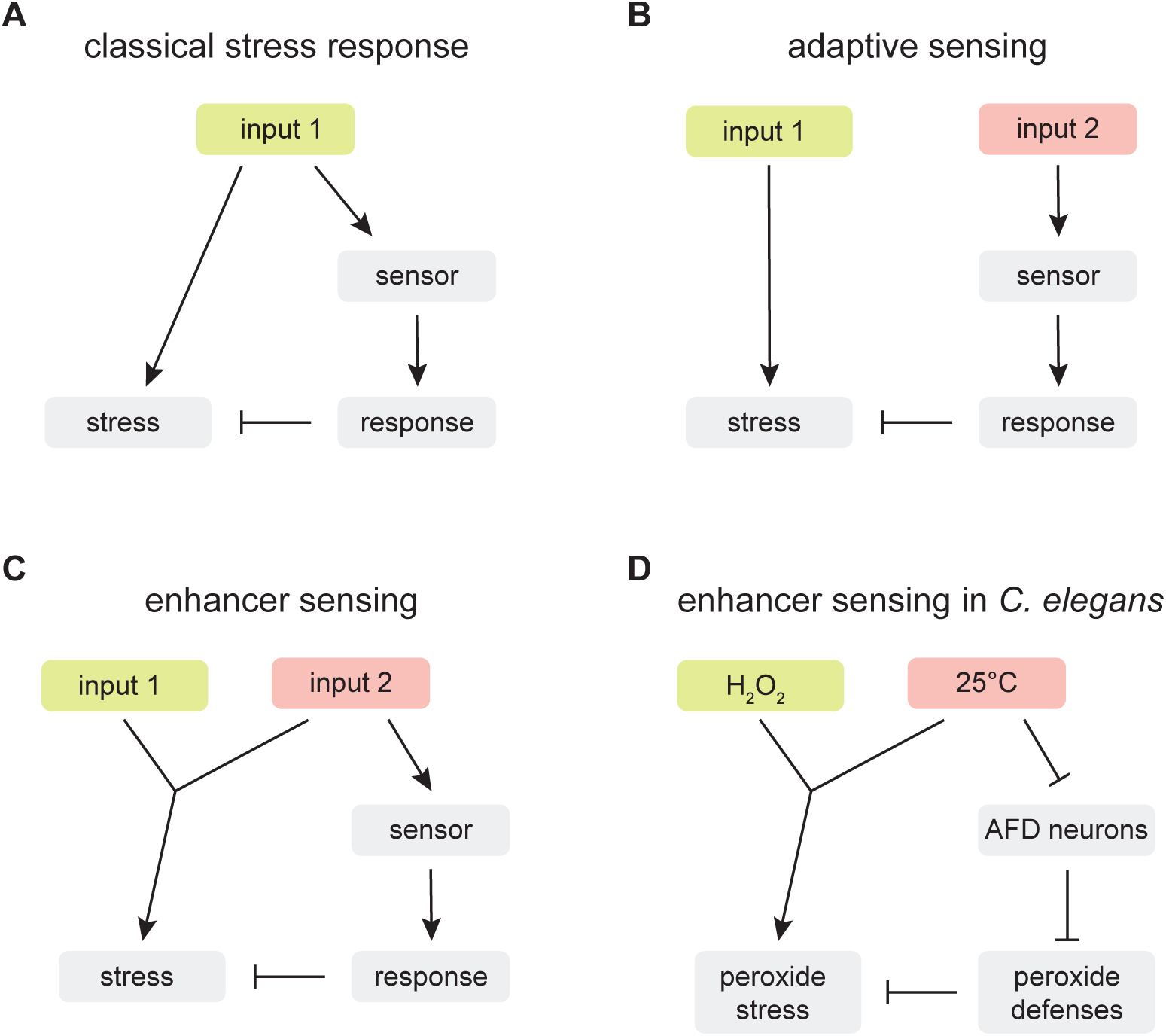
An enhancer sensing strategy enables *C. elegans* to assess faithfully the threat of hydrogen peroxide using temperature information. (A) Classical stress response: sensing is faithful to the threat the organism faces because the response that enables the organism to cope with the stress triggered by input 1 is coupled to the perception of input 1. (B) Adaptive sensing: sensing provides a good guess (informative based on the co-occurrence of inputs 1 and 2 in the ecological setting of the organism) but not necessarily faithful information, about the need for inducing a response, because input 2 does not trigger (nor affect the strength of an input that triggers) the stress that the organism copes with by inducing a response to input 2. (C) Enhancer sensing: faithful sensing occurs because the potency of the stress triggered by input 1 is enhanced input 2 and, therefore, perception of either input provides information about the need for responding to the threat posed by the interaction of those inputs. (D) The nematode *C. elegans* uses an enhancer sensing strategy that couples the de-repression of specific H_2_O_2_ defenses to the sensory perception of high temperature, an inherent enhancer of the reactivity of H_2_O_2_.

Enhancer sensing provides a framework for understanding the inherent adaptive value of coupling defense responses to the perception of inputs that enhance the need for those defenses. In classical stress responses, sensing is faithful to the threat the organism faces because the response that enables the organism to cope with the stress triggered by an input is coupled to the perception of that input (Figure 8A). Enhancer sensing is a strategy that realizes that faithful sensing also occurs when the potency of the stress triggered by an input is enhanced by another input and, therefore, perception of either input provides information about the need for responding to the threat posed by the interaction of those inputs (Figure 8C). In contrast, in adaptive sensing, the input does not trigger (nor affect the strength of an input that triggers) the stress that the organism copes with by inducing a response to that input; as a result, adaptive sensing provides a good guess informed by the ecological setting of the organism, but not necessarily faithful information, about the need for inducing a response (Figure 8B).

Previous studies have shown that high temperature induces H_2_O_2_ defenses in a wide variety of organisms, including bacteria (Engelmann et al., 1995; Mossialos et al., 2006), yeasts (Deegenaars and Watson, 1997; Mitchell *et al*., 2009; Wieser et al., 1991), plants (Hu et al., 2021; Nishizawa et al., 2006; Panchuk et al., 2002), cnidarians (Dash and Phillips, 2012), and human HeLa cells (Pallepati and Averill-Bates, 2010). Many of those examples could represent instances of enhancer sensing. More broadly, it will be interesting to determine the extent to which enhancer sensing strategies are used throughout the tree of life to couple specific defense responses to the perception of inputs that enhance the need for those defenses.

### *C. elegans* relies on a combination of sensing strategies to deal with the threat of hydrogen peroxide

*C. elegans* decides when to induce behavioral and cellular H_2_O_2_ defenses by relying on many classes of sensory neurons (Bhatla and Horvitz, 2015; Schiffer *et al*., 2020; Schiffer *et al*., 2021). These neurons are central to different types of sensing strategies that enable the nematode to deal with the lethal threat of environmental H_2_O_2_: enhancer sensing, adaptive anticipation, and classical stress sensing. The AFD sensory neurons provide enhancer sensing by coupling the induction of specific H_2_O_2_ defenses to temperature perception. The ASI sensory neurons provide adaptive anticipation by repressing H_2_O_2_ defenses in response to perception of *E. coli*, which protects the nematodes by depleting H_2_O_2_ from the nematode’s environment (Schiffer *et al*., 2020), like most bacteria in the nematodes’ ecological setting (Schiffer *et al*., 2021). The ASJ and I2 sensory neurons provide classical stress sensing by triggering locomotory escape and feeding inhibition, respectively, in response to perception of H_2_O_2_ (Bhatla and Horvitz, 2015; Schiffer *et al*., 2021). We speculate that by relying on a combination of sensing strategies, the nematodes can better manage the challenge of avoiding inducing costly H_2_O_2_ defenses that can cause undesirable side effects at inappropriate times.

## Materials and Methods

### *C. elegans* culture, strains, and transgenes

Wild-type *C. elegans* were Bristol N2. *C. elegans* were cultured at 20°C on NGM agar plates (Nematode Growth Medium, 17 g/L agar, 2.5 g/L Bacto Peptone, 3.0 g/L NaCl, 1 mM CaCl_2_, 1 mM MgSO_4_, 25 mM H_2_KPO_4_/HK_2_PO4 pH 6.0, 5 mg/L cholesterol) seeded with *E. coli* OP50, unless noted otherwise. Double mutant worms were generated by standard genetic methods. *ins-39(oy167[ins-39::SL2::GFP])* was made according to published protocols (Ghanta et al., 2021). Briefly, a dsDNA donor was made by amplifying SL2::GFP with 5’ SP9-modified oligos containing 35 bp overhangs homologous to the *ins-39* genomic locus for insertion immediately after the stop codon, using these primes: AGCAGGTCAAAGACGACTTCGTCACACTGCTCTGAGCTGTCTCATCCTACTTTCAC (forward) and ACTGGGCAAACGGAGAGTGAACGATGGAGCATTGACTATTTGTATAGTTCATCCATGCC (reverse). The genomic locus was cut with a crRNA targeting the sequence GATGGAGCATTGATCAGAGC. For a list of all bacterial and worm strains used in this study, see Tables S9 and S10, respectively. For a list of PCR genotyping primers and phenotypes used for strain construction, see Table S11.

### Survival assays

Automated survival assays were conducted using a *C. elegans* lifespan machine scanner cluster (Stroustrup et al., 2013) as described previously (Servello and Apfeld, 2020). This platform enables the acquisition of survival curves with very high temporal resolution and large population sizes. All chemicals were obtained from Sigma. For survival assays with 1 mM hydrogen peroxide and 6 mM tert-butyl hydroperoxide, the respective compound was added to molten agar immediately before pouring onto 50 mm NGM agar plates. Plates were dried (Stroustrup *et al*., 2013) and seeded with 100 µl of concentrated *E. coli* OP50 resuspended at an OD_600_ of 20 (Entchev et al., 2015). For RNAi experiments, the appropriate *E. coli* HT115 (DE3) strain was used instead. For hydrogen peroxide assays, *E. coli* JI377 was used instead (Seaver and Imlay, 2001). Nematodes were cultured at the specified developmental temperature until the onset of adulthood, and then cultured at the specified adult temperature, in groups of up to 100, on plates with 10 μg/ml 5-fluoro-2’-deoxyuridine (FUDR), to avoid vulval rupture (Leiser et al., 2016) and eliminate live progeny. Day 2 adults were transferred to lifespan machine assay plates. A typical experiment consisted of up to four genotypes or conditions, with 4 assay plates of each genotype or condition, each assay plate containing a maximum of 40 nematodes, and 16 assay plates housed in the same scanner. All experiments were repeated at least once, yielding the same results. Scanner temperature was calibrated to 20°C or 25°C with a thermocouple (ThermoWorks USB-REF) on the bottom of an empty assay plate. Death times were automatically detected by the lifespan machine’s image-analysis pipeline, with manual curation of each death time through visual inspection of all collected image data (Stroustrup *et al*., 2013), without knowledge of genotype or experimental condition.

### *Enterococcus faecium* survival assays

Nematodes were continuously fed *E. coli* JI377 for at least three generations and grown under the specified developmental temperature until the onset of adulthood, then cultivated at the specified adult temperature on plates with 10 μg/ml FUDR. *E. faecium* E007 was cultured in 500 mL of Brain Heart Infusion (BHI) medium overnight at 37°C without aeration and then aerated for four hours at 37°C prior to collecting the supernatant. The supernatant was stored at -20°C and incubated at the specified temperature before use. Nematodes were washed with M9 buffer with 0.01% Tween and transferred into 24 well plates containing either BHI or the appropriate amount of *E. faecium* E007 supernatant, *E. coli* JI377 at an OD_600_ of 2, in a final volume of 2 ml. Plates were sealed with parafilm, incubated at the specified temperature, and survival was scored after 16 hours by mixing the wells via pipette and scoring for movement using a dissection stereo microscope equipped with white-light transillumination.

### Transcriptomic analysis

mRNA for sequencing was extracted from day 2 adult animals. Worms were cultured on NGM agar plates seeded with *E. coli* OP50 and synchronized at the late L4 stage by transfer onto new NGM agar plates seeded with *E. coli* OP50 and supplemented with 10 μg/mL FUDR. Worms were cultured at 20°C except for the growth temperature assay for which worms were cultured at 25°C or 20°C for four generations before sampling at the respective temperatures. We adapted a nematode lysis protocol (Ly et al., 2015) for bulk lysis to pool 30 individuals per sample in 120 µL of lysis buffer. cDNA preparation from mRNA was performed by SmartSeq2 as described (Picelli et al., 2014). cDNA was purified using an in-house paramagnetic bead-based DNA purification system mimicking Agencourt AMPure XP magnetic beads. Dual-barcoded Nextera sequencing libraries were prepared according to the manufacturer’s protocol and purified twice with magnetic beads. Libraries were sequenced on an Illumina NextSeq 500 with a read length of 38 bases and approx. 2.0 x 10^6^ paired-end reads per sample. RNA-seq reads were aligned to the *C. elegans* Wormbase reference genome (release WS265) using STAR version 2.6.0c (Dobin et al., 2013) and quantified using featureCounts version 2.0.0 (Liao et al., 2014), both using default settings. To quantify the expression within intervals in the genomic region encoding the three *C. elegans* catalase genes, we created a GTF that matches genomic positions defined previously (Petriv and Rachubinski, 2004). The reads count matrix was normalized using scran (Lun et al., 2016). Differential analysis was performed using a negative binomial generalized linear model as implemented by DESeq2 (Love et al., 2014). A batch replicate term was added to the regression equation to control for confounding. Batch-corrected counts were obtained by matching the quantiles of distributions of counts to the batch-free distributions as in the Combat-seq method (Zhang et al., 2020). Principal component analysis was performed on the batch-corrected and normalized log counts with a pseudo-count of one. To access the significance of the slope between the sum of the “AFD(-) effect” and “AFD(-) *daf-16(-)* interaction effect” coefficients and the “AFD(-) effect” coefficient, we simulated 1000 sets of coefficients using a normal distribution with mean equal to the maximum-likelihood estimate of the coefficients and with standard deviation equal to the standard error of the estimates. We then fit a linear regression to each of the simulated coefficients and computed their coefficient of determination (*R^2^*). These simulations showed that, after accounting for the level of uncertainty on our estimates of the values of the coefficients for “AFD(-) effect” and “AFD(-) *daf-16(-)* interaction effect”, the average value of the regression’s slope was 0.6826 (99% confidence interval [0.6825, 0.6827]) and the average *R^2^* was 0.699 (99% confidence interval [0.6989, 0.6991]). Gene Ontology enrichments, tissue enrichment analysis, and phenotypic enrichment analysis were determined by using the WormBase Enrichment Suite (Angeles-Albores *et al*., 2016). We clustered and plotted GO terms with q-value *<* 10^-6^ using REVIGO (Supek *et al*., 2011). Curated gene expression data sets were obtained from WormExp (Yang et al., 2016). A curated hierarchical classification of genes into sets based on physiological function, molecular function, and cellular location was obtained from WormCat (Higgins *et al*., 2021; Holdorf *et al*., 2020).

### Microscopy

Nematodes carrying the *ins-39(oy167[ins-39::SL2::GFP])* allele were immobilized with 20 mM tetramisole, mounted on 10% agarose pads on slides, and imaged on a Zeiss Axio Imager M2 epifluorescent microscope with a 63x objective. Exposure time was set to 300 ms and images were acquired with 2×2 binning. Detectable GFP expression was determined to be entirely restricted to AFD, in day 1 adults, by co-expression of the AFD-specific marker *ttx-1p::mCherry* (Satterlee et al., 2001). Images for GFP quantification were acquired with no red marker in the background. Images were processed in ImageJ, and expression was quantified from a maximum projected z-stack as corrected total cell fluorescence (CTCF) by the equation CTCF = Integrated Density – (Area of selected cell ROI x Mean fluorescence of a nearby background ROI).

### RNA interference

*E. coli* HT115 (DE3) bacteria with plasmids expressing dsRNA targeting specific genes were obtained from the Ahringer and Vidal libraries (Kamath et al., 2001; Rual et al., 2004). Empty vector plasmid pL4440 was used as control. Bacterial cultures were grown in LB broth with 100 μg/ml ampicillin at 37°C and seeded onto NGM agar plates containing 50 μg/ml carbenicillin and 2 mM IPTG. Nematodes were cultivated on *E. coli* OP50 until day two of adulthood, then transferred to RNAi plates. Their progeny was subsequently used for each assay.

### Statistical analysis

Statistical analyses were performed in JMP Pro version 15 (SAS). Survival curves were calculated using the Kaplan-Meier method. We used ANOVA to determine whether the fold-change in gene expression of specific gene sets and of all genes were equal. We used ANOVA for GFP expression comparisons and, in cases where more than two groups were compared, used the Tukey HSD post-hoc test to determine which pairs of groups in the sample differed. We used the Cell chi-square test to determine if a cell in a table differed from its expected value in the overall table. We used ordinal linear regression to determine whether the proportions of dead animals after treatment with *E. faecium* supernatant were equal across groups and to quantify interactions between groups using the following linear model: data = Intercept + group 1 + group 2 + group 1 * group 2*+ ε*. The second to last term in this model quantifies the existence, magnitude, and type (synergistic or antagonistic) of interaction between groups. We used the Bonferroni correction to adjust *P* values when performing multiple comparisons.

## Acknowledgements

We thank Erin Cram, Jodie Schiffer, and Yuyan Xu for detailed comments on our manuscript. Joy Alcedo, Ryan Baugh, Danielle Garsin, James Imlay, and Yun Zhang kindly provided strains. Queelim Ch’ng and Joy Alcedo shared that *ins-39* is expressed in AFD. We benefitted from discussions with members of Javier Apfeld’s and Erin Cram’s labs. We derived some information from Wormbase, which is supported by the National Human Genome Research Institute at the NIH (grant #U41 HG002223), the UK Medical Research Council, and the UK Biotechnology and Biological Sciences Research Council. Some strains were provided by the CGC, which is funded by NIH Office of Research Infrastructure Programs (P40 OD010440).

## References

Amara, P., Andreoletti, P., Jouve, H.M., and Field, M.J. (2001). Ligand diffusion in the catalase from Proteus mirabilis: a molecular dynamics study. Protein Sci 10, 1927–1935. 10.1110/ps.14201.

Angeles-Albores, D., Puckett Robinson, C., Williams, B.A., Wold, B.J., and Sternberg, P.W. (2018). Reconstructing a metazoan genetic pathway with transcriptome-wide epistasis measurements. Proc Natl Acad Sci U S A 115, E2930–E2939. 10.1073/pnas.1712387115.

Angeles-Albores, D., RY, N.L., Chan, J., and Sternberg, P.W. (2016). Tissue enrichment analysis for C. elegans genomics. BMC Bioinformatics 17, 366. 10.1186/s12859-016-1229-9.

Arrhenius, S. (1889). Über die Reaktionsgeschwindigkeit bei der Inversion von Rohrzucker durch Säuren. Zeitschrift für physikalische Chemie 4, 226–248.

Ashburner, M., Ball, C.A., Blake, J.A., Botstein, D., Butler, H., Cherry, J.M., Davis, A.P., Dolinski, K., Dwight, S.S., Eppig, J.T., et al. (2000). Gene ontology: tool for the unification of biology. The Gene Ontology Consortium. Nature genetics 25, 25–29.

Avery, O.T., and Morgan, H.J. (1924). The Occurrence of Peroxide in Cultures of Pneumococcus. The Journal of experimental medicine 39, 275–287.

Barber, M.A. (1908). The rate of multiplication of Bacillus coli at different temperatures. The Journal of Infectious Diseases, 379–400.

Barrios, A., Ghosh, R., Fang, C., Emmons, S.W., and Barr, M.M. (2012). PDF-1 neuropeptide signaling modulates a neural circuit for mate-searching behavior in C. elegans. Nature neuroscience 15, 1675–1682. 10.1038/nn.3253.

Beets, I., Janssen, T., Meelkop, E., Temmerman, L., Suetens, N., Rademakers, S., Jansen, G., and Schoofs, L. (2012). Vasopressin/oxytocin-related signaling regulates gustatory associative learning in C. elegans. Science 338, 543–545. 10.1126/science.1226860.

Berman, J.R., and Kenyon, C. (2006). Germ-cell loss extends C. elegans life span through regulation of DAF-16 by kri-1 and lipophilic-hormone signaling. Cell 124, 1055–1068.

Beverly, M., Anbil, S., and Sengupta, P. (2011). Degeneracy and neuromodulation among thermosensory neurons contribute to robust thermosensory behaviors in Caenorhabditis elegans. The Journal of neuroscience : the official journal of the Society for Neuroscience 31, 11718–11727. 10.1523/JNEUROSCI.1098-11.2011.

Bhatla, N., and Horvitz, H.R. (2015). Light and hydrogen peroxide inhibit C. elegans Feeding through gustatory receptor orthologs and pharyngeal neurons. Neuron 85, 804–818. 10.1016/j.neuron.2014.12.061.

Biron, D., Wasserman, S., Thomas, J.H., Samuel, A.D., and Sengupta, P. (2008). An olfactory neuron responds stochastically to temperature and modulates Caenorhabditis elegans thermotactic behavior. Proc Natl Acad Sci U S A 105, 11002–11007. 10.1073/pnas.0805004105.

Bolm, M., Jansen, W.T., Schnabel, R., and Chhatwal, G.S. (2004). Hydrogen peroxide-mediated killing of Caenorhabditis elegans: a common feature of different streptococcal species. Infection and immunity 72, 1192–1194.

Brenner, S. (1974). The genetics of Caenorhabditis elegans. Genetics 77, 71–94.

Brunet, A., Sweeney, L.B., Sturgill, J.F., Chua, K.F., Greer, P.L., Lin, Y., Tran, H., Ross, S.E., Mostoslavsky, R., Cohen, H.Y., et al. (2004). Stress-dependent regulation of FOXO transcription factors by the SIRT1 deacetylase. Science 303, 2011–2015.

Chance, B., Sies, H., and Boveris, A. (1979). Hydroperoxide metabolism in mammalian organs. Physiological reviews 59, 527–605. 10.1152/physrev.1979.59.3.527.

Chatzigeorgiou, M., Yoo, S., Watson, J.D., Lee, W.H., Spencer, W.C., Kindt, K.S., Hwang, S.W., Miller, D.M., 3rd, Treinin, M., Driscoll, M., and Schafer, W.R. (2010). Specific roles for DEG/ENaC and TRP channels in touch and thermosensation in C. elegans nociceptors. Nature neuroscience 13, 861–868. 10.1038/nn.2581.

Chavez, V., Mohri-Shiomi, A., Maadani, A., Vega, L.A., and Garsin, D.A. (2007). Oxidative stress enzymes are required for DAF-16-mediated immunity due to generation of reactive oxygen species by Caenorhabditis elegans. Genetics 176, 1567–1577. 10.1534/genetics.107.072587.

Chelur, D.S., and Chalfie, M. (2007). Targeted cell killing by reconstituted caspases. Proc Natl Acad Sci U S A 104, 2283–2288. 10.1073/pnas.0610877104.

Chen, Y.C., Chen, H.J., Tseng, W.C., Hsu, J.M., Huang, T.T., Chen, C.H., and Pan, C.L. (2016). A C. elegans Thermosensory Circuit Regulates Longevity through crh-1/CREB-Dependent flp-6 Neuropeptide Signaling. Developmental cell 39, 209–223. 10.1016/j.devcel.2016.08.021.

Clancy, D.J., Gems, D., Harshman, L.G., Oldham, S., Stocker, H., Hafen, E., Leevers, S.J., and Partridge, L. (2001). Extension of life-span by loss of CHICO, a Drosophila insulin receptor substrate protein. Science 292, 104–106.

Coburn, C.M., and Bargmann, C.I. (1996). A putative cyclic nucleotide-gated channel is required for sensory development and function in C. elegans. Neuron 17, 695–706.

Crombie, T.A., Zdraljevic, S., Cook, D.E., Tanny, R.E., Brady, S.C., Wang, Y., Evans, K.S., Hahnel, S., Lee, D., Rodriguez, B.C., et al. (2019). Deep sampling of Hawaiian Caenorhabditis elegans reveals high genetic diversity and admixture with global populations. eLife 8. 10.7554/eLife.50465.

Dash, B., and Phillips, T.D. (2012). Molecular characterization of a catalase from Hydra vulgaris. Gene 501, 144–152. 10.1016/j.gene.2012.04.015.

Deegenaars, M.L., and Watson, K. (1997). Stress proteins and stress tolerance in an Antarctic, psychrophilic yeast, Candida psychrophila. FEMS Microbiol Lett 151, 191–196. 10.1111/j.1574-6968.1997.tb12569.x.

Dobin, A., Davis, C.A., Schlesinger, F., Drenkow, J., Zaleski, C., Jha, S., Batut, P., Chaisson, M., and Gingeras, T.R. (2013). STAR: ultrafast universal RNA-seq aligner. Bioinformatics 29, 15–21. 10.1093/bioinformatics/bts635.

Doonan, R., McElwee, J.J., Matthijssens, F., Walker, G.A., Houthoofd, K., Back, P., Matscheski, A., Vanfleteren, J.R., and Gems, D. (2008). Against the oxidative damage theory of aging: superoxide dismutases protect against oxidative stress but have little or no effect on life span in Caenorhabditis elegans. Genes & development 22, 3236–3241. 10.1101/gad.504808.

Engelmann, S., Lindner, C., and Hecker, M. (1995). Cloning, nucleotide sequence, and regulation of katE encoding a sigma B-dependent catalase in Bacillus subtilis. Journal of bacteriology 177, 5598–5605. 10.1128/jb.177.19.5598-5605.1995.

Entchev, E.V., Patel, D.S., Zhan, M., Steele, A.J., Lu, H., and Ch’ng, Q. (2015). A gene-expression-based neural code for food abundance that modulates lifespan. eLife 4, e06259. 10.7554/eLife.06259.

Eom, H.J., Kim, H., Kim, B.M., Chon, T.S., and Choi, J. (2014). Integrative assessment of benzene exposure to Caenorhabditis elegans using computational behavior and toxicogenomic analyses. Environ Sci Technol 48, 8143–8151. 10.1021/es500608e.

Evans, M.G., and Polanyi, M. (1935). Some applications of the transition state method to the calculation of reaction velocities, especially in solution. Transactions of the Faraday Society 31, 875–894.

Ewald, C.Y., Landis, J.N., Porter Abate, J., Murphy, C.T., and Blackwell, T.K. (2015). Dauer-independent insulin/IGF-1-signalling implicates collagen remodelling in longevity. Nature 519, 97–101. 10.1038/nature14021.

Eyring, H. (1935). The activated complex in chemical reactions. The Journal of Chemical Physics 3, 107–115.

Ghanta, K.S., Ishidate, T., and Mello, C.C. (2021). Microinjection for precision genome editing in Caenorhabditis elegans. STAR Protoc 2, 100748. 10.1016/j.xpro.2021.100748.

Ghazi, A., Henis-Korenblit, S., and Kenyon, C. (2009). A transcription elongation factor that links signals from the reproductive system to lifespan extension in Caenorhabditis elegans. PLoS genetics 5, e1000639. 10.1371/journal.pgen.1000639.

Glauser, D.A., Chen, W.C., Agin, R., Macinnis, B.L., Hellman, A.B., Garrity, P.A., Tan, M.W., and Goodman, M.B. (2011). Heat avoidance is regulated by transient receptor potential (TRP) channels and a neuropeptide signaling pathway in Caenorhabditis elegans. Genetics 188, 91–103. 10.1534/genetics.111.127100.

Golden, J.W., and Riddle, D.L. (1984). The Caenorhabditis elegans dauer larva: developmental effects of pheromone, food, and temperature. Developmental biology 102, 368–378. 10.1016/0012-1606(84)90201-x.

Gomez-Orte, E., Cornes, E., Zheleva, A., Saenz-Narciso, B., de Toro, M., Iniguez, M., Lopez, R., San-Juan, J.F., Ezcurra, B., Sacristan, B., et al. (2018). Effect of the diet type and temperature on the C. elegans transcriptome. Oncotarget 9, 9556–9571. 10.18632/oncotarget.23563.

Goodman, M.B., and Sengupta, P. (2019). How Caenorhabditis elegans Senses Mechanical Stress, Temperature, and Other Physical Stimuli. Genetics 212, 25–51. 10.1534/genetics.118.300241.

Greiss, S., Schumacher, B., Grandien, K., Rothblatt, J., and Gartner, A. (2008). Transcriptional profiling in C. elegans suggests DNA damage dependent apoptosis as an ancient function of the p53 family. BMC Genomics 9, 334. 10.1186/1471-2164-9-334.

Hedgecock, E.M., and Russell, R.L. (1975). Normal and mutant thermotaxis in the nematode Caenorhabditis elegans. Proc Natl Acad Sci U S A 72, 4061–4065. 10.1073/pnas.72.10.4061.

Higgins, D.P., Weisman, C.M., Lui, D.S., D’Agostino, F.A., and Walker, A.K. (2021). WormCat 2.0 defines characteristics and conservation of poorly annotated genes in *Caenorhabditis elegans*. bioRxiv.

Holdorf, A.D., Higgins, D.P., Hart, A.C., Boag, P.R., Pazour, G.J., Walhout, A.J.M., and Walker, A.K. (2020). WormCat: An Online Tool for Annotation and Visualization of Caenorhabditis elegans Genome-Scale Data. Genetics 214, 279–294. 10.1534/genetics.119.302919.

Holzenberger, M., Dupont, J., Ducos, B., Leneuve, P., Geloen, A., Even, P.C., Cervera, P., and Le Bouc, Y. (2003). IGF-1 receptor regulates lifespan and resistance to oxidative stress in mice. Nature 421, 182–187.

Hourihan, J.M., Moronetti Mazzeo, L.E., Fernandez-Cardenas, L.P., and Blackwell, T.K. (2016). Cysteine Sulfenylation Directs IRE-1 to Activate the SKN-1/Nrf2 Antioxidant Response. Mol Cell 63, 553–566. 10.1016/j.molcel.2016.07.019.

Hu, Z., Li, J., Ding, S., Cheng, F., Li, X., Jiang, Y., Yu, J., Foyer, C.H., and Shi, K. (2021). The protein kinase CPK28 phosphorylates ascorbate peroxidase and enhances thermotolerance in tomato. Plant Physiol 186, 1302–1317. 10.1093/plphys/kiab120.

Huffman, D.L., Abrami, L., Sasik, R., Corbeil, J., van der Goot, F.G., and Aroian, R.V. (2004). Mitogen-activated protein kinase pathways defend against bacterial pore-forming toxins. Proc Natl Acad Sci U S A 101, 10995–11000. 10.1073/pnas.0404073101.

Imlay, J.A. (2018). Where in the world do bacteria experience oxidative stress? Environmental microbiology. 10.1111/1462-2920.14445.

Jansen, W.T., Bolm, M., Balling, R., Chhatwal, G.S., and Schnabel, R. (2002). Hydrogen peroxide-mediated killing of Caenorhabditis elegans by Streptococcus pyogenes. Infection and immunity 70, 5202–5207.

Kamath, R.S., Martinez-Campos, M., Zipperlen, P., Fraser, A.G., and Ahringer, J. (2001). Effectiveness of specific RNA-mediated interference through ingested double-stranded RNA in Caenorhabditis elegans. Genome Biol 2, research2.1-2.10.

Kenyon, C., Chang, J., Gensch, E., Rudner, A., and Tabtiang, R. (1993). A C. elegans mutant that lives twice as long as wild type. Nature 366, 461–464.

Kim, K., and Li, C. (2004). Expression and regulation of an FMRFamide-related neuropeptide gene family in Caenorhabditis elegans. J Comp Neurol 475, 540–550. 10.1002/cne.20189.

Kimura, K.D., Miyawaki, A., Matsumoto, K., and Mori, I. (2004). The C. elegans thermosensory neuron AFD responds to warming. Curr Biol 14, 1291–1295. 10.1016/j.cub.2004.06.060.

Klass, M.R. (1977). Aging in the nematode Caenorhabditis elegans: major biological and environmental factors influencing life span. Mechanisms of ageing and development 6, 413–429.

Kniazeva, M., and Ruvkun, G. (2019). Rhizobium induces DNA damage in Caenorhabditis elegans intestinal cells. Proc Natl Acad Sci U S A 116, 3784–3792. 10.1073/pnas.1815656116.

Komatsu, H., Mori, I., Rhee, J.S., Akaike, N., and Ohshima, Y. (1996). Mutations in a cyclic nucleotide-gated channel lead to abnormal thermosensation and chemosensation in C. elegans. Neuron 17, 707–718.

Kramer-Drauberg, M., Liu, J.L., Desjardins, D., Wang, Y., Branicky, R., and Hekimi, S. (2020). ROS regulation of RAS and vulva development in Caenorhabditis elegans. PLoS genetics 16, e1008838. 10.1371/journal.pgen.1008838.

Kuhara, A., Okumura, M., Kimata, T., Tanizawa, Y., Takano, R., Kimura, K.D., Inada, H., Matsumoto, K., and Mori, I. (2008). Temperature sensing by an olfactory neuron in a circuit controlling behavior of C. elegans. Science 320, 803–807. 10.1126/science.1148922.

Kumar, N., Jain, V., Singh, A., Jagtap, U., Verma, S., and Mukhopadhyay, A. (2015). Genome-wide endogenous DAF-16/FOXO recruitment dynamics during lowered insulin signalling in C. elegans. Oncotarget 6, 41418–41433. 10.18632/oncotarget.6282.

Lee, S.J., and Kenyon, C. (2009). Regulation of the longevity response to temperature by thermosensory neurons in Caenorhabditis elegans. Curr Biol 19, 715–722, 2009. S0960-9822(09)00894-X [pii] 10.1016/j.cub.2009.03.041.

Leiser, S.F., Jafari, G., Primitivo, M., Sutphin, G.L., Dong, J., Leonard, A., Fletcher, M., and Kaeberlein, M. (2016). Age-associated vulval integrity is an important marker of nematode healthspan. Age 38, 419–431. 10.1007/s11357-016-9936-8.

Levins, R. (1968). Evolution in changing environments : some theoretical explorations (Princeton University Press).

Lewis, J.A., Szilagyi, M., Gehman, E., Dennis, W.E., and Jackson, D.A. (2009). Distinct patterns of gene and protein expression elicited by organophosphorus pesticides in Caenorhabditis elegans. BMC Genomics 10, 202. 10.1186/1471-2164-10-202.

Liao, Y., Smyth, G.K., and Shi, W. (2014). featureCounts: an efficient general purpose program for assigning sequence reads to genomic features. Bioinformatics 30, 923–930. 10.1093/bioinformatics/btt656.

Lin, K., Dorman, J.B., Rodan, A., and Kenyon, C. (1997). daf-16: An HNF-3/forkhead family member that can function to double the life-span of Caenorhabditis elegans. Science 278, 1319–1322.

Liu, S., Schulze, E., and Baumeister, R. (2012). Temperature- and touch-sensitive neurons couple CNG and TRPV channel activities to control heat avoidance in Caenorhabditis elegans. PLoS One 7, e32360. 10.1371/journal.pone.0032360.

Loew, O. (1901). Catalase, a new enzym of general occurrence, with special reference to the tobacco plant (Govt. print. off.).

Love, M.I., Huber, W., and Anders, S. (2014). Moderated estimation of fold change and dispersion for RNA-seq data with DESeq2. Genome Biol 15, 550. 10.1186/s13059-014-0550-8.

Lun, A.T., McCarthy, D.J., and Marioni, J.C. (2016). A step-by-step workflow for low-level analysis of single-cell RNA-seq data with Bioconductor. F1000Res 5, 2122. 10.12688/f1000research.9501.2.

Ly, K., Reid, S.J., and Snell, R.G. (2015). Rapid RNA analysis of individual Caenorhabditis elegans. MethodsX 2, 59–63. 10.1016/j.mex.2015.02.002.

McCarroll, S.A., Murphy, C.T., Zou, S., Pletcher, S.D., Chin, C.S., Jan, Y.N., Kenyon, C., Bargmann, C.I., and Li, H. (2004). Comparing genomic expression patterns across species identifies shared transcriptional profile in aging. Nature genetics 36, 197–204. 10.1038/ng1291.

Meng, J., Fu, L., Liu, K., Tian, C., Wu, Z., Jung, Y., Ferreira, R.B., Carroll, K.S., Blackwell, T.K., and Yang, J. (2021). Global profiling of distinct cysteine redox forms reveals wide-ranging redox regulation in C. elegans. Nature communications 12, 1415. 10.1038/s41467-021-21686-3.

Mishra, S., and Imlay, J. (2012). Why do bacteria use so many enzymes to scavenge hydrogen peroxide? Archives of biochemistry and biophysics 525, 145–160. 10.1016/j.abb.2012.04.014.

Mitchell, A., and Pilpel, Y. (2011). A mathematical model for adaptive prediction of environmental changes by microorganisms. Proc Natl Acad Sci U S A 108, 7271–7276. 10.1073/pnas.1019754108.

Mitchell, A., Romano, G.H., Groisman, B., Yona, A., Dekel, E., Kupiec, M., Dahan, O., and Pilpel, Y. (2009). Adaptive prediction of environmental changes by microorganisms. Nature 460, 220–224. 10.1038/nature08112.

Mori, I., and Ohshima, Y. (1995). Neural regulation of thermotaxis in Caenorhabditis elegans. Nature 376, 344–348. 10.1038/376344a0.

Mossialos, D., Tavankar, G.R., Zlosnik, J.E., and Williams, H.D. (2006). Defects in a quinol oxidase lead to loss of KatC catalase activity in Pseudomonas aeruginosa: KatC activity is temperature dependent and it requires an intact disulphide bond formation system. Biochem Biophys Res Commun 341, 697–702. 10.1016/j.bbrc.2005.12.225.

Moy, T.I., Mylonakis, E., Calderwood, S.B., and Ausubel, F.M. (2004). Cytotoxicity of hydrogen peroxide produced by Enterococcus faecium. Infection and immunity 72, 4512–4520. 10.1128/IAI.72.8.4512-4520.2004.

Mueller, M.M., Castells-Roca, L., Babu, V., Ermolaeva, M.A., Muller, R.U., Frommolt, P., Williams, A.B., Greiss, S., Schneider, J.I., Benzing, T., et al. (2014). DAF-16/FOXO and EGL-27/GATA promote developmental growth in response to persistent somatic DNA damage. Nature cell biology 16, 1168–1179. 10.1038/ncb3071.

Murphy, C.T., McCarroll, S.A., Bargmann, C.I., Fraser, A., Kamath, R.S., Ahringer, J., Li, H., and Kenyon, C. (2003). Genes that act downstream of DAF-16 to influence the lifespan of Caenorhabditis elegans. Nature 424, 277–283.

Nicholls, P. (2012). Classical catalase: ancient and modern. Archives of biochemistry and biophysics 525, 95–101. 10.1016/j.abb.2012.01.015.

Nishizawa, A., Yabuta, Y., Yoshida, E., Maruta, T., Yoshimura, K., and Shigeoka, S. (2006). Arabidopsis heat shock transcription factor A2 as a key regulator in response to several types of environmental stress. The Plant journal : for cell and molecular biology 48, 535–547. 10.1111/j.1365-313X.2006.02889.x.

Ogg, S., Paradis, S., Gottlieb, S., Patterson, G.I., Lee, L., Tissenbaum, H.A., and Ruvkun, G. (1997). The Fork head transcription factor DAF-16 transduces insulin-like metabolic and longevity signals in C. elegans. Nature 389, 994–999.

Oliveira, R.P., Porter Abate, J., Dilks, K., Landis, J., Ashraf, J., Murphy, C.T., and Blackwell, T.K. (2009). Condition-adapted stress and longevity gene regulation by Caenorhabditis elegans SKN-1/Nrf. Aging Cell 8, 524–541. ACE501 [pii] 10.1111/j.1474-9726.2009.00501.x.

Pallepati, P., and Averill-Bates, D. (2010). Mild thermotolerance induced at 40 degrees C increases antioxidants and protects HeLa cells against mitochondrial apoptosis induced by hydrogen peroxide: Role of p53. Archives of biochemistry and biophysics 495, 97–111. 10.1016/j.abb.2009.12.014.

Panchuk, II, Volkov, R.A., and Schoffl, F. (2002). Heat stress- and heat shock transcription factor-dependent expression and activity of ascorbate peroxidase in Arabidopsis. Plant Physiol 129, 838–853. 10.1104/pp.001362.

Passardi, F., Zamocky, M., Favet, J., Jakopitsch, C., Penel, C., Obinger, C., and Dunand, C. (2007). Phylogenetic distribution of catalase-peroxidases: are there patches of order in chaos? Gene 397, 101–113. 10.1016/j.gene.2007.04.016.

Patterson, G.I., Koweek, A., Wong, A., Liu, Y., and Ruvkun, G. (1997). The DAF-3 Smad protein antagonizes TGF-beta-related receptor signaling in the Caenorhabditis elegans dauer pathway. Genes & development 11, 2679–2690.

Petriv, O.I., and Rachubinski, R.A. (2004). Lack of peroxisomal catalase causes a progeric phenotype in Caenorhabditis elegans. J Biol Chem 279, 19996–20001. 10.1074/jbc.M400207200.

Picelli, S., Faridani, O.R., Bjorklund, A.K., Winberg, G., Sagasser, S., and Sandberg, R. (2014). Full-length RNA-seq from single cells using Smart-seq2. Nat Protoc 9, 171–181. 10.1038/nprot.2014.006.

Prahlad, V., Cornelius, T., and Morimoto, R.I. (2008). Regulation of the cellular heat shock response in Caenorhabditis elegans by thermosensory neurons. Science 320, 811–814. 10.1126/science.1156093.

Ramot, D., MacInnis, B.L., and Goodman, M.B. (2008). Bidirectional temperature-sensing by a single thermosensory neuron in C. elegans. Nature neuroscience 11, 908–915. 10.1038/nn.2157.

Rodaki, A., Bohovych, I.M., Enjalbert, B., Young, T., Odds, F.C., Gow, N.A., and Brown, A.J. (2009). Glucose promotes stress resistance in the fungal pathogen Candida albicans. Mol Biol Cell 20, 4845–4855. 10.1091/mbc.E09-01-0002.

Rosso, L., Lobry, J.R., and Flandrois, J.P. (1993). An unexpected correlation between cardinal temperatures of microbial growth highlighted by a new model. Journal of theoretical biology 162, 447–463. 10.1006/jtbi.1993.1099.

Rual, J.F., Ceron, J., Koreth, J., Hao, T., Nicot, A.S., Hirozane-Kishikawa, T., Vandenhaute, J., Orkin, S.H., Hill, D.E., van den Heuvel, S., and Vidal, M. (2004). Toward improving Caenorhabditis elegans phenome mapping with an ORFeome-based RNAi library. Genome research 14, 2162–2168. 10.1101/gr.2505604.

Sahu, S.N., Lewis, J., Patel, I., Bozdag, S., Lee, J.H., Sprando, R., and Cinar, H.N. (2013). Genomic analysis of stress response against arsenic in Caenorhabditis elegans. PLoS One 8, e66431. 10.1371/journal.pone.0066431.

Samuel, B.S., Rowedder, H., Braendle, C., Felix, M.A., and Ruvkun, G. (2016). Caenorhabditis elegans responses to bacteria from its natural habitats. Proc Natl Acad Sci U S A 113, E3941–3949. 10.1073/pnas.1607183113.

Satterlee, J.S., Sasakura, H., Kuhara, A., Berkeley, M., Mori, I., and Sengupta, P. (2001). Specification of thermosensory neuron fate in C. elegans requires ttx-1, a homolog of otd/Otx. Neuron 31, 943–956.

Schiffer, J.A., Servello, F.A., Heath, W.R., Amrit, F.R.G., Stumbur, S.V., Eder, M., Martin, O.M., Johnsen, S.B., Stanley, J.A., Tam, H., et al. (2020). Caenorhabditis elegans processes sensory information to choose between freeloading and self-defense strategies. eLife 9. 10.7554/eLife.56186.

Schiffer, J.A., Stumbur, S.V., Seyedolmohadesin, M., Xu, Y.Y., Serkin, W.T., McGowan, N.G., Banjo, O., Torkashvand, M., Lin, A.L., Hosea, C.N., et al. (2021). Modulation of sensory perception by hydrogen peroxide enables Caenorhabditis elegans to find a niche that provides both food and protection from hydrogen peroxide. PLoS pathogens 17. ARTN e1010112. 10.1371/journal.ppat.1010112.

Schild, L.C., Zbinden, L., Bell, H.W., Yu, Y.V., Sengupta, P., Goodman, M.B., and Glauser, D.A. (2014). The balance between cytoplasmic and nuclear CaM kinase-1 signaling controls the operating range of noxious heat avoidance. Neuron 84, 983–996. 10.1016/j.neuron.2014.10.039.

Schild, S., Tamayo, R., Nelson, E.J., Qadri, F., Calderwood, S.B., and Camilli, A. (2007). Genes induced late in infection increase fitness of Vibrio cholerae after release into the environment. Cell Host Microbe 2, 264–277. 10.1016/j.chom.2007.09.004.

Seaver, L.C., and Imlay, J.A. (2001). Alkyl hydroperoxide reductase is the primary scavenger of endogenous hydrogen peroxide in Escherichia coli. Journal of bacteriology 183, 7173–7181. 10.1128/JB.183.24.7173-7181.2001.

Servello, F.A., and Apfeld, J. (2020). The heat shock transcription factor HSF-1 protects Caenorhabditis elegans from peroxide stress. Translational Medicine of Aging 4, 88–92.

Shivers, R.P., Kooistra, T., Chu, S.W., Pagano, D.J., and Kim, D.H. (2009). Tissue-specific activities of an immune signaling module regulate physiological responses to pathogenic and nutritional bacteria in C. elegans. Cell Host Microbe 6, 321–330. 10.1016/j.chom.2009.09.001.

Starnes, D.L., Lichtenberg, S.S., Unrine, J.M., Starnes, C.P., Oostveen, E.K., Lowry, G.V., Bertsch, P.M., and Tsyusko, O.V. (2016). Distinct transcriptomic responses of Caenorhabditis elegans to pristine and sulfidized silver nanoparticles. Environ Pollut 213, 314–321. 10.1016/j.envpol.2016.01.020.

Steinbaugh, M.J., Narasimhan, S.D., Robida-Stubbs, S., Moronetti Mazzeo, L.E., Dreyfuss, J.M., Hourihan, J.M., Raghavan, P., Operana, T.N., Esmaillie, R., and Blackwell, T.K. (2015). Lipid-mediated regulation of SKN-1/Nrf in response to germ cell absence. eLife 4. 10.7554/eLife.07836.

Stroustrup, N., Anthony, W.E., Nash, Z.M., Gowda, V., Gomez, A., Lopez-Moyado, I.F., Apfeld, J., and Fontana, W. (2016). The temporal scaling of Caenorhabditis elegans ageing. Nature 530, 103–107. 10.1038/nature16550.

Stroustrup, N., Ulmschneider, B.E., Nash, Z.M., Lopez-Moyado, I.F., Apfeld, J., and Fontana, W. (2013). The Caenorhabditis elegans Lifespan Machine. Nat Methods 10, 665–670, 2013. nmeth.2475 [pii] 10.1038/nmeth.2475.

Sugi, T., Nishida, Y., and Mori, I. (2011). Regulation of behavioral plasticity by systemic temperature signaling in Caenorhabditis elegans. Nature neuroscience 14, 984–992. 10.1038/nn.2854.

Supek, F., Bosnjak, M., Skunca, N., and Smuc, T. (2011). REVIGO summarizes and visualizes long lists of gene ontology terms. PLoS One 6, e21800. 10.1371/journal.pone.0021800.

Sze, J.Y., Victor, M., Loer, C., Shi, Y., and Ruvkun, G. (2000). Food and metabolic signalling defects in a Caenorhabditis elegans serotonin-synthesis mutant. Nature 403, 560–564.

Tagkopoulos, I., Liu, Y.C., and Tavazoie, S. (2008). Predictive behavior within microbial genetic networks. Science 320, 1313–1317. 10.1126/science.1154456.

Tatar, M., Bartke, A., and Antebi, A. (2003). The endocrine regulation of aging by insulin-like signals. Science 299, 1346–1351.

Tatum, M.C., Ooi, F.K., Chikka, M.R., Chauve, L., Martinez-Velazquez, L.A., Steinbusch, H.W.M., Morimoto, R.I., and Prahlad, V. (2015). Neuronal serotonin release triggers the heat shock response in C. elegans in the absence of temperature increase. Curr Biol 25, 163–174. 10.1016/j.cub.2014.11.040.

Taylor, S.R., Santpere, G., Reilly, M., Glenwinkel, L., Poff, A., McWhirter, R., Xu, C., Weinreb, A., Basavaraju, M., Cook, S.J., et al. (2019). Expression profiling of the mature *C. elegans* nervous system by single-cell RNA-Sequencing. bioRxiv.

Togo, S.H., Maebuchi, M., Yokota, S., Bun-Ya, M., Kawahara, A., and Kamiryo, T. (2000). Immunological detection of alkaline-diaminobenzidine-negativeperoxisomes of the nematode Caenorhabditis elegans purification and unique pH optima of peroxisomal catalase. Eur J Biochem 267, 1307–1312. 10.1046/j.1432-1327.2000.01091.x.

Tullet, J.M., Hertweck, M., An, J.H., Baker, J., Hwang, J.Y., Liu, S., Oliveira, R.P., Baumeister, R., and Blackwell, T.K. (2008). Direct inhibition of the longevity-promoting factor SKN-1 by insulin-like signaling in C. elegans. Cell 132, 1025–1038. S0092-8674(08)00130-X [pii] 10.1016/j.cell.2008.01.030.

Tursun, B., Cochella, L., Carrera, I., and Hobert, O. (2009). A toolkit and robust pipeline for the generation of fosmid-based reporter genes in C. elegans. PLoS One 4, e4625. 10.1371/journal.pone.0004625.

Veal, E.A., Day, A.M., and Morgan, B.A. (2007). Hydrogen peroxide sensing and signaling. Mol Cell 26, 1–14, 2007. S1097-2765(07)00186-4 [pii] 10.1016/j.molcel.2007.03.016.

Wieser, R., Adam, G., Wagner, A., Schuller, C., Marchler, G., Ruis, H., Krawiec, Z., and Bilinski, T. (1991). Heat shock factor-independent heat control of transcription of the CTT1 gene encoding the cytosolic catalase T of Saccharomyces cerevisiae. J Biol Chem 266, 12406–12411.

Wolf, M., Nunes, F., Henkel, A., Heinick, A., and Paul, R.J. (2008). The MAP kinase JNK-1 of Caenorhabditis elegans: location, activation, and influences over temperature-dependent insulin-like signaling, stress responses, and fitness. J Cell Physiol 214, 721–729. 10.1002/jcp.21269.

Xiao, R., Zhang, B., Dong, Y., Gong, J., Xu, T., Liu, J., and Xu, X.Z. (2013). A genetic program promotes C. elegans longevity at cold temperatures via a thermosensitive TRP channel. Cell 152, 806–817. 10.1016/j.cell.2013.01.020.

Yang, W., Dierking, K., and Schulenburg, H. (2016). WormExp: a web-based application for a Caenorhabditis elegans-specific gene expression enrichment analysis. Bioinformatics 32, 943–945. 10.1093/bioinformatics/btv667.

Zhang, F., Berg, M., Dierking, K., Felix, M.A., Shapira, M., Samuel, B.S., and Schulenburg, H. (2017). Caenorhabditis elegans as a Model for Microbiome Research. Frontiers in microbiology 8, 485. 10.3389/fmicb.2017.00485.

Zhang, Y., Parmigiani, G., and Johnson, W.E. (2020). ComBat-seq: batch effect adjustment for RNA-seq count data. NAR Genom Bioinform 2, lqaa078. 10.1093/nargab/lqaa078.

