## supplementary information for "Neuronal temperature perception induces specific defenses that enable C. elegans to cope with the enhanced reactivity of hydrogen peroxide at high temperature"

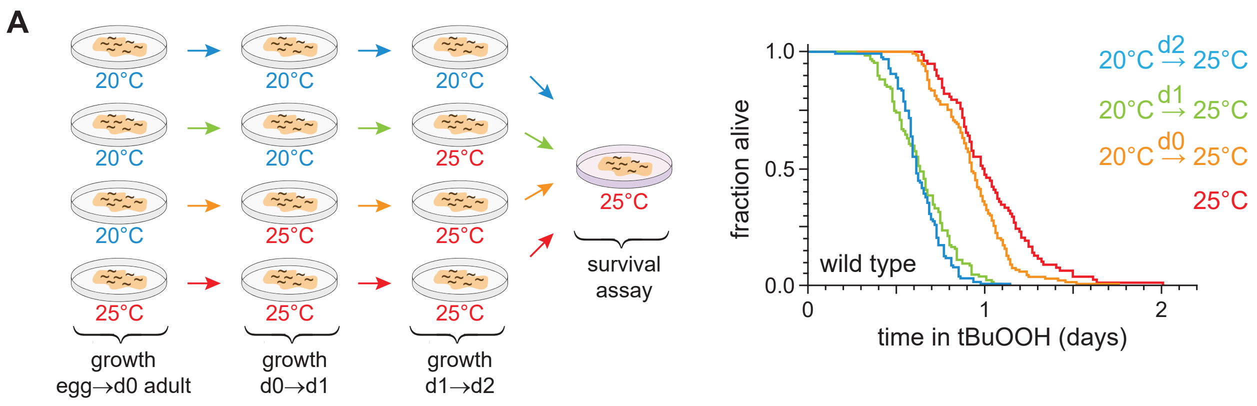

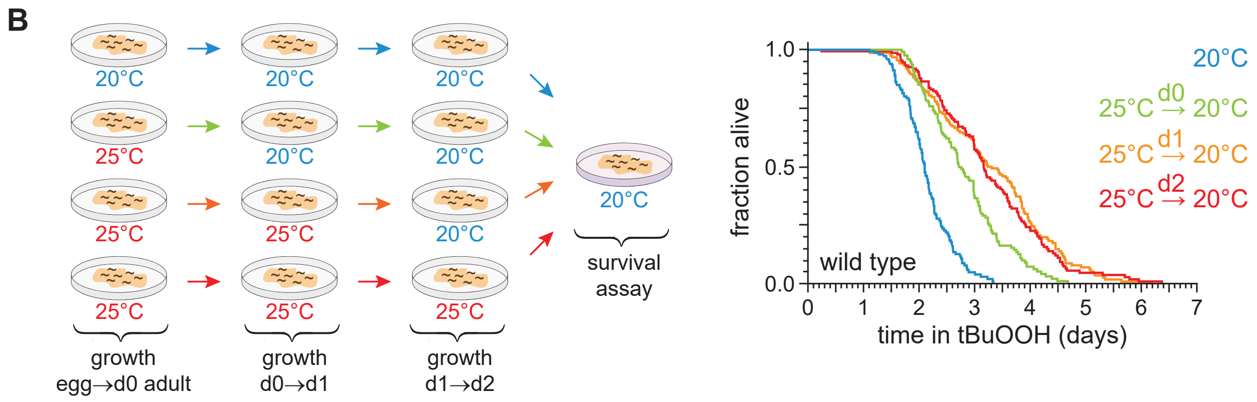

**Supplementary Figure 1.** ***C. elegans*** **can induce lasting peroxide defenses in response to high temperature.**

(A) Peroxide resistance at 25°C of wild-type *C. elegans* shifted from 20°C to 25°C at the specified time before the assay.
(B) Peroxide resistance at 20°C of wild-type *C. elegans* shifted from 25°C to 20°C at the specified time before the assay.
Statistical analyses for panels (A-B) are in Supplementary Table 1.

**A**

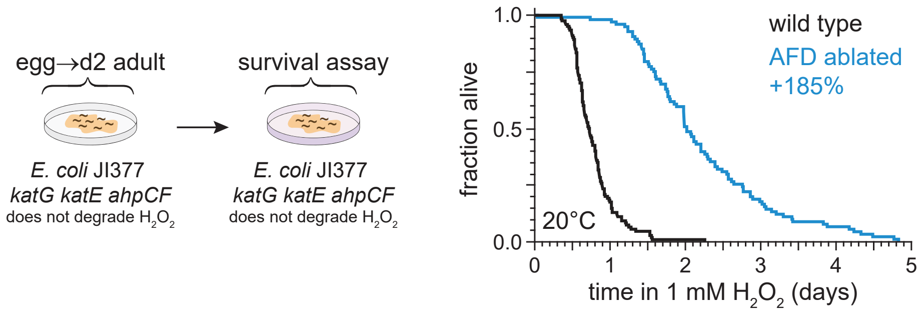

**Supplementary Figure 2. The AFD sensory neurons regulate *C. elegans* $H_2O_2$ resistance.**

(A) Hydrogen peroxide resistance of wild type and AFD-ablated *C. elegans* nematodes at 20°C. The nematodes were grown and assayed on lawns of *E. coli* JI377, a *katG katE ahpCF* triple null mutant strain unable to degrade $H_2O_2$ in the environment (Seaver and Imlay, 2001).
Statistical analysis for panel (A) is in Supplementary Table 2.

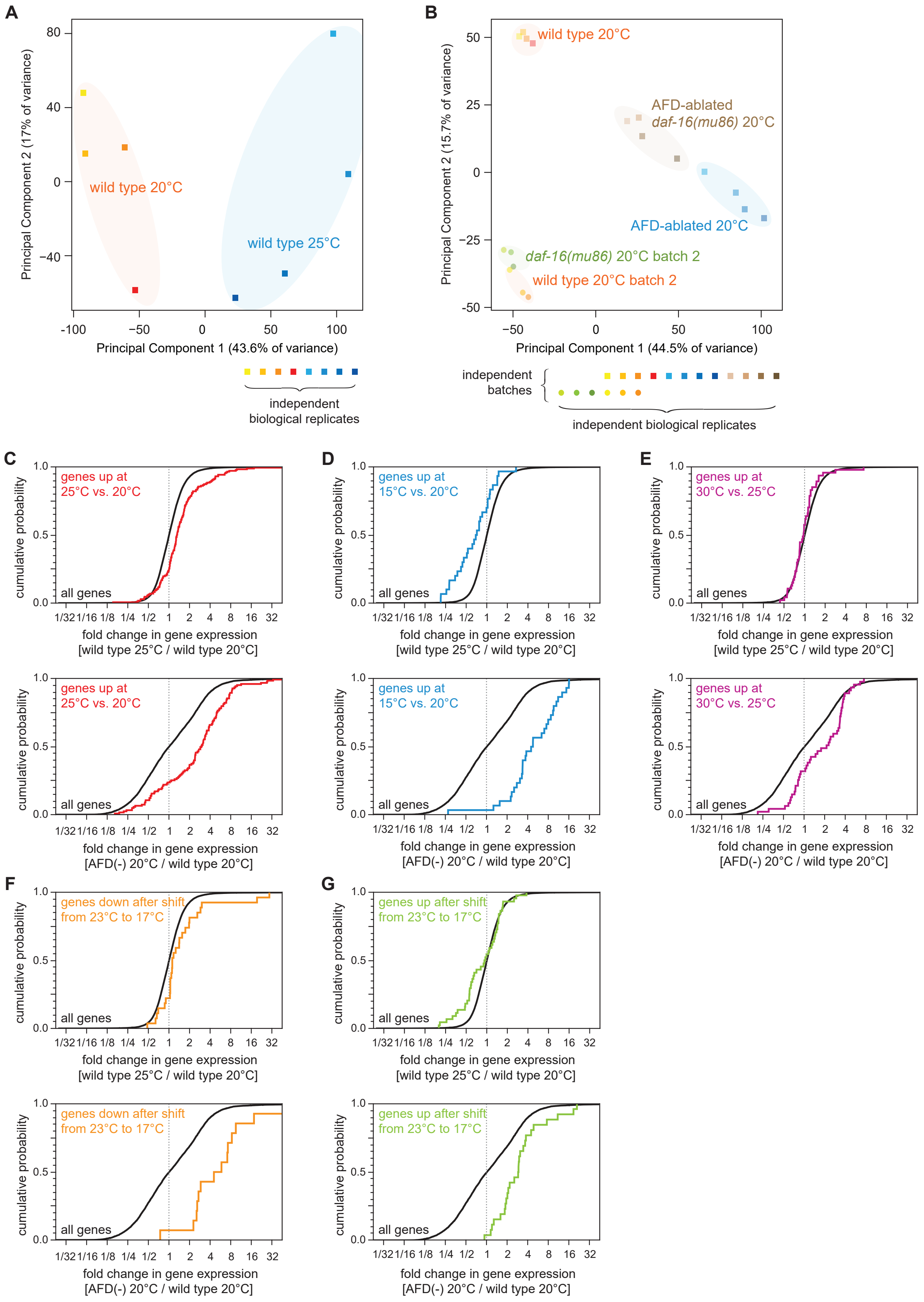

**Supplementary Figure 3. The AFD sensory neurons influence responses to noxious heat, high cultivation temperature, and low temperature.**

(A) Principal component analysis (PCA) of the sequenced samples of wild-type nematodes grown at 25°C and at 20°C.
(B) Principal component analysis (PCA) of the sequenced samples of wild type, *daf-16(mu86)* null mutants, AFD-ablated nematodes, and AFD-ablated *daf-16(mu86)* null mutants grown at 20°C.
(C-G) Effect on the expression of temperature-regulated gene-sets of (top panels) growth at 25°C relative to growth at 20°C in wild-type nematodes and of (bottom panels) AFD ablation in nematodes grown at 20°C relative to wild-type (unablated) nematodes grown at 20°C: (C) genes up at 25°C vs. 20°C (Gomez-Orte *et al.*, 2018); (D) genes up at 15°C vs. 20°C (Gomez-Orte *et al.*, 2018); (E) genes up at 30°C vs. 25°C (McCarroll *et al.*, 2004); (F) genes down after shift from 23°C to 17°C (Sugi *et al.*, 2011); (G) genes up after shift from 23°C to 17°C (Sugi *et al.*, 2011).
Statistical analyses for panels (C-G) are in Supplementary Table 3.

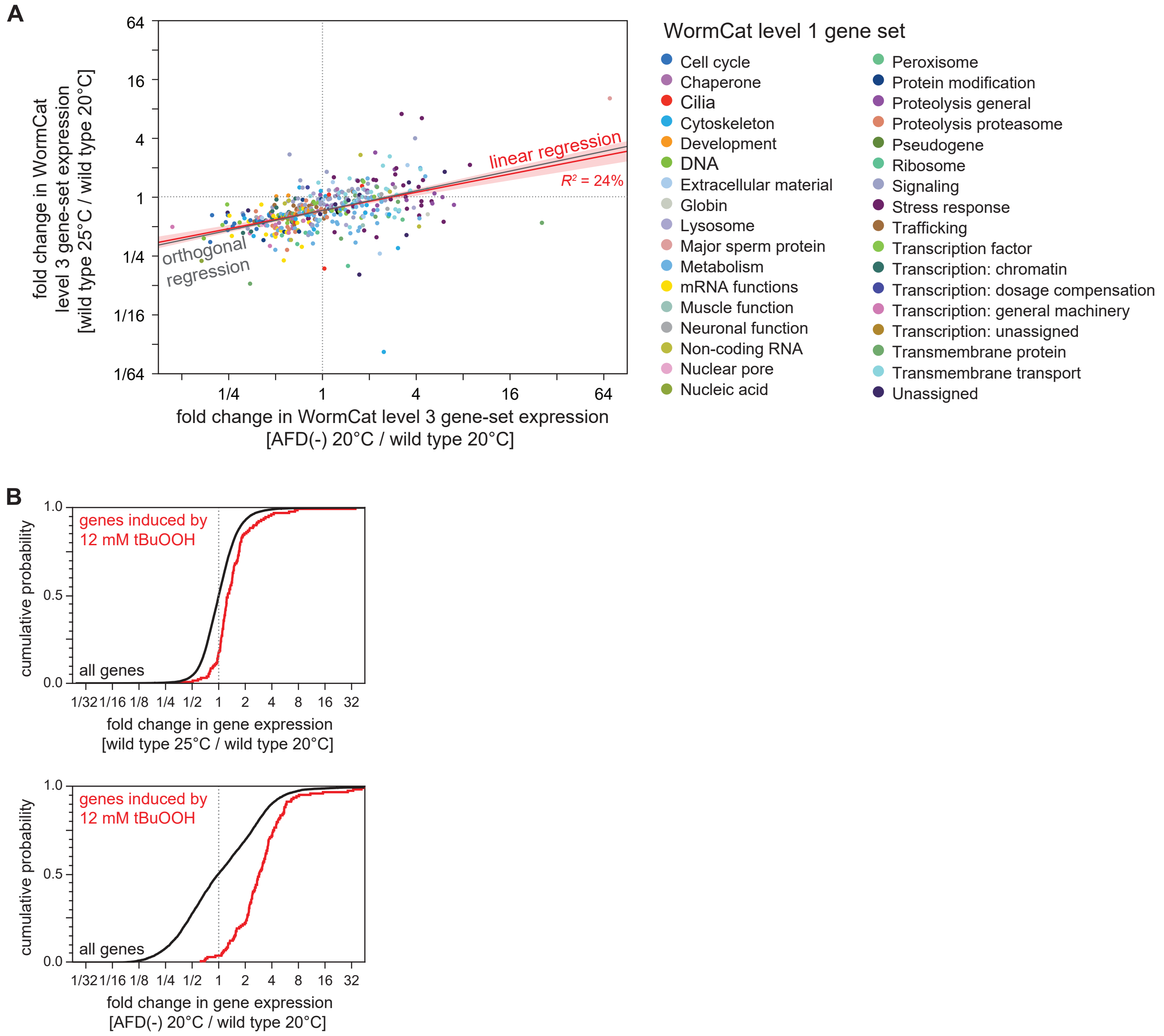

**Supplementary Figure 4. High cultivation temperature and AFD ablation pre-induce genes induced by peroxides.**

(A) Growth at 25°C and AFD ablation at 20°C induced correlated changes in the average gene expression level of gene sets affecting similar biological processes based on WormCat “level 3” nested categories (Higgins *et al.*, 2021; Holdorf *et al.*, 2020). Gene-sets are colored based on their WormCat “level 1” categories. Linear regression fit is shown as a red line flanked by a red area marking the 95% confidence interval of the fit. The orthogonal regression fit (grey line) makes no assumptions about the dependence or independence of the variables.
(B) Effect on the expression of genes induced by 12 mM tBuOOH (Oliveira *et al.*, 2009) of (top panel) growth at 25°C relative to growth at 20°C in wild-type nematodes and of (bottom panel) AFD ablation in nematodes grown at 20°C relative to wild-type (unablated) nematodes grown at 20°C. Statistical analysis for panel (B) is in Supplementary Table 3.

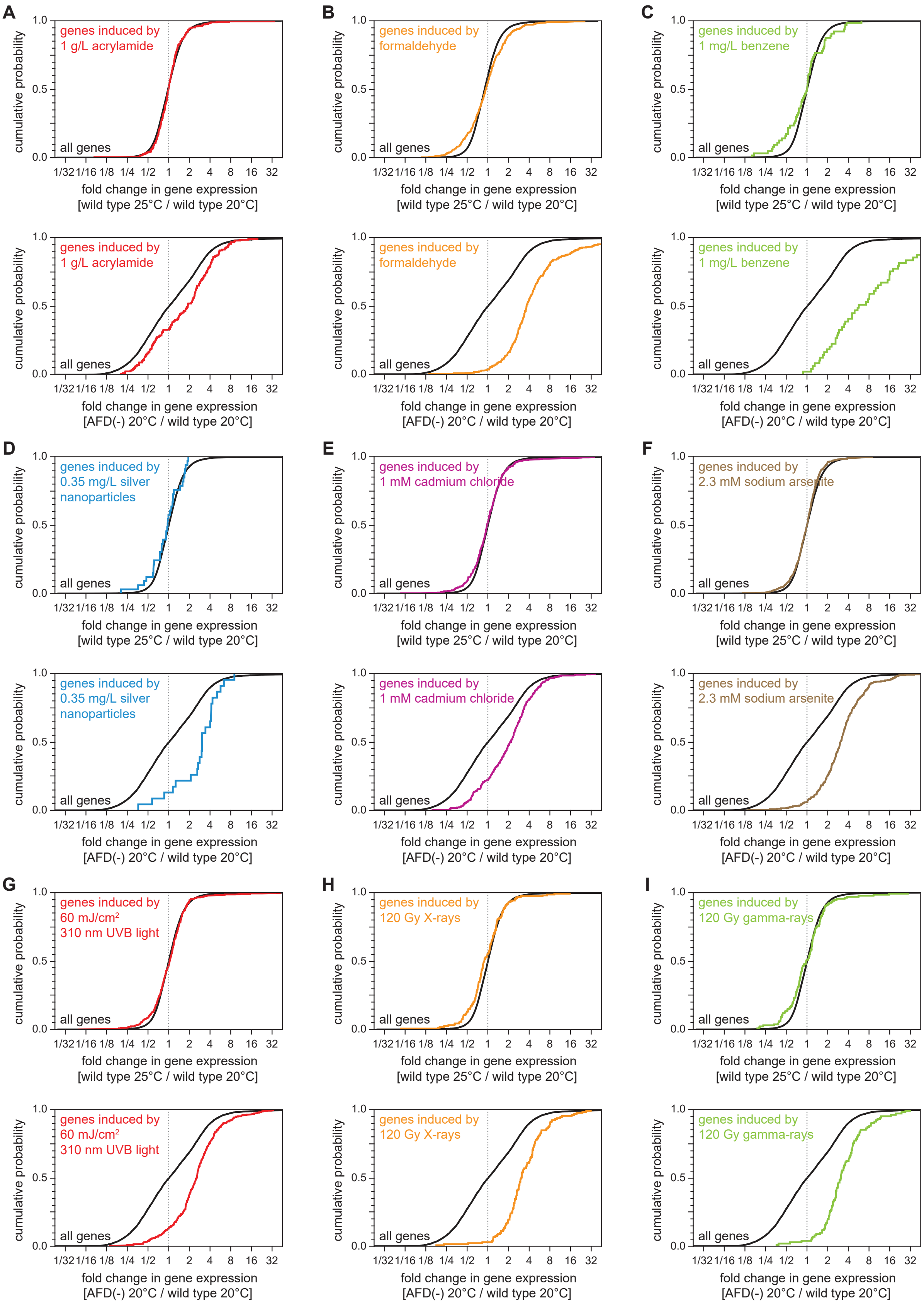

**Supplementary Figure 5. AFD ablation, but not high cultivation temperature, pre-induces genes induced by toxic organic compounds, toxic metals, and radiation.**

(A-I) Effect on the expression of genes induced by toxic organic compounds, toxic metals, and radiation of (top panels) growth at 25°C relative to growth at 20°C in wild-type nematodes and of (bottom panels) AFD ablation in nematodes grown at 20°C relative to wild-type (unablated) nematodes grown at 20°C:

(A) genes induced by 1 g/L acrylamide (Lewis *et al.*, 2009)2009; (B) genes induced by formaldehyde (Yang *et al.*, 2016); (C) genes induced by 1 mg/L benzene (Eom *et al.*, 2014); (D) genes induced by 0.35 mg/L silver nanoparticles (Starnes *et al.*, 2016); (E) genes induced by 1 mM cadmium chloride (Huffman *et al.*, 2004); (F) genes induced by 2.3 mM sodium arsenite (Sahu *et al.*, 2013)2013; (G) genes induced by 60 mJ/cm$^2$ 310 nm UVB light (Mueller *et al.*, 2014); (H) genes induced by 120 Gy X-rays (Greiss *et al.*, 2008); (I) genes induced by 120 Gy gamma-rays (Greiss *et al.*, 2008).

Statistical analyses for panels (A-I) are in Supplementary Table 3.

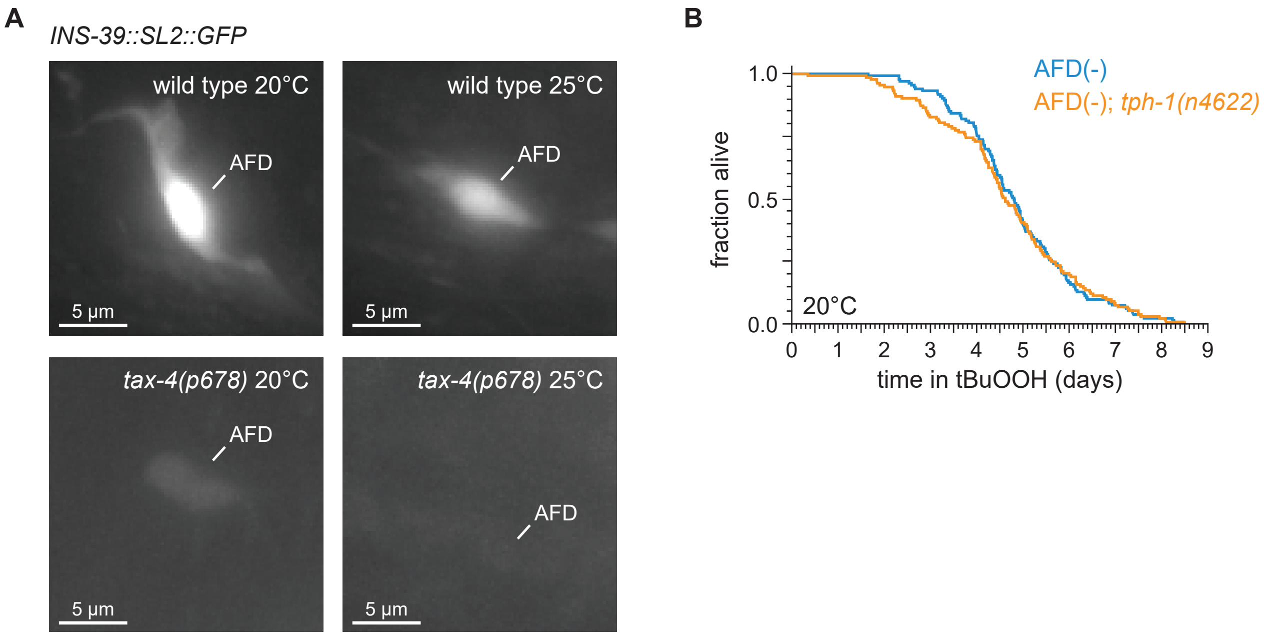

**Supplementary Figure 6. TAX-4 cyclic GMP-gated channels are required for *ins-39* gene expression in the AFD sensory neurons.**

(A) Representative images the expression of the *ins-39(oy167[ins-39::SL2::GFP])* reporter in wild-type nematodes (top panels) and *tax-4(p678)* mutants (bottom panels), grown at 20°C (left panels) and 25°C (right panels) in one of the bilateral AFD neurons. Scale bar = 5 µm.
(B) *tph-1(n4622)* did not affect the increased peroxide resistance of AFD-ablated nematodes at 20°C. Statistical analysis for panel (B) is in Supplementary Table 5.

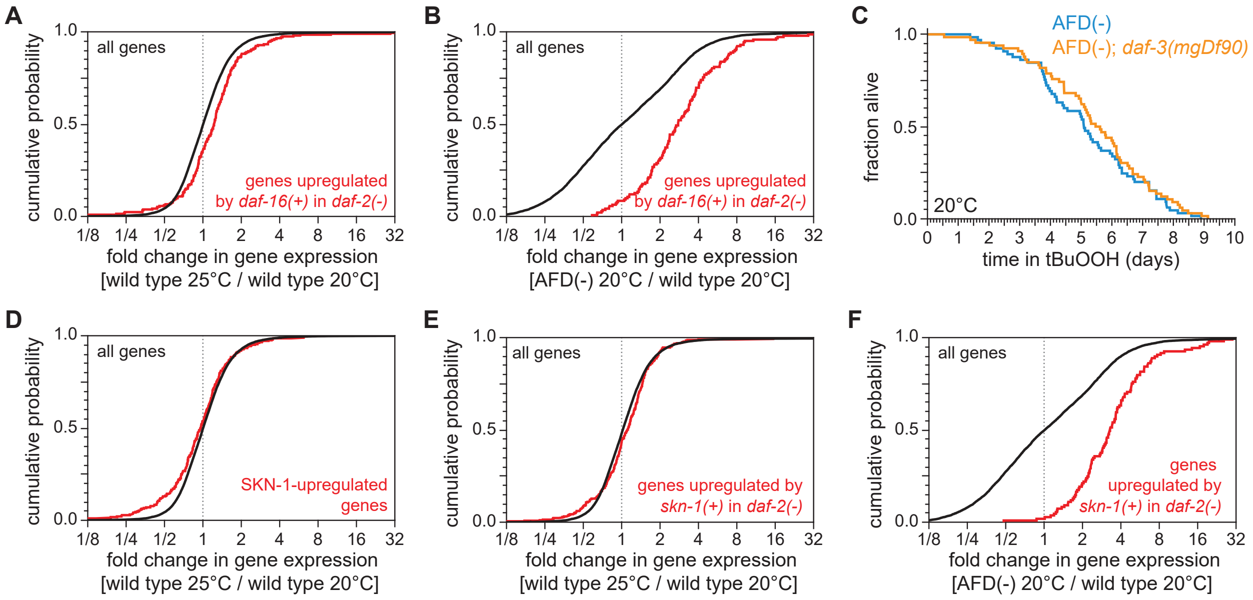

**Supplementary Figure 7. The AFD sensory neurons repress the expression of genes induced by DAF-16/FOXO and SKN-1/NRF.**

(A-B) Genes upregulated in a *daf-16*-dependent manner in *daf-2(-)* mutants (Murphy *et al.*, 2003) had higher expression (A) in nematodes grown at 25°C than in nematodes grown at 20°C and (B) in AFD-ablated nematodes grown at 20°C than in wild-type (unablated) nematodes grown at 20°C.
(C) *daf-3(mgDf90)* did not affect the increased peroxide resistance of AFD-ablated nematodes at 20°C.
(D-E) Genes upregulated by *skn-1(+)* in (D) wild type nematodes (Oliveira *et al.*, 2009) and (E) *daf-2* loss-of-function mutants (Ewald *et al.*, 2015) did not have higher expression in nematodes grown at 25°C than in nematodes grown at 20°C.
(F) Genes upregulated by *skn-1(+)* in *daf-2* loss-of-function mutants (Ewald *et al.*, 2015) had higher expression in AFD-ablated nematodes grown at 20°C than in wild-type (unablated) nematodes grown at 20°C.
Statistical analyses for panels (A-B, D-F) are in Supplementary Table 3 and statistical analysis for panel (C) is in Supplementary Table 7.

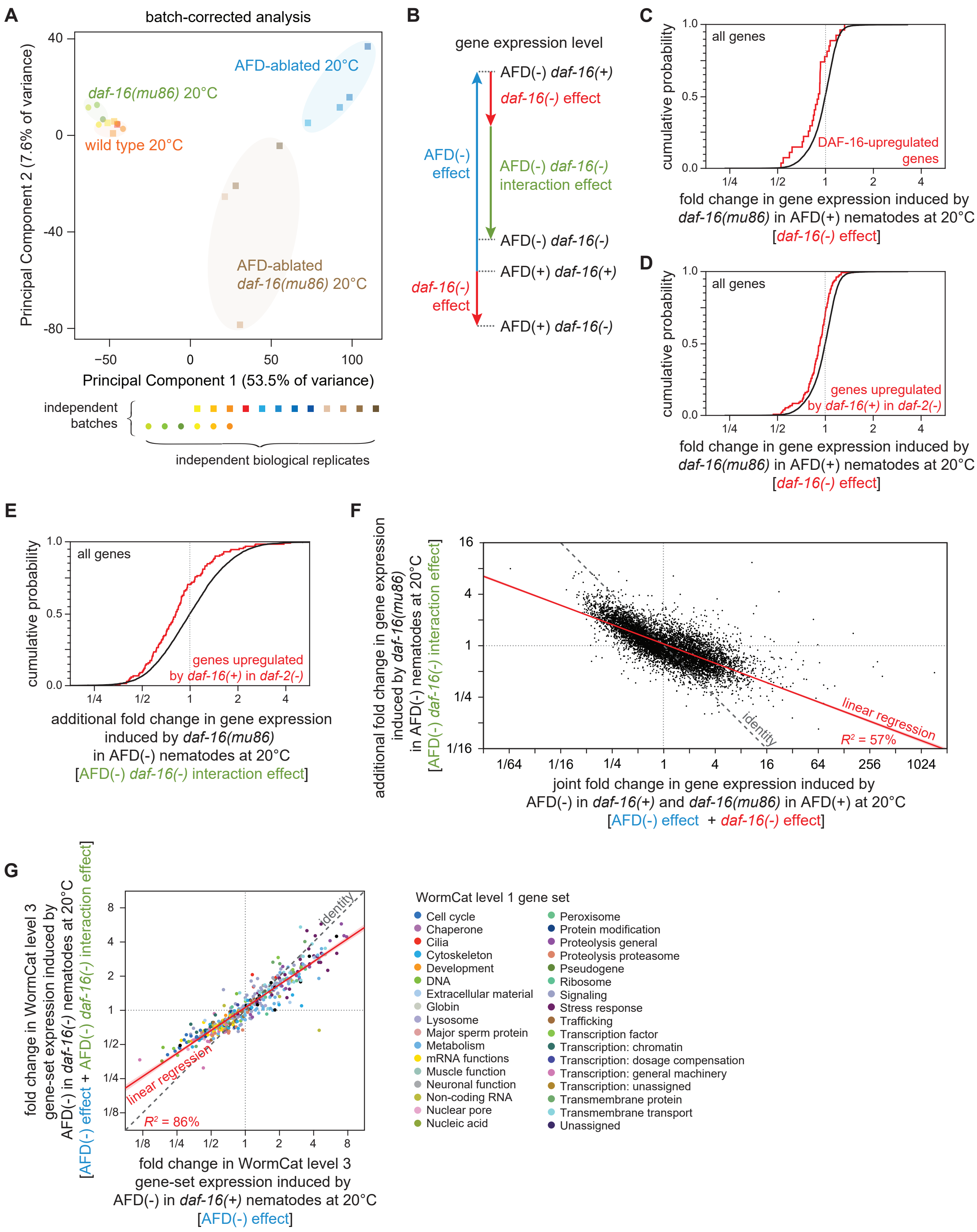

**Supplementary Figure 8. DAF-16/FOXO potentiates the changes in gene-set expression induced by the AFD sensory neurons.**

(A) Principal component analysis (PCA) of the batch-corrected sequenced samples of wild type, *daf-16(mu86)* null mutants, AFD-ablated nematodes, and AFD-ablated *daf-16(mu86)* null mutants grown at 20°C. See Figure S3A for the PCA before batch correction.
(B) We performed mRNA-seq on wild type [AFD(+) *daf-16(+)*], *daf-16(mu86)* null mutants [AFD(+) *daf-16(-)*], AFD-ablated nematodes [AFD(-) *daf-16(+)*], and AFD-ablated *daf-16(mu86)* null mutants [AFD(-) *daf-16(-)*] grown at 20°C, and used an epistasis model to quantify the extent to which AFD-ablation and *daf-16* mutation affected the expression of each gene, relative to wild type, in terms of the independent effects induced by AFD ablation (blue arrow) and by lack *daf-16* gene function (red arrow), and the additional effect induced by the interaction between AFD ablation and lack *daf-16* gene function (green arrow).
(C-D) Effect of lack *daf-16* gene function in unablated nematodes at 20°C on the expression of (C) genes directly upregulated by DAF-16 (Kumar *et al.*, 2015) and (D) genes upregulated in a *daf-16*-dependent manner in *daf-2(-)* mutants (Murphy *et al.*, 2003).
(E) Effect of the additional changes in gene expression induced by lack *daf-16* gene function in AFD-ablated nematodes at 20°C on the expression of genes upregulated in a *daf-16*-dependent manner in *daf-2(-)* mutants (Murphy *et al.*, 2003).
(F) The extent to which lack *daf-16* function had additional effects on gene expression when the AFD neurons were ablated was negatively correlated with the level of gene expression predicted if AFD and DAF-16 acted independently. Linear regression fit is shown as a red line flanked by a red area marking the 95% confidence interval of the fit. The slope and intercept of that line would have been 0, if AFD and DAF-16 had acted independently to influence fold changes in gene expression at the transcriptome level (Angeles-Albores *et al.*, 2018).
(G) The effect of AFD ablation at 20°C on the average gene expression level of gene sets affecting similar biological processes based on WormCat “level 3” nested categories (Higgins *et al.*, 2021; Holdorf *et al.*, 2020) was systematically smaller in *daf-16(mu86)* mutants (y-axis) than in *daf-16(+)* nematodes (x-axis). Gene-sets are colored based on their WormCat “level 1” categories. Linear regression fit is shown as a red line flanked by a red area marking the 95% confidence interval of the fit.
Statistical analyses for panels (C-E) are in Supplementary Table 3.

### Supplementary Table 1. Statistical analysis for Figures 1 and S1.

| Set | Genotype | Temperature: egg → day 2 adult | Temperature: survival assay | Mean survival ± SEM (days) | Median survival (days) | 75th percentile (days) | N dead / initial N | Group | % Mean survival change vs. group a | *P* value (log-rank) vs. group a | Figure |
|---|---|---|---|---|---|---|---|---|---|---|---|
| Survival in 6 mM tBuOOH, *E. coli* OP50 | | | | | | | | | | | |
| | wild type | 20°C | 20°C | 1.60 ± 0.02 | 1.58 | 1.80 | 131 / 145 | a | | | 1A |
| | wild type | 25°C | 25°C | 1.12 ± 0.02 | 1.15 | 1.27 | 86 / 86 | b | -30% | < 0.0001 | |
| | wild type | 20°C | 25°C | 0.68 ± 0.01 | 0.69 | 0.74 | 140 / 140 | a | | | 1C |
| | wild type | 25°C | 25°C | 1.05 ± 0.03 | 1.05 | 1.15 | 65 / 65 | b | 54% | < 0.0001 | |
| | wild type | 20°C | 20°C | 1.58 ± 0.03 | 1.52 | 1.76 | 146 / 146 | a | | | 1D |
| | wild type | 25°C | 20°C | 2.48 ± 0.08 | 2.25 | 3.09 | 131 / 131 | b | 57% | < 0.0001 | |

| Set | Genotype | Temperature: egg → day 0 adult | Temperature: day 0 → day 1 adult | Temperature: day 1 → day 2 adult | Temperature: survival assay | Mean survival ± SEM (days) | Median survival (days) | 75th percentile (days) | N dead / initial N | Group | % Mean survival change vs. group a | *P* value (log-rank) vs. group a | *P* value (log-rank) vs. group b | *P* value (log-rank) vs. group c | Figure |
|---|---|---|---|---|---|---|---|---|---|---|---|---|---|---|---|
| Survival in 6 mM tBuOOH, *E. coli* OP50 | | | | | | | | | | | | | | | |
| | wild type | 25°C | 25°C | 25°C | 25°C | 1.03 ± 0.03 | 0.99 | 1.18 | 78 / 78 | a | | | | | S1A |
| | wild type | 20°C | 25°C | 25°C | 25°C | 0.94 ± 0.02 | 0.93 | 1.06 | 133 / 133 | b | -9% | 0.0034 | | | |
| | wild type | 20°C | 20°C | 25°C | 25°C | 0.64 ± 0.02 | 0.63 | 0.76 | 127 / 127 | c | -38% | < 0.0001 | < 0.0001 | | |
| | wild type | 20°C | 20°C | 20°C | 25°C | 0.64 ± 0.01 | 0.61 | 0.73 | 127 / 127 | d | -38% | < 0.0001 | < 0.0001 | > 0.05 | |
| | wild type | 20°C | 20°C | 20°C | 20°C | 2.14 ± 0.04 | 2.10 | 2.40 | 130 / 155 | a | | | | | S1B |
| | wild type | 25°C | 20°C | 20°C | 20°C | 2.82 ± 0.07 | 2.72 | 3.26 | 119 / 129 | b | 32% | < 0.0001 | | | |
| | wild type | 25°C | 25°C | 20°C | 20°C | 3.29 ± 0.09 | 3.24 | 4.03 | 140 / 157 | c | 54% | < 0.0001 | < 0.0001 | | |
| | wild type | 25°C | 25°C | 25°C | 20°C | 3.26 ± 0.09 | 3.16 | 3.91 | 134 / 152 | d | 53% | < 0.0001 | < 0.0001 | > 0.05 | |

### Supplementary Table 2. Statistical analysis for Figures 2 and S2.

| | | Temperature | | | | | | | | | | | |
|---|---|---|---|---|---|---|---|---|---|---|---|---|---|
| **Set** | **Genotype** | **egg → day 2 adult** | **survival assay** | **Mean survival ± SEM (days)** | **Median survival (days)** | **75th percentile (days)** | **N dead / initial N** | **Group** | **% Mean survival change vs. group a** | ***P* value (log-rank) vs. group a** | ***P* value (log-rank) vs. group b** | ***P* value (log-rank) vs. group c** | **Figure** |
| Survival in 6 mM tBuOOH, E. coli OP50 | | | | | | | | | | | | | |
| | wild type | 20°C | 20°C | 1.85 ± 0.03 | 1.91 | 2.09 | 134 / 134 | a | | | | | 2B |
| | *tax-4(p678) III* | 20°C | 20°C | 4.80 ± 0.08 | 4.76 | 5.40 | 100 / 100 | b | 159% | < 0.0001 | | | |
| | wild type | 25°C | 25°C | 1.16 ± 0.02 | 1.17 | 1.29 | 138 / 151 | a | | | | | 2C |
| | *tax-4(p678) III* | 25°C | 25°C | 1.72 ± 0.03 | 1.74 | 1.97 | 131 / 147 | b | 49% | < 0.0001 | | | |
| | wild type | 20°C | 20°C | 1.98 ± 0.04 | 2.00 | 2.28 | 134 / 134 | a | | | | | 2D |
| | AFD ablated | 20°C | 20°C | 4.75 ± 0.12 | 4.69 | 5.43 | 109 / 109 | b | 140% | < 0.0001 | | | |
| | wild type | 25°C | 25°C | 0.92 ± 0.02 | 0.89 | 1.06 | 137 / 137 | a | | | | | 2E |
| | AFD ablated | 25°C | 25°C | 1.29 ± 0.03 | 1.27 | 1.47 | 138 / 138 | b | 39% | < 0.0001 | | | |
| | wild type | 20°C | 20°C | 1.68 ± 0.03 | 1.61 | 1.80 | 122 / 122 | a | | | | | 2H |
| | AFD ablated | 20°C | 20°C | 3.13 ± 0.10 | 3.07 | 3.71 | 110 / 112 | b | 86% | < 0.0001 | | | |
| | wild type | 25°C | 20°C | 2.62 ± 0.06 | 2.53 | 2.97 | 120 / 120 | b | 55% | < 0.0001 | < 0.0001 | | |
| | AFD ablated | 25°C | 20°C | 2.58 ± 0.07 | 2.40 | 3.06 | 90 / 90 | b | 54% | < 0.0001 | < 0.0001 | > 0.05 | |
| Survival in 1 mM $H_2O_2$, *E. coli* JI377 | | | | | | | | | | | | | |
| | wild type | 20°C | 20°C | 0.78 ± 0.03 | 0.73 | 0.90 | 118 / 130 | a | | | | | S2A |
| | AFD ablated | 20°C | 20°C | 2.24 ± 0.09 | 2.02 | 2.76 | 91 / 114 | b | 185% | < 0.0001 | | | |

### Supplementary Table 3. Statistical analysis of gene-set expression for Figures 5, 6, S3, S4, S5, S7, and S8.

| | | wild type 25°C vs. wild type 20°C | | | | AFD(-) 20°C vs. wild type 20°C | | | |
|---|---|---|---|---|---|---|---|---|---|
| **Gene set** | **Gene set reference** | **Number of genes in set with expression** | **Fold change in gene expression log2(*wild type 25°C / wild type 20°C*) Mean ± SEM** | ***P* value vs. all genes (ANOVA)** | **Figure** | **Number of genes in set with expression** | **Fold change in gene expression log2(*AFD(-) 20°C / wild type 20°C]*) Mean ± SEM** | ***P* value vs. all genes (ANOVA)** | **Figure** |
| All genes | | 18039 | 0.04 ± 0.01 | | | 7912 | 0.10 ± 0.02 | | |
| Genes up at 25°C vs. 20°C | Gomez-Orte et al., 2018 | 173 | 0.54 ± 0.08 | < 0.0001 | S3B | 123 | 1.29 ± 0.15 | < 0.0001 | S3B |
| Genes up at 15°C vs. 20°C | Gomez-Orte et al., 2018 | 30 | -0.61 ± 0.17 | < 0.0001 | S3C | 30 | 2.23 ± 0.23 | < 0.0001 | S3C |
| Genes up at 30°C vs. 25°C | McCarroll et al., 2004 | 47 | -0.02 ± 0.10 | > 0.05 | S3D | 47 | 0.77 ± 0.18 | 0.0033 | S3D |
| Genes down after shift from 23°C to 17°C | Sugi et al., 2011 | 27 | 0.60 ± 0.25 | 0.016 | S3E | 14 | 2.56 ± 0.55 | < 0.0001 | S3E |
| Genes up after shift from 23°C to 17°C | Sugi et al., 2011 | 44 | -0.21 ± 0.15 | < 0.0001 | S3F | 26 | 1.59 ± 0.22 | < 0.0001 | S3F |
| Genes induced by 12 mM tBuOOH | Oliveira et al., 2009 | 129 | 0.51 ± 0.07 | < 0.0001 | S4B | 129 | 1.59 ± 0.09 | < 0.0001 | S4B |
| Genes induced by 1 g/L acrylamide | Lewis et al., 2009 | 268 | 0.08 ± 0.05 | > 0.05 | S5A | 167 | 0.69 ± 0.11 | < 0.0001 | S5A |
| Genes induced by formaldehyde | Yang et al., 2016 | 528 | 0.06 ± 0.04 | > 0.05 | S5B | 294 | 2.18 ± 0.09 | < 0.0001 | S5B |
| Genes induced by 1 mg/L benzene | Eom et al., 2014 | 64 | -0.08 ± 0.13 | > 0.05 | S5C | 49 | 2.89 ± 0.28 | < 0.0001 | S5C |
| Genes induced by 0.35 mg/L silver nanoparticles | Starnes et al., 2016 | 33 | -0.12 ± 0.13 | > 0.05 | S5D | 23 | 1.42 ± 0.23 | < 0.0001 | S5D |
| Genes induced by 1 mM cadmium chloride | Huffman et al., 2004 | 421 | 0.00 ± 0.05 | > 0.05 | S5E | 344 | 0.97 ± 0.07 | < 0.0001 | S5E |
| Genes induced by 2.3 mM sodium arsenite | Sahu et al., 2013 | 597 | -0.04 ± 0.03 | 0.0063 | S5F | 338 | 1.64 ± 0.06 | < 0.0001 | S5F |
| Genes induced by 60 mJ/cm2 310nm UVB light | Mueller et al., 2014 | 518 | 0.03 ± 0.04 | > 0.05 | S5G | 342 | 1.29 ± 0.07 | < 0.0001 | S5G |
| Genes induced by 120 Gy X-rays | Greiss et al., 2008 | 161 | -0.09 ± 0.08 | 0.015 | S5H | 133 | 1.73 ± 0.10 | < 0.0001 | S5H |
| Genes induced by 120 Gy gamma-rays | Greiss et al., 2008 | 134 | 0.01 ± 0.09 | > 0.05 | S5I | 101 | 1.75 ± 0.11 | < 0.0001 | S5I |
| Genes upregulated by *daf-16(+)* in wildtype | Kumar et al., 2015 | 37 | 0.55 ± 0.11 | < 0.0001 | 5A | 29 | 1.62 ± 0.22 | < 0.0001 | 5B |
| Genes upregulated by *daf-16(+)* in *daf-2(-)* | Murphy et al., 2003 | 216 | 0.27 ± 0.07 | < 0.0001 | S7A | 146 | 1.53 ± 0.10 | < 0.0001 | S7B |
| Genes upregulated by *skn-1(+)* in wildtype | Oliveira et al., 2009 | 285 | -0.11 ± 0.05 | 0.0002 | S7D | 196 | 1.68 ± 0.08 | < 0.0001 | 6A |
| Genes upregulated by *skn-1(+)* in *daf-2(-)* | Ewald et al., 2015 | 341 | 0.06 ± 0.04 | > 0.05 | S7E | 109 | 1.78 ± 0.10 | < 0.0001 | S7F |

| **Gene set** | **Gene set reference** | **Number of genes in set with expression** | **fold change in gene expression induced by *daf-16(mu86)* in AFD(+) nematodes at 20°C log2([*daf-16(-) effect*]) Mean ± SEM** | ***P* value vs. all genes (ANOVA)** | **Figure** |
|---|---|---|---|---|---|
| All genes | | 7387 | -0.04 ± 0.00 | | |
| Genes upregulated by *daf-16(+)* in wildtype | Kumar et al., 2015 | 27 | -0.21 ± 0.06 | 0.0017 | S8C |
| Genes upregulated by *daf-16(+)* in *daf-2(-)* | Murphy et al., 2003 | 132 | -0.16 ± 0.02 | < 0.0001 | S8D |

| **Gene set** | **Gene set reference** | **Number of genes in set with expression** | **additional fold change in gene expression induced by *daf-16(mu86)* in AFD(-) nematodes at 20°C log2([AFD(-) *daf-16(-)* interaction effect]) Mean ± SEM** | ***P* value vs. all genes (ANOVA)** | **Figure** |
|---|---|---|---|---|---|
| All genes | | 7387 | 0.02 ± 0.01 | | |
| Genes upregulated by *daf-16(+)* in *daf-2(-)* | Murphy et al., 2003 | 132 | -0.24 ± 0.06 | < 0.0001 | S8E |

### Supplementary Table 4A. Gene Ontology analysis.

| Dataset | Gene-set | Analysis | Term | Description | Expected | Observed | Enrichment Fold Change | P value | Q value |
|---|---|---|---|---|---|---|---|---|---|
| wildtype 25°C vs wildtype 20°C | 508 genes upregulated >2x (q=0.01) | Gene Ontology Enrichment Analysis | GO:0044419 | biological process involved in interspecies intera | 6.1 | 28 | 4.6 | 7.9E-12 | 2.4E-09 |
| wildtype 25°C vs wildtype 20°C | 508 genes upregulated >2x (q=0.01) | Gene Ontology Enrichment Analysis | GO:0009607 | response to biotic stimulus | 6.1 | 28 | 4.6 | 7.9E-12 | 2.4E-09 |
| wildtype 25°C vs wildtype 20°C | 508 genes upregulated >2x (q=0.01) | Gene Ontology Enrichment Analysis | GO:0002376 | immune system process | 6.2 | 28 | 4.5 | 1.2E-11 | 2.4E-09 |
| wildtype 25°C vs wildtype 20°C | 508 genes upregulated >2x (q=0.01) | Gene Ontology Enrichment Analysis | GO:0006952 | defense response | 6.1 | 27 | 4.4 | 4.8E-11 | 3.6E-09 |
| wildtype 25°C vs wildtype 20°C | 508 genes upregulated >2x (q=0.01) | Gene Ontology Enrichment Analysis | GO:0050830 | defense response to Gram-positive bacterium | 1.1 | 12 | 11 | 5E-11 | 3.6E-09 |
| wildtype 25°C vs wildtype 20°C | 508 genes upregulated >2x (q=0.01) | Gene Ontology Enrichment Analysis | GO:0005576 | extracellular region | 8.6 | 29 | 3.4 | 5.7E-09 | 0.00000028 |
| wildtype 25°C vs wildtype 20°C | 629 genes downregulated >2x (q=0.01) | Gene Ontology Enrichment Analysis | GO:0003735 | structural constituent of ribosome | 5 | 78 | 16 | 6E-76 | 1.8E-73 |
| wildtype 25°C vs wildtype 20°C | 629 genes downregulated >2x (q=0.01) | Gene Ontology Enrichment Analysis | GO:0022625 | cytosolic large ribosomal subunit | 1.7 | 43 | 25 | 3.5E-60 | 5.3E-58 |
| wildtype 25°C vs wildtype 20°C | 629 genes downregulated >2x (q=0.01) | Gene Ontology Enrichment Analysis | GO:0043043 | peptide biosynthetic process | 13 | 91 | 7.1 | 2.1E-51 | 2.1E-49 |
| wildtype 25°C vs wildtype 20°C | 629 genes downregulated >2x (q=0.01) | Gene Ontology Enrichment Analysis | GO:0016072 | rRNA metabolic process | 5.3 | 40 | 7.5 | 5.7E-25 | 4.3E-23 |
| wildtype 25°C vs wildtype 20°C | 629 genes downregulated >2x (q=0.01) | Gene Ontology Enrichment Analysis | GO:0032040 | small-subunit processome | 1.2 | 14 | 11 | 2.1E-13 | 1.3E-11 |
| wildtype 25°C vs wildtype 20°C | 629 genes downregulated >2x (q=0.01) | Gene Ontology Enrichment Analysis | GO:0071826 | ribonucleoprotein complex subunit organization | 3.8 | 20 | 5.3 | 1.6E-10 | 8.1E-09 |
| wildtype 25°C vs wildtype 20°C | 629 genes downregulated >2x (q=0.01) | Gene Ontology Enrichment Analysis | GO:0031974 | membrane-enclosed lumen | 27 | 58 | 2.2 | 1.6E-08 | 0.0000007 |
| wildtype 25°C vs wildtype 20°C | 629 genes downregulated >2x (q=0.01) | Gene Ontology Enrichment Analysis | GO:0030490 | maturation of SSU-rRNA | 1.4 | 10 | 7.3 | 6.4E-08 | 0.0000024 |
| wildtype 25°C vs wildtype 20°C | 629 genes downregulated >2x (q=0.01) | Gene Ontology Enrichment Analysis | GO:0000375 | RNA splicing via transesterification reactions | 5.4 | 20 | 3.7 | 0.00000012 | 0.000004 |
| wildtype 25°C vs wildtype 20°C | 629 genes downregulated >2x (q=0.01) | Gene Ontology Enrichment Analysis | GO:0003724 | RNA helicase activity | 1.8 | 11 | 6.2 | 0.00000013 | 0.000004 |
| wildtype 25°C vs wildtype 20°C | 629 genes downregulated >2x (q=0.01) | Gene Ontology Enrichment Analysis | GO:0009059 | macromolecule biosynthetic process | 66 | 110 | 1.7 | 0.00000015 | 0.0000042 |
| wildtype 25°C vs wildtype 20°C | 629 genes downregulated >2x (q=0.01) | Gene Ontology Enrichment Analysis | GO:0006725 | cellular aromatic compound metabolic process | 83 | 131 | 1.6 | 0.00000016 | 0.0000042 |
| wildtype 25°C vs wildtype 20°C | 629 genes downregulated >2x (q=0.01) | Gene Ontology Enrichment Analysis | GO:0043226 | organelle | 200 | 271 | 1.4 | 0.00000023 | 0.0000052 |
| wildtype 25°C vs wildtype 20°C | 629 genes downregulated >2x (q=0.01) | Gene Ontology Enrichment Analysis | GO:0008152 | metabolic process | 200 | 274 | 1.3 | 0.00000024 | 0.0000052 |
| wildtype 25°C vs wildtype 20°C | 629 genes downregulated >2x (q=0.01) | Gene Ontology Enrichment Analysis | GO:0046483 | heterocycle metabolic process | 83 | 130 | 1.6 | 0.00000024 | 0.0000052 |
| wildtype 25°C vs wildtype 20°C | 629 genes downregulated >2x (q=0.01) | Gene Ontology Enrichment Analysis | GO:1901360 | organic cyclic compound metabolic process | 85 | 131 | 1.5 | 0.00000046 | 0.0000087 |
| wildtype 25°C vs wildtype 20°C | 629 genes downregulated >2x (q=0.01) | Gene Ontology Enrichment Analysis | GO:0030016 | myofibril | 3.6 | 14 | 3.9 | 0.0000031 | 0.000055 |
| wildtype 25°C vs wildtype 20°C | 629 genes downregulated >2x (q=0.01) | Gene Ontology Enrichment Analysis | GO:0042302 | structural constituent of cuticle | 6.2 | 19 | 3.1 | 0.0000042 | 0.00007 |
| AFD-ablated 20°C vs wildtype 20°C | 2001 genes upregulated >2x (q=0.01) | Gene Ontology Enrichment Analysis | GO:0002376 | immune system process | 48 | 149 | 3.1 | 3E-39 | 9E-37 |
| AFD-ablated 20°C vs wildtype 20°C | 2001 genes upregulated >2x (q=0.01) | Gene Ontology Enrichment Analysis | GO:0009607 | response to biotic stimulus | 47 | 146 | 3.1 | 2.7E-38 | 4.1E-36 |
| AFD-ablated 20°C vs wildtype 20°C | 2001 genes upregulated >2x (q=0.01) | Gene Ontology Enrichment Analysis | GO:0044419 | biological process involved in interspecies intera | 47 | 146 | 3.1 | 2.7E-38 | 4.1E-36 |
| AFD-ablated 20°C vs wildtype 20°C | 2001 genes upregulated >2x (q=0.01) | Gene Ontology Enrichment Analysis | GO:0006952 | defense response | 47 | 146 | 3.1 | 7E-38 | 5.3E-36 |
| AFD-ablated 20°C vs wildtype 20°C | 2001 genes upregulated >2x (q=0.01) | Gene Ontology Enrichment Analysis | GO:0005576 | extracellular region | 66 | 174 | 2.6 | 2.1E-34 | 1.3E-32 |
| AFD-ablated 20°C vs wildtype 20°C | 2001 genes upregulated >2x (q=0.01) | Gene Ontology Enrichment Analysis | GO:0006082 | organic acid metabolic process | 50 | 134 | 2.7 | 4.2E-28 | 2.1E-26 |
| AFD-ablated 20°C vs wildtype 20°C | 2001 genes upregulated >2x (q=0.01) | Gene Ontology Enrichment Analysis | GO:0030016 | myofibril | 11 | 49 | 4.4 | 9E-22 | 3.9E-20 |
| AFD-ablated 20°C vs wildtype 20°C | 2001 genes upregulated >2x (q=0.01) | Gene Ontology Enrichment Analysis | GO:0098857 | membrane microdomain | 5.6 | 30 | 5.3 | 1.2E-17 | 4.5E-16 |
| AFD-ablated 20°C vs wildtype 20°C | 2001 genes upregulated >2x (q=0.01) | Gene Ontology Enrichment Analysis | GO:0031672 | A band | 4.3 | 24 | 5.6 | 2.2E-15 | 7.2E-14 |
| AFD-ablated 20°C vs wildtype 20°C | 2001 genes upregulated >2x (q=0.01) | Gene Ontology Enrichment Analysis | GO:0099081 | supramolecular polymer | 35 | 84 | 2.4 | 7.7E-15 | 2.3E-13 |
| AFD-ablated 20°C vs wildtype 20°C | 2001 genes upregulated >2x (q=0.01) | Gene Ontology Enrichment Analysis | GO:0000323 | lytic vacuole | 18 | 52 | 2.9 | 6.3E-14 | 1.7E-12 |
| AFD-ablated 20°C vs wildtype 20°C | 2001 genes upregulated >2x (q=0.01) | Gene Ontology Enrichment Analysis | GO:0016042 | lipid catabolic process | 17 | 49 | 2.8 | 1.7E-12 | 4.3E-11 |
| AFD-ablated 20°C vs wildtype 20°C | 2001 genes upregulated >2x (q=0.01) | Gene Ontology Enrichment Analysis | GO:0017171 | serine hydrolase activity | 15 | 43 | 2.9 | 2.3E-11 | 5.4E-10 |
| AFD-ablated 20°C vs wildtype 20°C | 2001 genes upregulated >2x (q=0.01) | Gene Ontology Enrichment Analysis | GO:0042579 | microbody | 8.2 | 29 | 3.5 | 6.3E-11 | 1.4E-09 |
| AFD-ablated 20°C vs wildtype 20°C | 2001 genes upregulated >2x (q=0.01) | Gene Ontology Enrichment Analysis | GO:0072329 | monocarboxylic acid catabolic process | 7.7 | 28 | 3.6 | 6.8E-11 | 1.4E-09 |
| AFD-ablated 20°C vs wildtype 20°C | 2001 genes upregulated >2x (q=0.01) | Gene Ontology Enrichment Analysis | GO:0003012 | muscle system process | 7.9 | 28 | 3.6 | 1E-10 | 1.9E-09 |
| AFD-ablated 20°C vs wildtype 20°C | 2001 genes upregulated >2x (q=0.01) | Gene Ontology Enrichment Analysis | GO:0005215 | transporter activity | 110 | 179 | 1.6 | 1.2E-10 | 2.2E-09 |
| AFD-ablated 20°C vs wildtype 20°C | 2001 genes upregulated >2x (q=0.01) | Gene Ontology Enrichment Analysis | GO:0003779 | actin binding | 14 | 39 | 2.8 | 2.6E-10 | 4.3E-09 |
| AFD-ablated 20°C vs wildtype 20°C | 2001 genes upregulated >2x (q=0.01) | Gene Ontology Enrichment Analysis | GO:0045177 | apical part of cell | 11 | 33 | 3.1 | 3.1E-10 | 4.9E-09 |
| AFD-ablated 20°C vs wildtype 20°C | 2001 genes upregulated >2x (q=0.01) | Gene Ontology Enrichment Analysis | GO:0055120 | striated muscle dense body | 11 | 32 | 3 | 1.8E-09 | 2.8E-08 |
| AFD-ablated 20°C vs wildtype 20°C | 2001 genes upregulated >2x (q=0.01) | Gene Ontology Enrichment Analysis | GO:0005506 | iron ion binding | 14 | 37 | 2.7 | 2.2E-09 | 3.1E-08 |
| AFD-ablated 20°C vs wildtype 20°C | 2001 genes upregulated >2x (q=0.01) | Gene Ontology Enrichment Analysis | GO:0055085 | transmembrane transport | 110 | 173 | 1.5 | 2.5E-09 | 3.4E-08 |
| AFD-ablated 20°C vs wildtype 20°C | 2001 genes upregulated >2x (q=0.01) | Gene Ontology Enrichment Analysis | GO:0050830 | defense response to Gram-positive bacterium | 8.2 | 26 | 3.2 | 8.8E-09 | 0.00000011 |
| AFD-ablated 20°C vs wildtype 20°C | 2001 genes upregulated >2x (q=0.01) | Gene Ontology Enrichment Analysis | GO:0034440 | lipid oxidation | 5.3 | 19 | 3.6 | 4.7E-08 | 0.00000059 |
| AFD-ablated 20°C vs wildtype 20°C | 2001 genes upregulated >2x (q=0.01) | Gene Ontology Enrichment Analysis | GO:0030312 | external encapsulating structure | 11 | 30 | 2.7 | 8.7E-08 | 0.000001 |
| AFD-ablated 20°C vs wildtype 20°C | 2001 genes upregulated >2x (q=0.01) | Gene Ontology Enrichment Analysis | GO:0140359 | ABC-type transporter activity | 6 | 20 | 3.4 | 9.7E-08 | 0.0000011 |
| AFD-ablated 20°C vs wildtype 20°C | 2001 genes upregulated >2x (q=0.01) | Gene Ontology Enrichment Analysis | GO:0050801 | ion homeostasis | 17 | 39 | 2.3 | 0.00000014 | 0.0000015 |
| AFD-ablated 20°C vs wildtype 20°C | 2001 genes upregulated >2x (q=0.01) | Gene Ontology Enrichment Analysis | GO:0030414 | peptidase inhibitor activity | 9.9 | 27 | 2.7 | 0.0000002 | 0.0000021 |
| AFD-ablated 20°C vs wildtype 20°C | 2001 genes upregulated >2x (q=0.01) | Gene Ontology Enrichment Analysis | GO:0072330 | monocarboxylic acid biosynthetic process | 6.2 | 20 | 3.2 | 0.0000002 | 0.0000021 |
| AFD-ablated 20°C vs wildtype 20°C | 2001 genes upregulated >2x (q=0.01) | Gene Ontology Enrichment Analysis | GO:0009925 | basal plasma membrane | 5.3 | 18 | 3.4 | 0.00000027 | 0.0000027 |
| AFD-ablated 20°C vs wildtype 20°C | 2001 genes upregulated >2x (q=0.01) | Gene Ontology Enrichment Analysis | GO:0045861 | negative regulation of proteolysis | 10 | 27 | 2.7 | 0.00000033 | 0.0000032 |
| AFD-ablated 20°C vs wildtype 20°C | 2001 genes upregulated >2x (q=0.01) | Gene Ontology Enrichment Analysis | GO:0061135 | endopeptidase regulator activity | 9.5 | 26 | 2.7 | 0.00000033 | 0.0000032 |
| AFD-ablated 20°C vs wildtype 20°C | 2001 genes upregulated >2x (q=0.01) | Gene Ontology Enrichment Analysis | GO:0009986 | cell surface | 12 | 29 | 2.5 | 0.00000063 | 0.0000057 |
| AFD-ablated 20°C vs wildtype 20°C | 2001 genes upregulated >2x (q=0.01) | Gene Ontology Enrichment Analysis | GO:0030029 | actin filament-based process | 21 | 44 | 2.1 | 0.0000011 | 0.0000096 |
| AFD-ablated 20°C vs wildtype 20°C | 2001 genes upregulated >2x (q=0.01) | Gene Ontology Enrichment Analysis | GO:0070279 | vitamin B6 binding | 4.7 | 16 | 3.4 | 0.0000011 | 0.0000096 |
| AFD-ablated 20°C vs wildtype 20°C | 2001 genes upregulated >2x (q=0.01) | Gene Ontology Enrichment Analysis | GO:0016614 | oxidoreductase activity acting on CH-OH group c | 8.6 | 23 | 2.7 | 0.0000021 | 0.000018 |
| AFD-ablated 20°C vs wildtype 20°C | 2001 genes upregulated >2x (q=0.01) | Gene Ontology Enrichment Analysis | GO:0022610 | biological adhesion | 12 | 29 | 2.3 | 0.0000028 | 0.000023 |
| AFD-ablated 20°C vs wildtype 20°C | 2001 genes upregulated >2x (q=0.01) | Gene Ontology Enrichment Analysis | GO:0010927 | cellular component assembly involved in morpho | 4 | 14 | 3.5 | 0.0000031 | 0.000024 |
| AFD-ablated 20°C vs wildtype 20°C | 1832 genes downregulated >2x (q=0.01) | Gene Ontology Enrichment Analysis | GO:0051321 | meiotic cell cycle | 34 | 129 | 3.8 | 2.2E-49 | 6.5E-47 |
| AFD-ablated 20°C vs wildtype 20°C | 1832 genes downregulated >2x (q=0.01) | Gene Ontology Enrichment Analysis | GO:0048285 | organelle fission | 32 | 126 | 3.9 | 4.2E-49 | 6.5E-47 |
| AFD-ablated 20°C vs wildtype 20°C | 1832 genes downregulated >2x (q=0.01) | Gene Ontology Enrichment Analysis | GO:0031974 | membrane-enclosed lumen | 110 | 244 | 2.3 | 3.6E-37 | 3.6E-35 |
| AFD-ablated 20°C vs wildtype 20°C | 1832 genes downregulated >2x (q=0.01) | Gene Ontology Enrichment Analysis | GO:0000003 | reproduction | 130 | 269 | 2.1 | 1.8E-35 | 1.3E-33 |
| AFD-ablated 20°C vs wildtype 20°C | 1832 genes downregulated >2x (q=0.01) | Gene Ontology Enrichment Analysis | GO:0006260 | DNA replication | 15 | 64 | 4.3 | 1.4E-29 | 8.2E-28 |
| AFD-ablated 20°C vs wildtype 20°C | 1832 genes downregulated >2x (q=0.01) | Gene Ontology Enrichment Analysis | GO:0006725 | cellular aromatic compound metabolic process | 340 | 524 | 1.6 | 2.1E-26 | 1.1E-24 |
| AFD-ablated 20°C vs wildtype 20°C | 1832 genes downregulated >2x (q=0.01) | Gene Ontology Enrichment Analysis | GO:0043226 | organelle | 810 | 1093 | 1.3 | 2.6E-26 | 1.1E-24 |
| AFD-ablated 20°C vs wildtype 20°C | 1832 genes downregulated >2x (q=0.01) | Gene Ontology Enrichment Analysis | GO:0046483 | heterocycle metabolic process | 340 | 523 | 1.6 | 2.8E-26 | 1.1E-24 |
| AFD-ablated 20°C vs wildtype 20°C | 1832 genes downregulated >2x (q=0.01) | Gene Ontology Enrichment Analysis | GO:1901360 | organic cyclic compound metabolic process | 340 | 525 | 1.5 | 1.5E-24 | 4.8E-23 |
| AFD-ablated 20°C vs wildtype 20°C | 1832 genes downregulated >2x (q=0.01) | Gene Ontology Enrichment Analysis | GO:0051704 | multi-organism process | 49 | 120 | 2.4 | 2.8E-22 | 8.3E-21 |
| AFD-ablated 20°C vs wildtype 20°C | 1832 genes downregulated >2x (q=0.01) | Gene Ontology Enrichment Analysis | GO:0000775 | chromosome centromeric region | 10 | 44 | 4.4 | 6.1E-22 | 1.7E-20 |
| AFD-ablated 20°C vs wildtype 20°C | 1832 genes downregulated >2x (q=0.01) | Gene Ontology Enrichment Analysis | GO:0009792 | embryo development ending in birth or egg hatch | 51 | 119 | 2.3 | 1.9E-20 | 4.7E-19 |
| AFD-ablated 20°C vs wildtype 20°C | 1832 genes downregulated >2x (q=0.01) | Gene Ontology Enrichment Analysis | GO:0000375 | RNA splicing via transesterification reactions | 22 | 66 | 3 | 1.1E-18 | 2.6E-17 |
| AFD-ablated 20°C vs wildtype 20°C | 1832 genes downregulated >2x (q=0.01) | Gene Ontology Enrichment Analysis | GO:0051306 | mitotic sister chromatid separation | 5 | 25 | 5 | 1.1E-15 | 2.3E-14 |
| AFD-ablated 20°C vs wildtype 20°C | 1832 genes downregulated >2x (q=0.01) | Gene Ontology Enrichment Analysis | GO:0000428 | DNA-directed RNA polymerase complex | 11 | 38 | 3.3 | 1.9E-13 | 3.8E-12 |
| AFD-ablated 20°C vs wildtype 20°C | 1832 genes downregulated >2x (q=0.01) | Gene Ontology Enrichment Analysis | GO:0045495 | pole plasm | 10 | 35 | 3.4 | 5.3E-13 | 9.8E-12 |
| AFD-ablated 20°C vs wildtype 20°C | 1832 genes downregulated >2x (q=0.01) | Gene Ontology Enrichment Analysis | GO:0035770 | ribonucleoprotein granule | 17 | 45 | 2.7 | 3E-11 | 5.2E-10 |
| AFD-ablated 20°C vs wildtype 20°C | 1832 genes downregulated >2x (q=0.01) | Gene Ontology Enrichment Analysis | GO:0003697 | single-stranded DNA binding | 5.6 | 22 | 4 | 1E-10 | 1.7E-09 |
| AFD-ablated 20°C vs wildtype 20°C | 1832 genes downregulated >2x (q=0.01) | Gene Ontology Enrichment Analysis | GO:0009994 | oocyte differentiation | 8.1 | 27 | 3.4 | 2.7E-10 | 4.2E-09 |
| AFD-ablated 20°C vs wildtype 20°C | 1832 genes downregulated >2x (q=0.01) | Gene Ontology Enrichment Analysis | GO:0043632 | modification-dependent macromolecule catabolic | 38 | 76 | 2 | 4.1E-10 | 6.1E-09 |
| AFD-ablated 20°C vs wildtype 20°C | 1832 genes downregulated >2x (q=0.01) | Gene Ontology Enrichment Analysis | GO:0016569 | covalent chromatin modification | 23 | 53 | 2.3 | 5.2E-10 | 7.4E-09 |
| AFD-ablated 20°C vs wildtype 20°C | 1832 genes downregulated >2x (q=0.01) | Gene Ontology Enrichment Analysis | GO:0000725 | recombinational repair | 7.8 | 26 | 3.4 | 5.5E-10 | 7.5E-09 |
| AFD-ablated 20°C vs wildtype 20°C | 1832 genes downregulated >2x (q=0.01) | Gene Ontology Enrichment Analysis | GO:0016887 | ATP hydrolysis activity | 24 | 54 | 2.2 | 1E-09 | 1.4E-08 |
| AFD-ablated 20°C vs wildtype 20°C | 1832 genes downregulated >2x (q=0.01) | Gene Ontology Enrichment Analysis | GO:0070727 | cellular macromolecule localization | 72 | 122 | 1.7 | 1.4E-09 | 1.7E-08 |
| AFD-ablated 20°C vs wildtype 20°C | 1832 genes downregulated >2x (q=0.01) | Gene Ontology Enrichment Analysis | GO:0071695 | anatomical structure maturation | 6.7 | 23 | 3.4 | 2.6E-09 | 3.2E-08 |
| AFD-ablated 20°C vs wildtype 20°C | 1832 genes downregulated >2x (q=0.01) | Gene Ontology Enrichment Analysis | GO:0050684 | regulation of mRNA processing | 6.7 | 23 | 3.4 | 2.6E-09 | 3.2E-08 |
| AFD-ablated 20°C vs wildtype 20°C | 1832 genes downregulated >2x (q=0.01) | Gene Ontology Enrichment Analysis | GO:0000132 | establishment of mitotic spindle orientation | 7 | 23 | 3.3 | 8E-09 | 8.9E-08 |
| AFD-ablated 20°C vs wildtype 20°C | 1832 genes downregulated >2x (q=0.01) | Gene Ontology Enrichment Analysis | GO:0051653 | spindle localization | 11 | 31 | 2.7 | 9.4E-09 | 0.0000001 |
| AFD-ablated 20°C vs wildtype 20°C | 1832 genes downregulated >2x (q=0.01) | Gene Ontology Enrichment Analysis | GO:0045137 | development of primary sexual characteristics | 17 | 40 | 2.4 | 1.8E-08 | 0.00000019 |
| AFD-ablated 20°C vs wildtype 20°C | 1832 genes downregulated >2x (q=0.01) | Gene Ontology Enrichment Analysis | GO:0035825 | homologous recombination | 6.3 | 21 | 3.3 | 1.9E-08 | 0.00000019 |
| AFD-ablated 20°C vs wildtype 20°C | 1832 genes downregulated >2x (q=0.01) | Gene Ontology Enrichment Analysis | GO:0051028 | mRNA transport | 6 | 20 | 3.3 | 0.00000004 | 0.00000038 |
| AFD-ablated 20°C vs wildtype 20°C | 1832 genes downregulated >2x (q=0.01) | Gene Ontology Enrichment Analysis | GO:0051169 | nuclear transport | 13 | 32 | 2.5 | 7.9E-08 | 0.00000074 |
| AFD-ablated 20°C vs wildtype 20°C | 1832 genes downregulated >2x (q=0.01) | Gene Ontology Enrichment Analysis | GO:0050657 | nucleic acid transport | 8.8 | 25 | 2.8 | 8.1E-08 | 0.00000074 |
| AFD-ablated 20°C vs wildtype 20°C | 1832 genes downregulated >2x (q=0.01) | Gene Ontology Enrichment Analysis | GO:0003730 | mRNA 3'-UTR binding | 6.9 | 21 | 3.1 | 0.00000015 | 0.0000013 |
| AFD-ablated 20°C vs wildtype 20°C | 1832 genes downregulated >2x (q=0.01) | Gene Ontology Enrichment Analysis | GO:0031023 | microtubule organizing center organization | 7.5 | 22 | 2.9 | 0.00000018 | 0.0000016 |
| AFD-ablated 20°C vs wildtype 20°C | 1832 genes downregulated >2x (q=0.01) | Gene Ontology Enrichment Analysis | GO:0018394 | peptidyl-lysine acetylation | 6.6 | 20 | 3 | 0.00000031 | 0.0000026 |
| AFD-ablated 20°C vs wildtype 20°C | 1832 genes downregulated >2x (q=0.01) | Gene Ontology Enrichment Analysis | GO:0006475 | internal protein amino acid acetylation | 6.6 | 20 | 3 | 0.00000031 | 0.0000026 |
| AFD-ablated 20°C vs wildtype 20°C | 1832 genes downregulated >2x (q=0.01) | Gene Ontology Enrichment Analysis | GO:0044257 | cellular protein catabolic process | 47 | 81 | 1.7 | 0.00000033 | 0.0000026 |
| AFD-ablated 20°C vs wildtype 20°C | 1832 genes downregulated >2x (q=0.01) | Gene Ontology Enrichment Analysis | GO:0048566 | embryonic digestive tract development | 4.8 | 16 | 3.3 | 0.00000071 | 0.0000054 |
| AFD-ablated 20°C vs wildtype 20°C | 1832 genes downregulated >2x (q=0.01) | Gene Ontology Enrichment Analysis | GO:0009059 | macromolecule biosynthetic process | 270 | 344 | 1.3 | 0.00000073 | 0.0000055 |
| AFD-ablated 20°C vs wildtype 20°C | 1832 genes downregulated >2x (q=0.01) | Gene Ontology Enrichment Analysis | GO:0009952 | anterior/posterior pattern specification | 9.7 | 25 | 2.6 | 0.00000078 | 0.0000057 |
| AFD-ablated 20°C vs wildtype 20°C | 1832 genes downregulated >2x (q=0.01) | Gene Ontology Enrichment Analysis | GO:0051247 | positive regulation of protein metabolic process | 28 | 52 | 1.9 | 0.0000021 | 0.000015 |
| AFD-ablated 20°C vs wildtype 20°C | 1832 genes downregulated >2x (q=0.01) | Gene Ontology Enrichment Analysis | GO:0051170 | import into nucleus | 6.4 | 18 | 2.8 | 0.0000047 | 0.000033 |
| AFD-ablated 20°C vs wildtype 20°C | 1832 genes downregulated >2x (q=0.01) | Gene Ontology Enrichment Analysis | GO:0061458 | reproductive system development | 23 | 43 | 1.9 | 0.0000077 | 0.000052 |
| AFD-ablated 20°C vs wildtype 20°C | 1832 genes downregulated >2x (q=0.01) | Gene Ontology Enrichment Analysis | GO:0048569 | post-embryonic animal organ development | 26 | 48 | 1.8 | 0.0000087 | 0.000058 |

### Supplementary Table 4B. Tissue Enrichment analysis.

| Dataset | Gene-set | Analysis | Term | Description | Expected | Observed | Enrichment Fold Change | P value | Q value |
|---|---|---|---|---|---|---|---|---|---|
| wildtype 25°C vs wildtype 20°C | 508 genes upregulated >2x (q=0.01) | Tissue Enrichment Analysis | WBbt:0005730 | epithelial system | 110 | 158 | 1.5 | 0.0000011 | 0.00034 |
| wildtype 25°C vs wildtype 20°C | 508 genes upregulated >2x (q=0.01) | Tissue Enrichment Analysis | WBbt:0005772 | intestine | 150 | 190 | 1.3 | 0.00011 | 0.017 |
| wildtype 25°C vs wildtype 20°C | 508 genes upregulated >2x (q=0.01) | Tissue Enrichment Analysis | WBbt:0004697 | head mesodermal cell | 66 | 97 | 1.5 | 0.00012 | 0.017 |
| wildtype 25°C vs wildtype 20°C | 629 genes downregulated >2x (q=0.01) | Tissue Enrichment Analysis | WBbt:0005784 | germ line | 230 | 395 | 1.7 | 2.3E-25 | 7.2E-23 |
| wildtype 25°C vs wildtype 20°C | 629 genes downregulated >2x (q=0.01) | Tissue Enrichment Analysis | WBbt:0005747 | reproductive system | 280 | 421 | 1.5 | 3E-17 | 4.7E-15 |
| wildtype 25°C vs wildtype 20°C | 629 genes downregulated >2x (q=0.01) | Tissue Enrichment Analysis | WBbt:0005779 | striated muscle | 48 | 111 | 2.3 | 4.6E-16 | 4.8E-14 |
| wildtype 25°C vs wildtype 20°C | 629 genes downregulated >2x (q=0.01) | Tissue Enrichment Analysis | WBbt:0006874 | Psub1 | 8 | 30 | 3.7 | 1.6E-10 | 1.3E-08 |
| wildtype 25°C vs wildtype 20°C | 629 genes downregulated >2x (q=0.01) | Tissue Enrichment Analysis | WBbt:0008366 | gonadal primordium | 98 | 162 | 1.7 | 2.7E-10 | 1.7E-08 |
| wildtype 25°C vs wildtype 20°C | 629 genes downregulated >2x (q=0.01) | Tissue Enrichment Analysis | WBbt:0005772 | intestine | 230 | 322 | 1.4 | 2.2E-09 | 0.00000011 |
| wildtype 25°C vs wildtype 20°C | 629 genes downregulated >2x (q=0.01) | Tissue Enrichment Analysis | WBbt:0004292 | anal depressor muscle | 16 | 35 | 2.2 | 0.0000064 | 0.00028 |
| wildtype 25°C vs wildtype 20°C | 629 genes downregulated >2x (q=0.01) | Tissue Enrichment Analysis | WBbt:0005842 | AVA | 28 | 51 | 1.8 | 0.000019 | 0.00075 |
| wildtype 25°C vs wildtype 20°C | 629 genes downregulated >2x (q=0.01) | Tissue Enrichment Analysis | WBbt:0005737 | muscular system | 230 | 292 | 1.3 | 0.000039 | 0.0013 |
| wildtype 25°C vs wildtype 20°C | 629 genes downregulated >2x (q=0.01) | Tissue Enrichment Analysis | WBbt:0008406 | cephalic sheath cell | 24 | 45 | 1.8 | 0.000046 | 0.0014 |
| wildtype 25°C vs wildtype 20°C | 629 genes downregulated >2x (q=0.01) | Tissue Enrichment Analysis | WBbt:0007849 | hermaphrodite | 77 | 105 | 1.4 | 0.00062 | 0.018 |
| AFD-ablated 20°C vs wildtype 20°C | 2001 genes upregulated >2x (q=0.01) | Tissue Enrichment Analysis | WBbt:0005772 | intestine | 870 | 1394 | 1.6 | 3.1E-74 | 9.9E-72 |
| AFD-ablated 20°C vs wildtype 20°C | 2001 genes upregulated >2x (q=0.01) | Tissue Enrichment Analysis | WBbt:0004697 | head mesodermal cell | 400 | 712 | 1.8 | 8.1E-56 | 1.3E-53 |
| AFD-ablated 20°C vs wildtype 20°C | 2001 genes upregulated >2x (q=0.01) | Tissue Enrichment Analysis | WBbt:0005737 | muscular system | 870 | 1213 | 1.4 | 7.5E-34 | 7.8E-32 |
| AFD-ablated 20°C vs wildtype 20°C | 2001 genes upregulated >2x (q=0.01) | Tissue Enrichment Analysis | WBbt:0006831 | PVD | 360 | 537 | 1.5 | 1.1E-21 | 8.9E-20 |
| AFD-ablated 20°C vs wildtype 20°C | 2001 genes upregulated >2x (q=0.01) | Tissue Enrichment Analysis | WBbt:0005501 | outer labial sensillum | 370 | 543 | 1.5 | 3.9E-21 | 2.4E-19 |
| AFD-ablated 20°C vs wildtype 20°C | 2001 genes upregulated >2x (q=0.01) | Tissue Enrichment Analysis | WBbt:0008422 | sex organ | 220 | 311 | 1.4 | 1E-11 | 5.2E-10 |
| AFD-ablated 20°C vs wildtype 20°C | 2001 genes upregulated >2x (q=0.01) | Tissue Enrichment Analysis | WBbt:0007849 | hermaphrodite | 290 | 381 | 1.3 | 4.5E-09 | 0.0000002 |
| AFD-ablated 20°C vs wildtype 20°C | 2001 genes upregulated >2x (q=0.01) | Tissue Enrichment Analysis | WBbt:0005736 | excretory system | 120 | 183 | 1.5 | 5.8E-09 | 0.00000023 |
| AFD-ablated 20°C vs wildtype 20°C | 2001 genes upregulated >2x (q=0.01) | Tissue Enrichment Analysis | WBbt:0006850 | excretory secretory system | 120 | 183 | 1.5 | 6.3E-09 | 0.00000023 |
| AFD-ablated 20°C vs wildtype 20°C | 2001 genes upregulated >2x (q=0.01) | Tissue Enrichment Analysis | WBbt:0005812 | excretory cell | 110 | 165 | 1.5 | 1.7E-08 | 0.00000052 |
| AFD-ablated 20°C vs wildtype 20°C | 2001 genes upregulated >2x (q=0.01) | Tissue Enrichment Analysis | WBbt:0008406 | cephalic sheath cell | 92 | 136 | 1.5 | 0.0000012 | 0.000035 |
| AFD-ablated 20°C vs wildtype 20°C | 2001 genes upregulated >2x (q=0.01) | Tissue Enrichment Analysis | WBbt:0005798 | anal sphincter muscle | 16 | 36 | 2.2 | 0.0000013 | 0.000035 |
| AFD-ablated 20°C vs wildtype 20°C | 2001 genes upregulated >2x (q=0.01) | Tissue Enrichment Analysis | WBbt:0004292 | anal depressor muscle | 60 | 93 | 1.6 | 0.0000054 | 0.00013 |
| AFD-ablated 20°C vs wildtype 20°C | 2001 genes upregulated >2x (q=0.01) | Tissue Enrichment Analysis | WBbt:0005237 | touch receptor neuron | 250 | 305 | 1.2 | 0.000045 | 0.001 |
| AFD-ablated 20°C vs wildtype 20°C | 2001 genes upregulated >2x (q=0.01) | Tissue Enrichment Analysis | WBbt:0005749 | coelomic system | 110 | 152 | 1.3 | 0.00013 | 0.0027 |
| AFD-ablated 20°C vs wildtype 20°C | 2001 genes upregulated >2x (q=0.01) | Tissue Enrichment Analysis | WBbt:0005785 | somatic gonad | 72 | 102 | 1.4 | 0.00014 | 0.0027 |
| AFD-ablated 20°C vs wildtype 20°C | 2001 genes upregulated >2x (q=0.01) | Tissue Enrichment Analysis | WBbt:0003681 | pharynx | 650 | 740 | 1.1 | 0.00014 | 0.0027 |
| AFD-ablated 20°C vs wildtype 20°C | 2001 genes upregulated >2x (q=0.01) | Tissue Enrichment Analysis | WBbt:0006791 | uv1 | 6 | 15 | 2.5 | 0.00014 | 0.0027 |
| AFD-ablated 20°C vs wildtype 20°C | 2001 genes upregulated >2x (q=0.01) | Tissue Enrichment Analysis | WBbt:0006769 | lateral nerve cord | 52 | 76 | 1.5 | 0.00021 | 0.0035 |
| AFD-ablated 20°C vs wildtype 20°C | 2001 genes upregulated >2x (q=0.01) | Tissue Enrichment Analysis | WBbt:0005342 | uterine muscle | 17 | 31 | 1.8 | 0.00026 | 0.0041 |
| AFD-ablated 20°C vs wildtype 20°C | 2001 genes upregulated >2x (q=0.01) | Tissue Enrichment Analysis | WBbt:0005730 | epithelial system | 650 | 723 | 1.1 | 0.00045 | 0.0067 |
| AFD-ablated 20°C vs wildtype 20°C | 2001 genes upregulated >2x (q=0.01) | Tissue Enrichment Analysis | WBbt:0006789 | uterine seam cell | 5.5 | 13 | 2.4 | 0.00063 | 0.009 |
| AFD-ablated 20°C vs wildtype 20°C | 2001 genes upregulated >2x (q=0.01) | Tissue Enrichment Analysis | WBbt:0006104 | Eala | 5 | 12 | 2.4 | 0.00078 | 0.011 |
| AFD-ablated 20°C vs wildtype 20°C | 2001 genes upregulated >2x (q=0.01) | Tissue Enrichment Analysis | WBbt:0005319 | spermatheca | 74 | 100 | 1.3 | 0.00095 | 0.012 |
| AFD-ablated 20°C vs wildtype 20°C | 2001 genes upregulated >2x (q=0.01) | Tissue Enrichment Analysis | WBbt:0006749 | nerve ring | 110 | 143 | 1.3 | 0.0013 | 0.016 |
| AFD-ablated 20°C vs wildtype 20°C | 2001 genes upregulated >2x (q=0.01) | Tissue Enrichment Analysis | WBbt:0006546 | Ealp | 5.2 | 12 | 2.3 | 0.0013 | 0.016 |
| AFD-ablated 20°C vs wildtype 20°C | 2001 genes upregulated >2x (q=0.01) | Tissue Enrichment Analysis | WBbt:0006750 | dorsal nerve cord | 76 | 100 | 1.3 | 0.0016 | 0.018 |
| AFD-ablated 20°C vs wildtype 20°C | 2001 genes upregulated >2x (q=0.01) | Tissue Enrichment Analysis | WBbt:0006161 | Eara | 5.4 | 12 | 2.2 | 0.0017 | 0.019 |
| AFD-ablated 20°C vs wildtype 20°C | 2001 genes upregulated >2x (q=0.01) | Tissue Enrichment Analysis | WBbt:0006646 | Earp | 5.4 | 12 | 2.2 | 0.0017 | 0.019 |
| AFD-ablated 20°C vs wildtype 20°C | 1832 genes downregulated >2x (q=0.01) | Tissue Enrichment Analysis | WBbt:0005784 | germ line | 860 | 1809 | 2.1 | 1.8E-224 | 5.5E-222 |
| AFD-ablated 20°C vs wildtype 20°C | 1832 genes downregulated >2x (q=0.01) | Tissue Enrichment Analysis | WBbt:0005747 | reproductive system | 1000 | 1813 | 1.7 | 1.2E-133 | 2E-131 |
| AFD-ablated 20°C vs wildtype 20°C | 1832 genes downregulated >2x (q=0.01) | Tissue Enrichment Analysis | WBbt:0004015 | AB | 15 | 59 | 4 | 1.1E-23 | 1.1E-21 |
| AFD-ablated 20°C vs wildtype 20°C | 1832 genes downregulated >2x (q=0.01) | Tissue Enrichment Analysis | WBbt:0006876 | EMS | 8.9 | 43 | 4.8 | 1.9E-22 | 1.5E-20 |
| AFD-ablated 20°C vs wildtype 20°C | 1832 genes downregulated >2x (q=0.01) | Tissue Enrichment Analysis | WBbt:0006875 | Psub3 | 8.4 | 41 | 4.9 | 9.7E-22 | 6.1E-20 |
| AFD-ablated 20°C vs wildtype 20°C | 1832 genes downregulated >2x (q=0.01) | Tissue Enrichment Analysis | WBbt:0006874 | Psub1 | 30 | 85 | 2.9 | 8.9E-21 | 4.6E-19 |
| AFD-ablated 20°C vs wildtype 20°C | 1832 genes downregulated >2x (q=0.01) | Tissue Enrichment Analysis | WBbt:0004575 | Z3 | 6.8 | 34 | 5 | 5.7E-19 | 2.6E-17 |
| AFD-ablated 20°C vs wildtype 20°C | 1832 genes downregulated >2x (q=0.01) | Tissue Enrichment Analysis | WBbt:0004576 | Z2 | 6.8 | 34 | 5 | 5.7E-19 | 2.6E-17 |
| AFD-ablated 20°C vs wildtype 20°C | 1832 genes downregulated >2x (q=0.01) | Tissue Enrichment Analysis | WBbt:0006797 | oocyte | 21 | 65 | 3.1 | 2.4E-18 | 8.5E-17 |
| AFD-ablated 20°C vs wildtype 20°C | 1832 genes downregulated >2x (q=0.01) | Tissue Enrichment Analysis | WBbt:0006873 | Psub2 | 6 | 29 | 4.8 | 8.7E-16 | 2.7E-14 |
| AFD-ablated 20°C vs wildtype 20°C | 1832 genes downregulated >2x (q=0.01) | Tissue Enrichment Analysis | WBbt:0003810 | C | 5.4 | 22 | 4.1 | 1.7E-10 | 5E-09 |
| AFD-ablated 20°C vs wildtype 20°C | 1832 genes downregulated >2x (q=0.01) | Tissue Enrichment Analysis | WBbt:0005744 | reproductive tract | 240 | 308 | 1.3 | 0.00000035 | 0.0000092 |
| AFD-ablated 20°C vs wildtype 20°C | 1832 genes downregulated >2x (q=0.01) | Tissue Enrichment Analysis | WBbt:0006892 | P4.p | 7.8 | 20 | 2.6 | 0.000012 | 0.00029 |
| AFD-ablated 20°C vs wildtype 20°C | 1832 genes downregulated >2x (q=0.01) | Tissue Enrichment Analysis | WBbt:0006891 | P3.p | 7.8 | 20 | 2.6 | 0.000012 | 0.00029 |
| AFD-ablated 20°C vs wildtype 20°C | 1832 genes downregulated >2x (q=0.01) | Tissue Enrichment Analysis | WBbt:0006896 | P8.p | 7.9 | 20 | 2.5 | 0.000016 | 0.00033 |
| AFD-ablated 20°C vs wildtype 20°C | 1832 genes downregulated >2x (q=0.01) | Tissue Enrichment Analysis | WBbt:0006894 | P6.p | 8.9 | 21 | 2.4 | 0.000035 | 0.00068 |
| AFD-ablated 20°C vs wildtype 20°C | 1832 genes downregulated >2x (q=0.01) | Tissue Enrichment Analysis | WBbt:0006893 | P5.p | 8.9 | 20 | 2.3 | 0.00011 | 0.002 |
| AFD-ablated 20°C vs wildtype 20°C | 1832 genes downregulated >2x (q=0.01) | Tissue Enrichment Analysis | WBbt:0006895 | P7.p | 9 | 20 | 2.2 | 0.00014 | 0.0024 |
| AFD-ablated 20°C vs wildtype 20°C | 1832 genes downregulated >2x (q=0.01) | Tissue Enrichment Analysis | WBbt:0008378 | somatic cell | 8.4 | 18 | 2.1 | 0.00042 | 0.0069 |
| AFD-ablated 20°C vs wildtype 20°C | 1832 genes downregulated >2x (q=0.01) | Tissue Enrichment Analysis | WBbt:0008406 | cephalic sheath cell | 91 | 120 | 1.3 | 0.00069 | 0.011 |
| AFD-ablated 20°C vs wildtype 20°C | 1832 genes downregulated >2x (q=0.01) | Tissue Enrichment Analysis | WBbt:0006201 | MSpapp | 4.4 | 11 | 2.5 | 0.00087 | 0.013 |

### Supplementary Table 4C. Phenotype Enrichment analysis.

| Dataset | Gene-set | Analysis | Term | Description | Expected | Observed | Enrichment Fold Change | P value | Q value |
|---|---|---|---|---|---|---|---|---|---|
| wildtype 25°C vs wildtype 20°C | 508 genes upregulated >2x (q=0.01) | Phenotype Enrichment Analysis | WBPhenotype:0000886 | nematode phenotype | 95 | 149 | 1.6 | 1.4E-08 | 0.0000034 |
| wildtype 25°C vs wildtype 20°C | 508 genes upregulated >2x (q=0.01) | Phenotype Enrichment Analysis | WBPhenotype:0001655 | cadmium hypersensitive | 0.88 | 7 | 7.9 | 0.0000032 | 0.00039 |
| wildtype 25°C vs wildtype 20°C | 508 genes upregulated >2x (q=0.01) | Phenotype Enrichment Analysis | WBPhenotype:0002539 | antimicrobial gene expression increased | 0.49 | 4 | 8.2 | 0.00013 | 0.01 |
| wildtype 25°C vs wildtype 20°C | 629 genes downregulated >2x (q=0.01) | Phenotype Enrichment Analysis | WBPhenotype:0000270 | pleiotropic defects severe early emb | 8.4 | 65 | 7.8 | 6.8E-43 | 1.6E-40 |
| wildtype 25°C vs wildtype 20°C | 629 genes downregulated >2x (q=0.01) | Phenotype Enrichment Analysis | WBPhenotype:0001941 | rachis narrow | 18 | 66 | 3.7 | 3.4E-21 | 4E-19 |
| wildtype 25°C vs wildtype 20°C | 629 genes downregulated >2x (q=0.01) | Phenotype Enrichment Analysis | WBPhenotype:0001946 | pachytene progression during oogenesis variant | 18 | 65 | 3.6 | 4.9E-20 | 3.9E-18 |
| wildtype 25°C vs wildtype 20°C | 629 genes downregulated >2x (q=0.01) | Phenotype Enrichment Analysis | WBPhenotype:0001979 | gonad vesiculated | 13 | 53 | 4 | 5.7E-19 | 3.4E-17 |
| wildtype 25°C vs wildtype 20°C | 629 genes downregulated >2x (q=0.01) | Phenotype Enrichment Analysis | WBPhenotype:0001944 | oocyte number decreased | 22 | 67 | 3.1 | 7.1E-17 | 3.4E-15 |
| wildtype 25°C vs wildtype 20°C | 629 genes downregulated >2x (q=0.01) | Phenotype Enrichment Analysis | WBPhenotype:0001792 | nuclei small | 11 | 43 | 4 | 4.2E-16 | 1.7E-14 |
| wildtype 25°C vs wildtype 20°C | 629 genes downregulated >2x (q=0.01) | Phenotype Enrichment Analysis | WBPhenotype:0001954 | diplotene absent during oogenesis | 7.1 | 34 | 4.8 | 1.6E-15 | 5.4E-14 |
| wildtype 25°C vs wildtype 20°C | 629 genes downregulated >2x (q=0.01) | Phenotype Enrichment Analysis | WBPhenotype:0001980 | germ cell compartment expansion variant | 27 | 73 | 2.7 | 2.3E-15 | 6.7E-14 |
| wildtype 25°C vs wildtype 20°C | 629 genes downregulated >2x (q=0.01) | Phenotype Enrichment Analysis | WBPhenotype:0001950 | diplotene region organization variant | 13 | 43 | 3.4 | 2.3E-13 | 6.1E-12 |
| wildtype 25°C vs wildtype 20°C | 629 genes downregulated >2x (q=0.01) | Phenotype Enrichment Analysis | WBPhenotype:0001827 | RNA metabolism variant | 10 | 37 | 3.6 | 1.7E-12 | 4.2E-11 |
| wildtype 25°C vs wildtype 20°C | 629 genes downregulated >2x (q=0.01) | Phenotype Enrichment Analysis | WBPhenotype:0001957 | gonad small | 5.6 | 26 | 4.6 | 4.2E-12 | 9.2E-11 |
| wildtype 25°C vs wildtype 20°C | 629 genes downregulated >2x (q=0.01) | Phenotype Enrichment Analysis | WBPhenotype:0001538 | mRNA surveillance defective | 7.1 | 29 | 4.1 | 1E-11 | 2E-10 |
| wildtype 25°C vs wildtype 20°C | 629 genes downregulated >2x (q=0.01) | Phenotype Enrichment Analysis | WBPhenotype:0000508 | nonsense mRNA accumulation | 6.9 | 28 | 4.1 | 2.7E-11 | 5E-10 |
| wildtype 25°C vs wildtype 20°C | 629 genes downregulated >2x (q=0.01) | Phenotype Enrichment Analysis | WBPhenotype:0000402 | avoids bacterial lawn | 28 | 66 | 2.3 | 7.5E-11 | 1.3E-09 |
| wildtype 25°C vs wildtype 20°C | 629 genes downregulated >2x (q=0.01) | Phenotype Enrichment Analysis | WBPhenotype:0001794 | cytoplasmic processing body variant | 16 | 45 | 2.8 | 1.5E-10 | 2.5E-09 |
| wildtype 25°C vs wildtype 20°C | 629 genes downregulated >2x (q=0.01) | Phenotype Enrichment Analysis | WBPhenotype:0001259 | hermaphrodite fertility reduced | 65 | 117 | 1.8 | 3.9E-10 | 5.9E-09 |
| wildtype 25°C vs wildtype 20°C | 629 genes downregulated >2x (q=0.01) | Phenotype Enrichment Analysis | WBPhenotype:0001942 | rachis absent | 5.4 | 22 | 4.1 | 2.5E-09 | 3.6E-08 |
| wildtype 25°C vs wildtype 20°C | 629 genes downregulated >2x (q=0.01) | Phenotype Enrichment Analysis | WBPhenotype:0000184 | apoptosis fails to occur | 5.9 | 23 | 3.9 | 3E-09 | 3.9E-08 |
| wildtype 25°C vs wildtype 20°C | 629 genes downregulated >2x (q=0.01) | Phenotype Enrichment Analysis | WBPhenotype:0000676 | growth rate variant | 120 | 168 | 1.4 | 0.0000022 | 0.000027 |
| wildtype 25°C vs wildtype 20°C | 629 genes downregulated >2x (q=0.01) | Phenotype Enrichment Analysis | WBPhenotype:0000774 | gametogenesis variant | 81 | 120 | 1.5 | 0.0000088 | 0.0001 |
| wildtype 25°C vs wildtype 20°C | 629 genes downregulated >2x (q=0.01) | Phenotype Enrichment Analysis | WBPhenotype:0001015 | developmental growth variant | 120 | 161 | 1.4 | 0.00001 | 0.00012 |
| wildtype 25°C vs wildtype 20°C | 629 genes downregulated >2x (q=0.01) | Phenotype Enrichment Analysis | WBPhenotype:0000033 | developmental timing variant | 130 | 176 | 1.3 | 0.000073 | 0.00079 |
| wildtype 25°C vs wildtype 20°C | 629 genes downregulated >2x (q=0.01) | Phenotype Enrichment Analysis | WBPhenotype:0001945 | oocytes small | 8 | 19 | 2.4 | 0.00014 | 0.0014 |
| wildtype 25°C vs wildtype 20°C | 629 genes downregulated >2x (q=0.01) | Phenotype Enrichment Analysis | WBPhenotype:0000032 | sick | 41 | 64 | 1.5 | 0.00025 | 0.0025 |
| wildtype 25°C vs wildtype 20°C | 629 genes downregulated >2x (q=0.01) | Phenotype Enrichment Analysis | WBPhenotype:0001982 | cell membrane organization biogenesis variant | 20 | 35 | 1.7 | 0.00058 | 0.0056 |
| wildtype 25°C vs wildtype 20°C | 629 genes downregulated >2x (q=0.01) | Phenotype Enrichment Analysis | WBPhenotype:0002041 | molt variant | 17 | 31 | 1.8 | 0.00066 | 0.0061 |
| wildtype 25°C vs wildtype 20°C | 629 genes downregulated >2x (q=0.01) | Phenotype Enrichment Analysis | WBPhenotype:0001029 | patchy coloration | 6.5 | 15 | 2.3 | 0.0007 | 0.0062 |
| wildtype 25°C vs wildtype 20°C | 629 genes downregulated >2x (q=0.01) | Phenotype Enrichment Analysis | WBPhenotype:0001925 | oocytes disorganized | 4.5 | 11 | 2.5 | 0.0015 | 0.013 |
| wildtype 25°C vs wildtype 20°C | 629 genes downregulated >2x (q=0.01) | Phenotype Enrichment Analysis | WBPhenotype:0000679 | transgene subcellular localization variant | 76 | 101 | 1.3 | 0.0016 | 0.013 |
| wildtype 25°C vs wildtype 20°C | 629 genes downregulated >2x (q=0.01) | Phenotype Enrichment Analysis | WBPhenotype:0000049 | postembryonic development variant | 140 | 173 | 1.2 | 0.0017 | 0.014 |
| AFD-ablated 20°C vs wildtype 20°C | 2001 genes upregulated >2x (q=0.01) | Phenotype Enrichment Analysis | WBPhenotype:0000603 | muscle system morphology variant | 27 | 59 | 2.2 | 1.9E-09 | 0.00000044 |
| AFD-ablated 20°C vs wildtype 20°C | 2001 genes upregulated >2x (q=0.01) | Phenotype Enrichment Analysis | WBPhenotype:0001763 | uptake by intestinal cell defective | 4.6 | 18 | 4 | 3.3E-08 | 0.000004 |
| AFD-ablated 20°C vs wildtype 20°C | 2001 genes upregulated >2x (q=0.01) | Phenotype Enrichment Analysis | WBPhenotype:0001401 | mitochondria morphology variant | 22 | 49 | 2.2 | 4.8E-08 | 0.000004 |
| AFD-ablated 20°C vs wildtype 20°C | 2001 genes upregulated >2x (q=0.01) | Phenotype Enrichment Analysis | WBPhenotype:0000012 | dauer constitutive | 12 | 32 | 2.7 | 7.6E-08 | 0.0000046 |
| AFD-ablated 20°C vs wildtype 20°C | 2001 genes upregulated >2x (q=0.01) | Phenotype Enrichment Analysis | WBPhenotype:0000644 | paralyzed | 16 | 39 | 2.4 | 0.00000011 | 0.0000053 |
| AFD-ablated 20°C vs wildtype 20°C | 2001 genes upregulated >2x (q=0.01) | Phenotype Enrichment Analysis | WBPhenotype:0001184 | fat content increased | 25 | 51 | 2.1 | 0.00000025 | 0.00001 |
| AFD-ablated 20°C vs wildtype 20°C | 2001 genes upregulated >2x (q=0.01) | Phenotype Enrichment Analysis | WBPhenotype:0001645 | protein degradation variant | 17 | 36 | 2.1 | 0.0000046 | 0.00016 |
| AFD-ablated 20°C vs wildtype 20°C | 2001 genes upregulated >2x (q=0.01) | Phenotype Enrichment Analysis | WBPhenotype:0001010 | clear | 22 | 43 | 2 | 0.0000073 | 0.00022 |
| AFD-ablated 20°C vs wildtype 20°C | 2001 genes upregulated >2x (q=0.01) | Phenotype Enrichment Analysis | WBPhenotype:0000782 | body wall muscle thick filament variant | 14 | 31 | 2.2 | 0.0000082 | 0.00022 |
| AFD-ablated 20°C vs wildtype 20°C | 2001 genes upregulated >2x (q=0.01) | Phenotype Enrichment Analysis | WBPhenotype:0000061 | extended life span | 65 | 98 | 1.5 | 0.000027 | 0.00064 |
| AFD-ablated 20°C vs wildtype 20°C | 2001 genes upregulated >2x (q=0.01) | Phenotype Enrichment Analysis | WBPhenotype:0000229 | small | 57 | 87 | 1.5 | 0.000028 | 0.00064 |
| AFD-ablated 20°C vs wildtype 20°C | 2001 genes upregulated >2x (q=0.01) | Phenotype Enrichment Analysis | WBPhenotype:0000646 | sluggish | 21 | 39 | 1.9 | 0.000044 | 0.00088 |
| AFD-ablated 20°C vs wildtype 20°C | 2001 genes upregulated >2x (q=0.01) | Phenotype Enrichment Analysis | WBPhenotype:0002056 | eating variant | 15 | 30 | 2 | 0.000084 | 0.0015 |
| AFD-ablated 20°C vs wildtype 20°C | 2001 genes upregulated >2x (q=0.01) | Phenotype Enrichment Analysis | WBPhenotype:0002059 | pore forming toxin hypersensitive | 12 | 25 | 2.1 | 0.000088 | 0.0015 |
| AFD-ablated 20°C vs wildtype 20°C | 1832 genes downregulated >2x (q=0.01) | Phenotype Enrichment Analysis | WBPhenotype:0000763 | embryonic cell physiology variant | 82 | 196 | 2.4 | 1.1E-36 | 2.7E-34 |
| AFD-ablated 20°C vs wildtype 20°C | 1832 genes downregulated >2x (q=0.01) | Phenotype Enrichment Analysis | WBPhenotype:0000770 | embryonic cell morphology variant | 34 | 102 | 3 | 7.1E-31 | 8.5E-29 |
| AFD-ablated 20°C vs wildtype 20°C | 1832 genes downregulated >2x (q=0.01) | Phenotype Enrichment Analysis | WBPhenotype:0001114 | cell cycle defective early emb | 19 | 67 | 3.6 | 1.8E-27 | 1.5E-25 |
| AFD-ablated 20°C vs wildtype 20°C | 1832 genes downregulated >2x (q=0.01) | Phenotype Enrichment Analysis | WBPhenotype:0001143 | multiple nuclei early emb | 21 | 72 | 3.4 | 2.5E-27 | 1.5E-25 |
| AFD-ablated 20°C vs wildtype 20°C | 1832 genes downregulated >2x (q=0.01) | Phenotype Enrichment Analysis | WBPhenotype:0001427 | cytoplasmic appearance variant | 27 | 83 | 3.1 | 1E-26 | 4.8E-25 |
| AFD-ablated 20°C vs wildtype 20°C | 1832 genes downregulated >2x (q=0.01) | Phenotype Enrichment Analysis | WBPhenotype:0001079 | cytoplasmic dynamics defective early emb | 13 | 52 | 4 | 3.2E-26 | 1.3E-24 |
| AFD-ablated 20°C vs wildtype 20°C | 1832 genes downregulated >2x (q=0.01) | Phenotype Enrichment Analysis | WBPhenotype:0000759 | spindle defective early emb | 47 | 119 | 2.5 | 6E-26 | 2.1E-24 |
| AFD-ablated 20°C vs wildtype 20°C | 1832 genes downregulated >2x (q=0.01) | Phenotype Enrichment Analysis | WBPhenotype:0002408 | cytokinesis variant | 58 | 137 | 2.4 | 1.3E-25 | 3.8E-24 |
| AFD-ablated 20°C vs wildtype 20°C | 1832 genes downregulated >2x (q=0.01) | Phenotype Enrichment Analysis | WBPhenotype:0002501 | ectopic cleavage furrow early emb | 11 | 45 | 4.3 | 4.4E-25 | 1.2E-23 |
| AFD-ablated 20°C vs wildtype 20°C | 1832 genes downregulated >2x (q=0.01) | Phenotype Enrichment Analysis | WBPhenotype:0002612 | cytoplasmic streaming phenotype | 15 | 57 | 3.7 | 5.9E-25 | 1.4E-23 |
| AFD-ablated 20°C vs wildtype 20°C | 1832 genes downregulated >2x (q=0.01) | Phenotype Enrichment Analysis | WBPhenotype:0000504 | nuclear division variant | 44 | 111 | 2.5 | 2.6E-24 | 5.7E-23 |
| AFD-ablated 20°C vs wildtype 20°C | 1832 genes downregulated >2x (q=0.01) | Phenotype Enrichment Analysis | WBPhenotype:0001588 | microtubule organization biogenesis variant | 55 | 130 | 2.3 | 3.3E-24 | 6.5E-23 |
| AFD-ablated 20°C vs wildtype 20°C | 1832 genes downregulated >2x (q=0.01) | Phenotype Enrichment Analysis | WBPhenotype:0000774 | gametogenesis variant | 260 | 402 | 1.6 | 1.7E-21 | 3.1E-20 |
| AFD-ablated 20°C vs wildtype 20°C | 1832 genes downregulated >2x (q=0.01) | Phenotype Enrichment Analysis | WBPhenotype:0001499 | meiotic chromosome segregation variant | 41 | 98 | 2.4 | 6.9E-20 | 1.2E-18 |
| AFD-ablated 20°C vs wildtype 20°C | 1832 genes downregulated >2x (q=0.01) | Phenotype Enrichment Analysis | WBPhenotype:0001026 | nuclear morphology variation early emb | 12 | 44 | 3.6 | 2.4E-19 | 3.8E-18 |
| AFD-ablated 20°C vs wildtype 20°C | 1832 genes downregulated >2x (q=0.01) | Phenotype Enrichment Analysis | WBPhenotype:0001882 | aneuploidy | 37 | 88 | 2.4 | 7.5E-18 | 1.1E-16 |
| AFD-ablated 20°C vs wildtype 20°C | 1832 genes downregulated >2x (q=0.01) | Phenotype Enrichment Analysis | WBPhenotype:0000069 | progeny variant | 160 | 259 | 1.6 | 1.1E-17 | 1.5E-16 |
| AFD-ablated 20°C vs wildtype 20°C | 1832 genes downregulated >2x (q=0.01) | Phenotype Enrichment Analysis | WBPhenotype:0001374 | pronuclear nuclear appearance variant emb | 35 | 83 | 2.4 | 5.1E-17 | 6.8E-16 |
| AFD-ablated 20°C vs wildtype 20°C | 1832 genes downregulated >2x (q=0.01) | Phenotype Enrichment Analysis | WBPhenotype:0001116 | absolute cell cycle timing defective early emb | 9.8 | 35 | 3.6 | 1.4E-15 | 1.8E-14 |
| AFD-ablated 20°C vs wildtype 20°C | 1832 genes downregulated >2x (q=0.01) | Phenotype Enrichment Analysis | WBPhenotype:0000762 | spindle position defective early emb | 12 | 40 | 3.3 | 1.8E-15 | 2.1E-14 |
| AFD-ablated 20°C vs wildtype 20°C | 1832 genes downregulated >2x (q=0.01) | Phenotype Enrichment Analysis | WBPhenotype:0000697 | protruding vulva | 160 | 244 | 1.5 | 2.8E-13 | 3.2E-12 |
| AFD-ablated 20°C vs wildtype 20°C | 1832 genes downregulated >2x (q=0.01) | Phenotype Enrichment Analysis | WBPhenotype:0001335 | hermaphrodite reproductive system morphology v | 180 | 264 | 1.5 | 1.6E-12 | 1.7E-11 |
| AFD-ablated 20°C vs wildtype 20°C | 1832 genes downregulated >2x (q=0.01) | Phenotype Enrichment Analysis | WBPhenotype:0001681 | spindle orientation variant | 20 | 50 | 2.5 | 5.3E-12 | 5.5E-11 |
| AFD-ablated 20°C vs wildtype 20°C | 1832 genes downregulated >2x (q=0.01) | Phenotype Enrichment Analysis | WBPhenotype:0001082 | large cytoplasmic granules early emb | 10 | 31 | 3.1 | 1.8E-11 | 1.8E-10 |
| AFD-ablated 20°C vs wildtype 20°C | 1832 genes downregulated >2x (q=0.01) | Phenotype Enrichment Analysis | WBPhenotype:0000749 | embryonic development variant | 700 | 857 | 1.2 | 3E-11 | 2.9E-10 |
| AFD-ablated 20°C vs wildtype 20°C | 1832 genes downregulated >2x (q=0.01) | Phenotype Enrichment Analysis | WBPhenotype:0001408 | pattern protein expression variant | 64 | 113 | 1.8 | 1.1E-10 | 1E-09 |
| AFD-ablated 20°C vs wildtype 20°C | 1832 genes downregulated >2x (q=0.01) | Phenotype Enrichment Analysis | WBPhenotype:0000040 | one cell arrest early emb | 12 | 32 | 2.8 | 6.7E-10 | 5.9E-09 |
| AFD-ablated 20°C vs wildtype 20°C | 1832 genes downregulated >2x (q=0.01) | Phenotype Enrichment Analysis | WBPhenotype:0001403 | antibody subcellular localization variant | 13 | 34 | 2.6 | 2.4E-09 | 2.1E-08 |
| AFD-ablated 20°C vs wildtype 20°C | 1832 genes downregulated >2x (q=0.01) | Phenotype Enrichment Analysis | WBPhenotype:0000218 | vulval cell induction increased | 46 | 82 | 1.8 | 8.6E-09 | 7.1E-08 |
| AFD-ablated 20°C vs wildtype 20°C | 1832 genes downregulated >2x (q=0.01) | Phenotype Enrichment Analysis | WBPhenotype:0000777 | polar body extrusion defective early emb | 9.8 | 27 | 2.8 | 1.3E-08 | 0.0000001 |
| AFD-ablated 20°C vs wildtype 20°C | 1832 genes downregulated >2x (q=0.01) | Phenotype Enrichment Analysis | WBPhenotype:0001276 | ectopic expression transgene | 28 | 55 | 1.9 | 8.2E-08 | 0.00000063 |
| AFD-ablated 20°C vs wildtype 20°C | 1832 genes downregulated >2x (q=0.01) | Phenotype Enrichment Analysis | WBPhenotype:0000220 | vulva cell fate specification variant | 59 | 97 | 1.6 | 8.8E-08 | 0.00000066 |
| AFD-ablated 20°C vs wildtype 20°C | 1832 genes downregulated >2x (q=0.01) | Phenotype Enrichment Analysis | WBPhenotype:0001378 | mitotic chromosome segregation variant | 13 | 31 | 2.4 | 0.00000012 | 0.00000083 |
| AFD-ablated 20°C vs wildtype 20°C | 1832 genes downregulated >2x (q=0.01) | Phenotype Enrichment Analysis | WBPhenotype:0001301 | P granule abnormal | 11 | 28 | 2.5 | 0.00000012 | 0.00000083 |
| AFD-ablated 20°C vs wildtype 20°C | 1832 genes downregulated >2x (q=0.01) | Phenotype Enrichment Analysis | WBPhenotype:0001041 | meiosis defective early emb | 16 | 36 | 2.2 | 0.00000028 | 0.0000019 |
| AFD-ablated 20°C vs wildtype 20°C | 1832 genes downregulated >2x (q=0.01) | Phenotype Enrichment Analysis | WBPhenotype:0000425 | antibody staining reduced | 18 | 39 | 2.1 | 0.00000029 | 0.0000019 |
| AFD-ablated 20°C vs wildtype 20°C | 1832 genes downregulated >2x (q=0.01) | Phenotype Enrichment Analysis | WBPhenotype:0001020 | embryonic lethal late emb | 15 | 33 | 2.3 | 0.00000036 | 0.0000023 |
| AFD-ablated 20°C vs wildtype 20°C | 1832 genes downregulated >2x (q=0.01) | Phenotype Enrichment Analysis | WBPhenotype:0000038 | exploded through vulva | 81 | 120 | 1.5 | 0.0000012 | 0.0000077 |
| AFD-ablated 20°C vs wildtype 20°C | 1832 genes downregulated >2x (q=0.01) | Phenotype Enrichment Analysis | WBPhenotype:0001208 | RNAi resistant | 28 | 51 | 1.9 | 0.0000012 | 0.0000077 |
| AFD-ablated 20°C vs wildtype 20°C | 1832 genes downregulated >2x (q=0.01) | Phenotype Enrichment Analysis | WBPhenotype:0001595 | somatic transgene silencing variant | 20 | 40 | 2 | 0.0000017 | 0.00001 |
| AFD-ablated 20°C vs wildtype 20°C | 1832 genes downregulated >2x (q=0.01) | Phenotype Enrichment Analysis | WBPhenotype:0001036 | sterile F1 | 14 | 30 | 2.2 | 0.0000022 | 0.000013 |
| AFD-ablated 20°C vs wildtype 20°C | 1832 genes downregulated >2x (q=0.01) | Phenotype Enrichment Analysis | WBPhenotype:0001752 | vesicle trafficking variant | 160 | 207 | 1.3 | 0.0000036 | 0.000021 |
| AFD-ablated 20°C vs wildtype 20°C | 1832 genes downregulated >2x (q=0.01) | Phenotype Enrichment Analysis | WBPhenotype:0000889 | sexually dimorphic physiology variant | 150 | 198 | 1.3 | 0.0000042 | 0.000023 |
| AFD-ablated 20°C vs wildtype 20°C | 1832 genes downregulated >2x (q=0.01) | Phenotype Enrichment Analysis | WBPhenotype:0000772 | sister chromatid segregation defective early emb | 18 | 35 | 2 | 0.0000062 | 0.000034 |
| AFD-ablated 20°C vs wildtype 20°C | 1832 genes downregulated >2x (q=0.01) | Phenotype Enrichment Analysis | WBPhenotype:0000691 | gonad development variant | 99 | 138 | 1.4 | 0.000013 | 0.000069 |
| AFD-ablated 20°C vs wildtype 20°C | 1832 genes downregulated >2x (q=0.01) | Phenotype Enrichment Analysis | WBPhenotype:0000351 | failure to hatch | 12 | 25 | 2.1 | 0.000022 | 0.00012 |
| AFD-ablated 20°C vs wildtype 20°C | 1832 genes downregulated >2x (q=0.01) | Phenotype Enrichment Analysis | WBPhenotype:0000701 | epithelial development variant | 17 | 33 | 1.9 | 0.000035 | 0.00018 |
| AFD-ablated 20°C vs wildtype 20°C | 1832 genes downregulated >2x (q=0.01) | Phenotype Enrichment Analysis | WBPhenotype:0002075 | eggshell formation variant | 30 | 50 | 1.7 | 0.000058 | 0.00029 |
| AFD-ablated 20°C vs wildtype 20°C | 1832 genes downregulated >2x (q=0.01) | Phenotype Enrichment Analysis | WBPhenotype:0000111 | pattern of gene expression variant | 370 | 435 | 1.2 | 0.000063 | 0.00031 |
| AFD-ablated 20°C vs wildtype 20°C | 1832 genes downregulated >2x (q=0.01) | Phenotype Enrichment Analysis | WBPhenotype:0000172 | cell proliferation increased | 20 | 36 | 1.8 | 0.000076 | 0.00036 |
| AFD-ablated 20°C vs wildtype 20°C | 1832 genes downregulated >2x (q=0.01) | Phenotype Enrichment Analysis | WBPhenotype:0001038 | tumorous germline | 17 | 32 | 1.8 | 0.000089 | 0.00042 |
| AFD-ablated 20°C vs wildtype 20°C | 1832 genes downregulated >2x (q=0.01) | Phenotype Enrichment Analysis | WBPhenotype:0000673 | brood size variant | 250 | 308 | 1.2 | 0.0001 | 0.00048 |
| AFD-ablated 20°C vs wildtype 20°C | 1832 genes downregulated >2x (q=0.01) | Phenotype Enrichment Analysis | WBPhenotype:0002369 | sister cell division timing asynchrony reduced | 15 | 29 | 1.9 | 0.00011 | 0.0005 |
| AFD-ablated 20°C vs wildtype 20°C | 1832 genes downregulated >2x (q=0.01) | Phenotype Enrichment Analysis | WBPhenotype:0000228 | spontaneous mutation rate increased | 18 | 31 | 1.8 | 0.00028 | 0.0012 |
| AFD-ablated 20°C vs wildtype 20°C | 1832 genes downregulated >2x (q=0.01) | Phenotype Enrichment Analysis | WBPhenotype:0001022 | hermaphrodite sexual development variant | 14 | 26 | 1.8 | 0.0003 | 0.0013 |
| AFD-ablated 20°C vs wildtype 20°C | 1832 genes downregulated >2x (q=0.01) | Phenotype Enrichment Analysis | WBPhenotype:0001180 | accumulated germline cell corpses | 48 | 69 | 1.4 | 0.00037 | 0.0016 |
| AFD-ablated 20°C vs wildtype 20°C | 1832 genes downregulated >2x (q=0.01) | Phenotype Enrichment Analysis | WBPhenotype:0000683 | masculinization of germline | 10 | 20 | 2 | 0.00043 | 0.0018 |
| AFD-ablated 20°C vs wildtype 20°C | 1832 genes downregulated >2x (q=0.01) | Phenotype Enrichment Analysis | WBPhenotype:0000729 | cell death variant | 160 | 198 | 1.2 | 0.00048 | 0.002 |
| AFD-ablated 20°C vs wildtype 20°C | 1832 genes downregulated >2x (q=0.01) | Phenotype Enrichment Analysis | WBPhenotype:0000274 | dead eggs laid | 18 | 30 | 1.7 | 0.00079 | 0.0032 |
| AFD-ablated 20°C vs wildtype 20°C | 1832 genes downregulated >2x (q=0.01) | Phenotype Enrichment Analysis | WBPhenotype:0001977 | germ cell partition morphology variant | 16 | 28 | 1.7 | 0.0009 | 0.0036 |
| AFD-ablated 20°C vs wildtype 20°C | 1832 genes downregulated >2x (q=0.01) | Phenotype Enrichment Analysis | WBPhenotype:0002188 | male somatic gonad development variant | 26 | 40 | 1.6 | 0.00092 | 0.0036 |
| AFD-ablated 20°C vs wildtype 20°C | 1832 genes downregulated >2x (q=0.01) | Phenotype Enrichment Analysis | WBPhenotype:0001925 | oocytes disorganized | 14 | 25 | 1.8 | 0.00096 | 0.0037 |
| AFD-ablated 20°C vs wildtype 20°C | 1832 genes downregulated >2x (q=0.01) | Phenotype Enrichment Analysis | WBPhenotype:0001384 | fertility reduced | 710 | 777 | 1.1 | 0.0013 | 0.005 |
| AFD-ablated 20°C vs wildtype 20°C | 1832 genes downregulated >2x (q=0.01) | Phenotype Enrichment Analysis | WBPhenotype:0001235 | cell division polarity variant | 15 | 24 | 1.6 | 0.0034 | 0.013 |
| AFD-ablated 20°C vs wildtype 20°C | 1832 genes downregulated >2x (q=0.01) | Phenotype Enrichment Analysis | WBPhenotype:0001361 | chromosome condensation variant | 42 | 58 | 1.4 | 0.0039 | 0.014 |
| AFD-ablated 20°C vs wildtype 20°C | 1832 genes downregulated >2x (q=0.01) | Phenotype Enrichment Analysis | WBPhenotype:0001897 | light response variant | 17 | 27 | 1.6 | 0.0042 | 0.015 |
| AFD-ablated 20°C vs wildtype 20°C | 1832 genes downregulated >2x (q=0.01) | Phenotype Enrichment Analysis | WBPhenotype:0000365 | embryo osmotic integrity defective early emb | 26 | 38 | 1.5 | 0.005 | 0.018 |

### Supplementary Table 5. Statistical analysis for Figures 4 and S6.

| Set | Genotype | Temperature: egg → day 2 adult | Temperature: survival assay | Mean survival ± SEM (days) | Median survival (days) | 75th percentile (days) | N dead / initial N | Group | % Mean survival change vs. group a | *P* value (log-rank) vs. group a | *P* value (log-rank) vs. group b | *P* value (log-rank) vs. group c | Figure |
|---|---|---|---|---|---|---|---|---|---|---|---|---|---|
| Survival in 6 mM tBuOOH, *E. coli* OP50 | | | | | | | | | | | | | |
| | wild type | 20°C | 20°C | 1.94 ± 0.03 | 1.90 | 2.11 | 102 / 102 | a | | | | | 4F |
| | *ins-39(tm6467) X* | 20°C | 20°C | 2.51 ± 0.05 | 2.58 | 2.86 | 133 / 158 | b | 29% | < 0.0001 | | | |
| | wild type | 25°C | 20°C | 3.29 ± 0.08 | 3.10 | 3.89 | 129 / 129 | c | 69% | < 0.0001 | < 0.0001 | | |
| | *ins-39(tm6467) X* | 25°C | 20°C | 3.44 ± 0.08 | 3.42 | 4.05 | 124 / 124 | d | 77% | < 0.0001 | < 0.0001 | > 0.05 | |
| | AFD ablated | 20°C | 20°C | 4.86 ± 0.11 | 4.81 | 5.62 | 133 / 133 | a | | | | | S6B |
| | AFD ablated; *tph-1(n4622) II* | 20°C | 20°C | 4.69 ± 0.14 | 4.62 | 5.62 | 133 / 133 | b | -4% | > 0.05 | | | |

**Supplementary Table 6. Regulation of DAF-16 target genes among genes up- and down-regulated significantly (q value < 0.01) by growth at 25°C and by AFD ablation at 20°C.**

| | | DAF-16 targets (Kumar et al., 2015) | | Not DAF-16 targets (Kumar et al., 2015) | |
|---|---|---|---|---|---|
| gene expression effect [wild type 25°C / wild type 20°C] | gene expression effect [AFD(-) 20°C / wild type 20°C] | genes (% total) | *P* value (cell chi-square test) | genes (% total) | P value (cell chi-square test) |
| up at 25°C | up in AFD(-) | 8 (3.0%) | < 0.0001 | 257 (97.0%) | > 0.05 |
| up at 25°C | no effect | 2 (1.4%) | 0.037 | 137 (98.6%) | > 0.05 |
| up at 25°C | down in AFD(-) | 0 (0.0%) | > 0.05 | 10 (100.0%) | > 0.05 |
| no effect | up in AFD(-) | 11 (0.7%) | 0.016 | 1464 (99.3%) | > 0.05 |
| no effect | no effect | 6 (0.2%) | > 0.05 | 2539 (99.8%) | > 0.05 |
| no effect | down in AFD(-) | 1 (0.1%) | > 0.05 | 1194 (99.9%) | > 0.05 |
| down at 25°C | up in AFD(-) | 0 (0.0%) | > 0.05 | 475 (100.0%) | > 0.05 |
| down at 25°C | no effect | 1 (0.1%) | > 0.05 | 925 (99.9%) | > 0.05 |
| down at 25°C | down in AFD(-) | 0 (0.0%) | > 0.05 | 872 (100.0%) | > 0.05 |

### Supplementary Table 7. Statistical analysis for Figures 5 and S7.

| | | Temperature | | | | | | | | | | | |
|---|---|---|---|---|---|---|---|---|---|---|---|---|---|
| **Set** | **Genotype** | **egg → day 2 adult** | **survival assay** | **Mean survival ± SEM (days)** | **Median survival (days)** | **75th percentile (days)** | **N dead / initial N** | **Group** | **% Mean survival change vs. group a** | ***P* value (log-rank) vs. group a** | ***P* value (log-rank) vs. group b** | ***P* value (log-rank) vs. group c** | **Figure** |
| Survival in 6 mM tBuOOH, *E. coli* OP50 | | | | | | | | | | | | | |
| | wild type | 20°C | 20°C | 1.94 ± 0.04 | 1.82 | 2.21 | 150 / 152 | a | | | | | 5C |
| | *daf-16(mu86) I* | 20°C | 20°C | 1.54 ± 0.04 | 1.55 | 1.69 | 84 / 87 | b | -21% | < 0.0001 | | | |
| | wild type | 25°C | 20°C | 3.04 ± 0.07 | 2.95 | 3.54 | 140 / 149 | c | 57% | < 0.0001 | < 0.0001 | | |
| | *daf-16(mu86) I* | 25°C | 20°C | 1.97 ± 0.04 | 1.95 | 2.31 | 136 / 150 | d | 2% | > 0.05 | < 0.0001 | < 0.0001 | |
| | wild type | 20°C | 20°C | 2.14 ± 0.04 | 2.08 | 2.42 | 110 / 110 | a | | | | | 5D |
| | AFD ablated | 20°C | 20°C | 5.52 ± 0.11 | 5.51 | 6.51 | 127 / 138 | b | 158% | < 0.0001 | | | |
| | *daf-16(mu86) I* | 20°C | 20°C | 1.76 ± 0.04 | 1.75 | 2.03 | 156 / 192 | c | -18% | < 0.0001 | < 0.0001 | | |
| | AFD ablated; *daf-16(mu86) I* | 20°C | 20°C | 2.30 ± 0.09 | 2.34 | 2.75 | 68 / 68 | d | 7% | 0.033 | < 0.0001 | < 0.0001 | |
| | AFD ablated; *daf-16(mu86) I* | 20°C | 20°C | 2.24 ± 0.04 | 2.25 | 2.55 | 161 / 176 | a | | | | | 5E |
| | AFD ablated; *daf-16(mu86) I; qyIs292[Pges-1::GFP::daf-16]* | 20°C | 20°C | 3.24 ± 0.14 | 3.52 | 4.10 | 52 / 70 | b | 44% | < 0.0001 | | | |
| | AFD ablated | 20°C | 20°C | 3.93 ± 0.08 | 3.98 | 4.57 | 108 / 123 | c | 75% | < 0.0001 | < 0.0001 | 0.0003 | |
| | *daf-16(mu86) I* | 20°C | 20°C | 1.77 ± 0.03 | 1.75 | 2.00 | 142 / 155 | a | | | | | 5F |
| | *daf-16(mu86) I; qyIs292[Pges-1::GFP::daf-16]* | 20°C | 20°C | 2.03 ± 0.05 | 2.05 | 2.36 | 101 / 116 | b | 15% | < 0.0001 | | | |
| | *daf-16(mu86) I* | 25°C | 20°C | 1.99 ± 0.04 | 1.94 | 2.29 | 155 / 155 | c | 12% | < 0.0001 | > 0.05 | | |
| | *daf-16(mu86) I; qyIs292[Pges-1::GFP::daf-16]* | 25°C | 20°C | 2.77 ± 0.06 | 2.70 | 3.27 | 136 / 136 | d | 57% | < 0.0001 | < 0.0001 | < 0.0001 | |
| | AFD ablated | 20°C | 20°C | 5.15 ± 0.23 | 5.09 | 6.32 | 65 / 65 | a | | | | | S7C |
| | AFD ablated; *daf-3(mgDf90) X* | 20°C | 20°C | 5.45 ± 0.23 | 5.54 | 6.68 | 66 / 66 | b | 6% | > 0.05 | | | |

### Supplementary Table 8. Statistical analysis for Figure 6.

| Set | Genotype | Temperature: egg → day 2 adult | Temperature: survival assay | Mean survival ± SEM (days) | Median survival (days) | 75th percentile (days) | N dead / initial N | Group | % Mean survival change vs. group a | *P* value (log-rank) vs. group a | *P* value (log-rank) vs. group b | *P* value (log-rank) vs. group c | Figure |
|---|---|---|---|---|---|---|---|---|---|---|---|---|---|
| Survival in 6 mM tBuOOH, *E. coli* OP50 | | | | | | | | | | | | | |
| | wild type + empty vector | 20°C | 20°C | 2.77 ± 0.05 | 2.72 | 2.98 | 129 / 141 | a | | | | | 6B |
| | AFD ablation + empty vector | 20°C | 20°C | 6.70 ± 0.19 | 6.46 | 8.12 | 109 / 115 | b | 142% | < 0.0001 | | | |
| | wild type + *skn-1(RNAi)* | 25°C | 20°C | 2.02 ± 0.06 | 1.95 | 2.27 | 132 / 132 | c | -27% | < 0.0001 | < 0.0001 | | |
| | AFD ablation + *skn-1(RNAi)* | 25°C | 20°C | 2.80 ± 0.10 | 2.78 | 3.24 | 77 / 77 | d | 1% | > 0.05 | < 0.0001 | < 0.0001 | |
| | *daf-16(mu86) I* + empty vector | 20°C | 20°C | 1.44 ± 0.04 | 1.35 | 1.61 | 142 / 161 | a | | | | | 6C |
| | AFD ablation; *daf-16(mu86) I* + empty vector | 20°C | 20°C | 2.92 ± 0.06 | 3.06 | 3.31 | 151 / 166 | b | 103% | < 0.0001 | | | |
| | *daf-16(mu86) I* + *skn-1(RNAi)* | 25°C | 20°C | 0.67 ± 0.03 | 0.64 | 0.74 | 69 / 77 | c | -54% | < 0.0001 | < 0.0001 | | |
| | AFD ablation; *daf-16(mu86) I* + *skn-1(RNAi)* | 25°C | 20°C | 1.04 ± 0.02 | 1.02 | 1.15 | 155 / 172 | d | -28% | < 0.0001 | < 0.0001 | < 0.0001 | |

#### Supplementary Table 9. Bacterial strains.

| Strain | Genotype | Source | Reference |
|---|---|---|---|
| OP50 | *E. coli* B, uracil auxotroph | CGC | Brenner, 1974 |
| HT115 | *E. coli* $F^-$ *mcrA, mcrB, IN(rrnD-rrnE)1, rnc14::Tn10(DE3 lysogen: lavUV5 promoter -T7 polymerase).* Tetracycline resistant | CGC | Kamath et al., 2001 |
| JI377 | *E. coli* K12 $F^-$ *ahpCF katG katE* | James Imlay | Seaver and Imlay, 2001 |
| E007 | *Enterococcus faecium* wild type | Danielle Garsin | Chavez et al, 2007 |

#### Supplementary Table 10. Nematode strains.

| Strain | Species | Genotype | Source | Reference |
|---|---|---|---|---|
| QZ0 | *Caenorhabditis elegans* | N2 wild type | Joy Alcedo | Schiffer et. al, 2020 |
| PR678 | *Caenorhabditis elegans* | *tax-4(p678) III* | CGC | |
| SAY141 | *Caenorhabditis elegans* | *pgIs2[gcy-8p::TU#813 + gcy-8p::TU#814 + unc-122p::GFP + gcy-8p::mCherry + gcy-8p::GFP + ttx-3p::GFP] 6x outcrossed* | CGC | Glauser et al., 2011; Schiffer et. al, 2020 |
| MT14984 | *Caenorhabditis elegans* | *tph-1(n4622) II* | CGC | Shivers et al., 2009 |
| SAY182 | *Caenorhabditis elegans* | *tph-1(n4622) II; pgIs2[gcy-8p::TU#813 + gcy-8p::TU#814 + unc-122p::GFP + gcy-8p::mCherry + gcy-8p::GFP + ttx-3p::GFP]* | This work | |
| PY12304 | *Caenorhabditis elegans* | *ins-39(oy167[ins-39::SL2::GFP]) X* | This work | |
| PY12305 | *Caenorhabditis elegans* | *tax-4(p678) III; ins-39(oy167[ins-39::SL2::GFP]) X* | This work | |
| ZC2751 | *Caenorhabditis elegans* | *ins-39(tm6467) X* | Yun Zhang | Wu et. al, 2019 |
| QZ60 | *Caenorhabditis elegans* | *daf-16(mu86) I* | This work | |
| SAY144 | *Caenorhabditis elegans* | *daf-16(mu86); pgIs2[gcy-8p::TU#813 + gcy-8p::TU#814 + unc-122p::GFP + gcy-8p::mCherry + gcy-8p::GFP + ttx-3p::GFP]* | This work | |
| SAY203 | *Caenorhabditis elegans* | *daf-16(mu86) I; pgIs2[gcy-8p::TU#813 + gcy-8p::TU#814 + unc-122p::GFP + gcy-8p::mCherry + gcy-8p::GFP + ttx-3p::GFP]; qyIs292[Pges-1::GFP::daf-16]* | This work | |
| NK1232 | *Caenorhabditis elegans* | *daf-16(mu86); qyIs292[Pges-1::GFP::daf-16]* | Ryan Baugh | Kaplan et al., 2015 |
| SAY190 | *Caenorhabditis elegans* | *pgIs2[gcy-8p::TU#813 + gcy-8p::TU#814 + unc-122p::GFP + gcy-8p::mCherry + gcy-8p::GFP + ttx-3p::GFP]; daf-3(mgDf90)* | This work | |
| GR1311 | *Caenorhabditis elegans* | *daf-3(mgDf90) X* | CGC | |

**Supplementary Table 11. PCR genotyping primers and phenotypes used for strain construction.**

| Allele | Phenotype | Primers |
|---|---|---|
| *tax-4(p678)* | sequencing | GAGACGATCATCCACTTGGTAAG<br>AAATCTTCCGGATCCAACTCTAC |
| *pgIs2[gcy-8p::TU#813 + gcy-8p::TU#814 + unc-122p::GFP + gcy-8p::mCherry + gcy-8p::GFP + ttx-3p::GFP]* | fluorescent marker | - |
| *tph-1(n4622)* | - | AGATTGTGTGGCAGGCGGC<br>TGATGTCTGAGCGGAGAGAGTT |
| *ins-39(oy167[ins-39::SL2::GFP]* | fluorescent marker | - |
| *daf-16(mu86)* | - | AGAACACCATGGGGGCACTGGAT<br>GGCGGGAATGAAGCAAGAGCCAA<br>TGACGCTCACCTTGAAAAGGTCAAT<br>GGAACCGATTCGCCAACCCATGA |
| *qyIs292[Pges-1::GFP::daf-16]* | fluorescent marker | - |
| *daf-3(mgDf90)* | - | AAACGCGTCATGTGGACCAC<br>ACCCTCATGCCTACTGTCAG |
